# The mesoSPIM initiative: open-source light-sheet mesoscopes for imaging in cleared tissue

**DOI:** 10.1101/577122

**Authors:** Fabian F. Voigt, Daniel Kirschenbaum, Evgenia Platonova, Stéphane Pagès, Robert A. A. Campbell, Rahel Kästli, Martina Schaettin, Ladan Egolf, Alexander van der Bourg, Philipp Bethge, Karen Haenraets, Noémie Frézel, Thomas Topilko, Paola Perin, Daniel Hillier, Sven Hildebrand, Anna Schueth, Alard Roebroeck, Botond Roska, Esther Stoeckli, Roberto Pizzala, Nicolas Renier, Hanns Ulrich Zeilhofer, Theofanis Karayannis, Urs Ziegler, Laura Batti, Anthony Holtmaat, Christian Lüscher, Adriano Aguzzi, Fritjof Helmchen

## Abstract

Over the course of the past decade, tissue clearing methods have reached a high level of sophistication with a wide variety of approaches now available^1^. To image large cleared samples, light-sheet microscopes have proven to be ideal due to their excellent optical sectioning capability in transparent tissue^2^. Such instruments have recently seen extensive technological and commercial development. However, despite this progress, the community is lacking instruments capable of exploring large samples with near-isotropic resolution within minutes. Here, we introduce the mesoscale selective plane-illumination microscopy (mesoSPIM) initiative, an open-hardware project that provides researchers with instructions and software to easily build and operate light-sheet microscopes for centimeter-sized cleared samples (http://www.mesospim.org).

A wide range of commercial light-sheet microscopes have become available since the invention of SPIM^3^, most of them optimized towards time-lapse imaging in transparent developing embryos or – if designed for cleared tissue – tailored for only a narrow selection of clearing techniques and immersion media. To overcome these limitations, we set out to design a modular light-sheet mesoscope (Fig. 1a) that combines simple and versatile sample handling with large fields-of-view (FOV) of 2-20 mm. Large-FOV light-sheet microscopes typically suffer from non-uniform axial resolution due to the varying thickness of the light sheet (Fig. 1b). To address this issue, we use a tunable lens to shift the excitation beam waist through the sample in synchrony with the rolling shutter of the camera, a method called ‘axially scanned light-sheet microscopy’ (ASLM)^4^ (Fig. 1b). For whole mouse brains (≈ 1 cm^3^), typical datasets are isotropic (6.5 µm sampling), small (12-16 GB), and they are recorded quickly (7-8 minutes) with minimal shadow artifacts. Together with standardized quick-exchange holders, these features allow for the fast screening of samples. With a travel range of 44×44×100 mm, very large samples such as a whole mouse central nervous system can be imaged. After acquiring overview datasets, users can zoom in and record multidimensional data at higher resolution by mosaic acquisitions, for example revealing cellular distribution and long-range axonal projections of Purkinje cells in the mouse cerebellum (Fig. 1c-e) or fine neurites in the developing nervous system of a chick embryo (Fig. 1f). The instrument has been tested in combination with all common clearing methods ranging from CLARITY^5^, CUBIC^6^ to organic solvent approaches such as iDISCO^7^ and BABB^2^. Due to the modular design of the mesoSPIM, switching between different imaging media can be done in less than a minute. The mesoSPIM setup, performance characterization, and example applications are presented with detail in Supplementary Notes including Supplementary Figures 1-30. Inspired by the openSPIM^8^ and openSPIN^9^ projects, documentation for the mesoSPIM hardware and software (written in Python) is freely available (https://github.com/mesoSPIM). Currently, there are 5 mesoSPIM setups in operation across Europe. The mesoSPIM is the ideal instrument to quickly bridge scales from the µm- to the cm-level, and serves as an excellent tool for detailed three-dimensional anatomical investigations in neuroscience and developmental biology.

**Figure 1:**
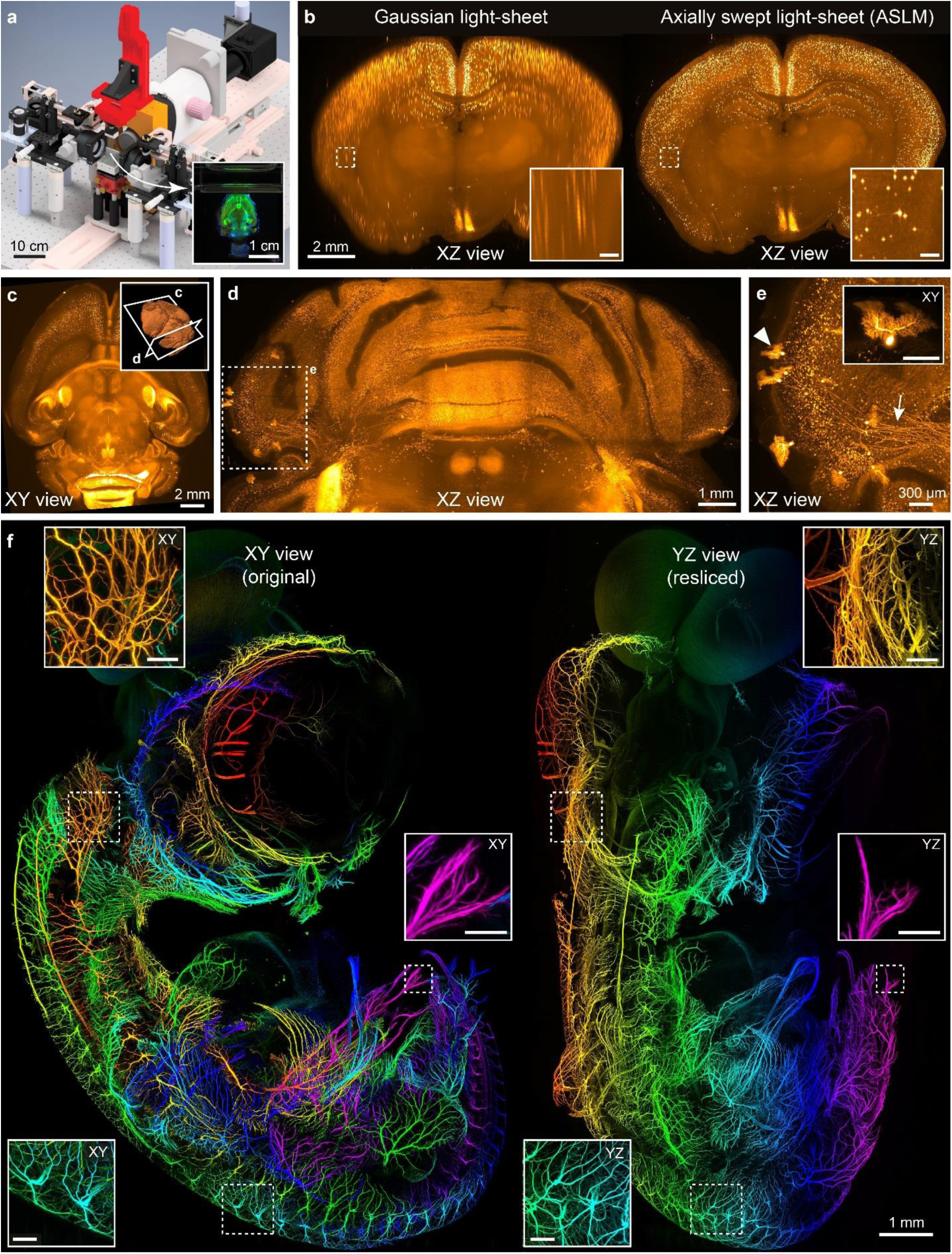
Example demonstrations of the mesoSPIM light-sheet mesoscope. **a**) Overview of the mesoSPIM instrument. Inset: Photograph of a Thy1-YFP mouse brain during image acquisition. **b**) Comparison of axial image quality achieved in a CLARITY-cleared VIP-tdTomato mouse brain for scanning with a Gaussian beam (left) and for the axially swept light-sheet mode, ASLM (right). Images are maximum intensity projections over 250-µm range. **c**) Overview image of a CLARITY-cleared TPH2Cre-tdTomato mouse brain. Inset shows 3D orientation of views in c and d. **d**) XZ maximum intensity projection of a high-resolution dataset (4× zoom, range: 500 µm) taken in the cerebellum of the sample in c. **e**) Volume rendering of sparsely labeled Purkinje cells and their axonal projections (arrow). Inset: individual Purkinje cell (arrow head, scale bar: 200 µm). **f**) Depth-coded original XY (left) and resliced YZ (right) projections of a dataset taken from a 7-day old chicken embryo (neurofilament labeling) cleared using BABB. Throughout the dataset (acquired at 1.6×1.6×2 µm^3^ sampling), neurites are visible in great detail. Because of the ASLM mode this same holds true for the original (transverse) and the resliced (axial) direction. The assignment of color to Z-position is similar for both the XY and YZ view. Scale bars of all insets: 200 µm.

## Code availability

The mesoSPIM software and documentation are available on Github (https://github.com/mesoSPIM). mesoSPIM-control is licensed under the GNU General Public License v3.0 (GPL v3).

## Acknowledgements

This work was supported by grants from the Swiss National Science Foundation (<u>310030B_170269</u>, F.H.; <u>31003A-153448, 31003A_173125, CRSII3_154453, and NCCR Synapsy 51NF40-158776</u>, A.H.), the European Research Council (<u>ERC Advanced Grant BRAINCOMPATH, project 670757</u>; F.H.), ERC Starting Grant (<u>MULTICONNECT, project 639938; AR</u>), the Dutch science foundation (<u>NWO VIDI Grant, project 14637</u>; AR), and a gift from a private foundation with public interest through the International Foundation for Research in Paraplegia (A.H. and S.P.) In addition, we would like to thank Dubravka Göckeritz-Dujmovic and Sandrine Bichet for help with sample preparation and Martin Wieckhorst for help with custom electronics.

## Author information - Contributions

F.F.V.: planned the project, designed the microscope, wrote control software and documentation, set up several mesoSPIM instruments, coordinated the mesoSPIM initiative, imaged samples, analyzed data and wrote the manuscript.

R.K.,M.S, L.E., A. v.d. B, K. H., N.F., T.T., N.R., H-U. Z., T.K., P.P., R.P., D.H., B.R, S.H., A.S., A.R.: Prepared samples for imaging.

S.P., E. P., D.K., R.A.A.C, F.M., L.B. A.H., C.L., A.A.: Set up mesoSPIM instruments.

F.H. planned the project and wrote the manuscript.

## Competing interests

None to disclose.

