## Supplementary Notes and Figures for "The mesoSPIM initiative: open-source light-sheet mesoscopes for imaging in cleared tissue"

**Supplementary Note: The mesoSPIM initative – overview and mesoSPIM features**

**Overview**

The 'mesoscale selective plane illumination microscopy' (mesoSPIM) initiative aims to provide the imaging community with open-source light-sheet microscopes for large cleared samples. On the one hand, it is aimed at neuroscientists and developmental biologists seeking high-quality anatomical data from cleared samples, on the other hand, it strives to provide instrumentation developers with an imaging platform that can be tailored towards specific needs – i.e. to accommodate uncommonly large samples or different illumination schemes. To achieve these goals, we have designed the mesoSPIM to be compatible with all major clearing techniques, ranging from hydrogel-based approaches such as CLARITY to organic-solvent based clearing techniques such as BABB and iDISCO (see **Supplementary Note on compatibility with different clearing techniques**). Both the immersion cuvettes and sample holders are highly modular to allow quick sample exchange and customization (see **Supplementary Notes on the mesoSPIM setup)**. In addition, we noted that in a light-sheet microscope, data acquisition, visualization and analysis are greatly facilitated if the microscope generates raw data at high and uniform axial resolution across the FOV. The mesoSPIM achieves this by incorporating an axially scanned light-sheet microscopy (ASLM) mode (see **Supplementary Note on ASLM**). ASLM facilitates data acquisition as every generated raw image has uniform axial resolution which mean that fewer tiles are necessary to cover a sample compared to a microscope with varying light-sheet thickness (where only a narrow stripe along the light-sheet waist has the best axial resolution). Scanning fewer tiles leads to smaller datasets and faster acquisition times. Having near-isotropic raw data also allows visualization without extensive post-processing or deconvolution and aids in segmenting structures of interest. While the mesoSPIM initative will evolve to include improved features in the instruments, we believe that the current configuration already satisfies a wide range of application requirements.

**Existing setups**

As of February 2019, five mesoSPIM V4 (Version 4) setups are operational across various research labs and imaging facilities in Europe:

- Brain Research Institute, University of Zurich (Helmchen group)
- Institute of Neuropathology, University Hospital Zurich (Aguzzi group)
- Wyss Center Geneva (Advanced Lightsheet Imaging Center 'ALICE')
- Center for Microscopy and Image Analysis, University of Zurich
- Sainsbury Wellcome Center for Neural Circuits and Behavior London (Advanced Microscopy Facility)

Several additional mesoSPIM V4 and V5 instruments are currently under construction. Version 5 is the latest (and recommended) design, it incorporates better sample stages with larger travel range. All data in this manuscript was acquired using a mesoSPIM V4.

**Core features**

- Horizontal detection path as in the original SPIM[1] - which allows for sample rotation by suspending the sample from the top without changing the direction of gravity acting on the sample.
- Macro-zoom system (Olympus MVX-10) in the detection path, enabling large FOVs of 2-21 mm in combination with a 1x air objective (Olympus MVPLAPO1x).
- Dual-sided illumination
- Exitation path designed for minimal shadowing artifacts (stripes cause by refraction, scatting, and absorption inside the sample) by using an illumination scheme which delivers comparable results to the multidirectional SPIM (mSPIM)[2].
- Uniform axial resolution across such FOVs by axially scanned light-sheet microscopy (ASLM)[3] based on a tunable lens.
- Large travel range (44.5 mm × 44.5 mm × 100 mm) to accommodate specimens such as a whole cleared mouse CNS without the need for remounting or cutting.

- Compatibility with all major clearing techniques ranging from hydrogel-based methods such as CLARITY[4], hydrophilic reagent-based techniques such as CUBIC[5] to organic solvent approaches such as 3DISCO[6,7] and BABB[8,9].
- Switching between different imaging media optimized for specific clearing methods is accomplished by quick-exchange immersion cuvettes on magnetic mounts and can be done in tens of seconds. Samples are suspended in the immersion cuvettes from the top.
- Sample mounting can be done either in sample cuvettes (as for the COLM setup[10]) or using 3D-printed sample clamps (i.e. for iDISCO[11]). Magnetic sample holders allow for quick sample exchange within tens of seconds.
- Open and modular hardware and software to adapt the instrument to custom imaging requirements.

**Comparison to commercial and open-source light-sheet instruments**

In the light-sheet community several open-source light-sheet projects exist, for example the openSPIM[12] (**openspim.org**) and openSPIN[13] (**https://sites.google.com/site/openspinmicroscopy/**) projects, which aim to provide scientists without prior knowledge in building optical systems with low-cost entry-level light-sheet microscopes. Simlilarly, the mesoSPIM initiative aims to build a community of users and developers of mesoSPIM setups to foster the exchange of ideas and solutions around imaging technologies for cleared tissue. We hope that the mesoSPIM design provides a starting point for other developers to add and share their own improvements and modifications.

However, open-source instrumentation also has issues: Achieving consistent data quality and sample throughput appropriate for publication with low-cost setups is challenging[14,15]. For example, the openSPIM project is highly successful as a tool to train microscopists on how to build and use a light-sheet microscope. However, only a small fraction[16-22] of the 152 citations (as of March 2019) of the original openSPIM publication present original biological research acquired with an openSPIM.

With the mesoSPIM initiative, we hope to provide a mesoscopic imaging instrument for cleared tissue *en par* or better than existing options. The setup is thus suitable for research groups or imaging facilities in search of alternatives to the existing palette of commercial instruments. Typically, we recommend the mesoSPIM to teams in which at least one team member has experience in setting up custom microscopes such as a two-photon microscope or a lattice light-sheet instrument. In addition, we would like to emphasize that in the long run—as with every custom microscope—a mesoSPIM is only as good as its support staff and that experienced local support is absolutely necessary.

Compared to the current generation of commercial light-sheet instruments suitable for cleared tissue, the mesoSPIM is most similar to the LaVisionBiotec Ultramicroscope II. They share the usage of the Olympus MVX-10 macro zoom system to achieve large FOVs in the cm-range. Whereas the Ultramicroscope has a vertical detection path and a light-sheet parallel to the horizontal plane (which simplifies the usage of dipping objectives), the mesoSPIM has a horizontal detection path and a vertical light-sheet (which allows quick exchange of samples and immersion cuvettes and allows sample rotation around a vertical rotation axis). Other commercial instruments such as the Zeiss Z.1 or Luxendo MuVi SPIM CS share this vertical light-sheet geometry but only fit small samples due to travel range limitations (<10 mm in XYZ vs. 44.5 mm × 44.5 mm × 100 mm of the mesoSPIM V4). However, these instruments can be used with high-NA immersion objectives and thus allow higher imaging resolution. This option is currently not implemented in the mesoSPIM and would require the design of special imaging chambers. Another instrument is the ct-diSPIM by Applied Scientific Instrumentation (ASI), which is based on the inverted SPIM (iSPIM) geometry[23] and can fit samples of comparable size to the mesoSPIM but only with a maximum thickness of 5 mm (limited by the working distance of the objectives). With its multi-immersion objectives ($n_D$ = 1.33-1.56) this microscope is also one of the few commercial instruments that is compatible with all clearing techniques without major modifications. However, owing to its light-sheet geometry it lacks a rotation stage for multi-view acquisitions. In addition, it is currently not available with FOVs in the cm-range.

Several existing commercial microscopes also feature imaging modes yielding uniform axial resolution across the FOV. For example, the LaVisionBiotec Ultramicroscope II has a “dynamic focus” mode that mechanically translates the waist location while merging subsequent images. This approach is, however, much slower than the ASLM implementation used in the mesoSPIM. Other commercial light-sheet instruments such as the LifeCanvas SmartSPIM and the 3I light-sheet microscope feature imaging modes with uniform axial resolution that are faster, for example by adopting a tiling light-sheet approach in the 3I instrument[24,25] using spatial light modulators. Both instruments are using vertical detection paths like the LaVisionBiotec Ultramicroscope and the mesoSPIM.

A key advantage of the mesoSPIM compared to all existing commercial setups is that it supports imaging samples immersed in sample cuvettes, an idea that was introduced by Tomer et al. in the CLARITY-optimized light-sheet microscope (COLM)[10]. Because developing mounting strategies for different types of samples[26-29] is a key aspect of using a light-sheet microscope, being able to quickly image samples by putting them in an imaging cuvette renders mesoSPIM usage highly ergonomic. Restricting the volume needed for index-matching the sample is beneficial if the refractive index matching solution (RIMS) is costly. Furthermore, if the excitation light-sheet has to pass through less medium solution, the refractive index inhomogeneities of water-based high-index RIMS media do not degrade light-sheet quality as much as compared to immersing the sample in a larger chamber. For example, the LaVision Ultramicroscope with its large (10 cm) imaging cuvette is less suited for imaging CLARITY samples for that reason. In summary, we believe that the combination of mesoSPIM features is unique and only partially available commercially and that the mesoSPIM constitutes a highly capable multi-user imaging platform for large cleared samples.

### mesoSPIM Budget

While the parts lists for both mesoSPIM version 4 and 5 can be used to prepare a detailed budget (github.com/mesoSPIM/mesoSPIM-hardware-documentation ) a coarse overview is provided in Supplementary Table 1.

| Component | Cost (USD) |
|---|---|
| Optical table | 8000 |
| Camera | 17000 |
| Olympus MVX-10 | 11500 |
| Filter wheel | 7400 |
| Filters | 8500 |
| Laser engine | 40000 – 110000 |
| Mechanical stage | 26000 |
| Imaging Computer | 9000 |
| Electronics | 10000 |
| Optics + Optomechanics | 32200 |
| **Total** | **169600 – 239600** |

**Supplementary Table 1: mesoSPIM V4/V5 budget overview:** The laser engine (fiber-coupled laser combiner with multiple laser lines) is the most expensive single item. For this list, it is assumed that the mesoSPIM will be built in a full configuration with dual-sided illumination and a wide variety of excitation laser lines and emission filters. Prices are based on quotations we obtained for setting up several mesoSPIM instruments in Switzerland and were converted to USD at an exchange ratio of 1:1 (as of February 2019). Owing to import, customs, and shipping expenditures, costs are likely to be considerably different in other countries.

**Supplementary Note: Axially scanned light-sheet microscopy (ASLM): A key mesoSPIM imaging mode**

Users of early light-sheet setups such as the orthogonal-plane fluorescence optical sectioning (OPFOS) instrument by Voie et al.[30,31] or the selective plane illumination microscope (SPIM) by Huisken et al.[1] noted that the quality of the 3D reconstruction of specimens were critically dependent on the effective thickness of the optical sections. As these first-generation fluorescence-based light-sheet instruments relied on Gaussian beams in their illumination paths, the variation of the beam profile along the excitation direction caused images to be sharp in the center but increasingly blurry towards the edges if the datasets were viewed from the side (in the XZ-plane). Therefore, the illumination path was usually set up in a way that the Rayleigh-range $z_R$ of the illumination beam was approximately twice the FOV size provided by the camera. In this way, the light-sheet thickness varied only by $\sqrt{2}$ across the FOV. However, this is unsatisfactory at low magnification (<20×) as the sub-µm lateral resolution provided by the emission path outweighs the multi-µm axial resolution provided by the light-sheet[32].

A simple solution is to restrict the readout of the 2D imaging detector to a region around the center of the light-sheet waist and move the sample through this region while taking sequential images[33] – essentially performing extensive tiling acquisitions. This takes more time because for each z-plane the sample has to be moved several times in the lateral direction. Essentially, the parallel 2D readout (which massively speeds up volume acquisition in a SPIM compared to a point-scanning microscope) is reduced to a quasi-1D readout if the best axial resolution is desired. Nonetheless, the slow acquisition allows the resulting datasets to exhibit a high degree of axial uniformity. As most of the sensor area is thus not used for collecting data, it made sense to simplify the sensor from a 2D array to a Time Delay and Integration (TDI) detector where the movement of the sample is synchronized with the readout of the camera – an idea first implemented in a thin-sheet laser imaging microscope (TSLIM) in 2010[34].

During the same time, other groups explored the usage of a confocal slit as a spatial filter (similar to a confocal microscope) to improve the axial resolution and uniformity of light-sheet microscopes[35] in a digitally scanned light-sheet microscope (DSLM)[36]. As the detector 'sees' only ballistic photons from the light-sheet, this approach helps to reduce the impact of scattering. With the introduction of scientific sCMOS cameras into microscopy, it was noted that the rolling-shutter readout mode of such a camera could be used to approximate a confocal slit[37]. The rolling shutter restricts the active line to a few rows, which can be synchronized with the DSLM scanning motion to create a light-sheet.

At the same time, several groups explored the capabilities of other excitation geometries to create thinner as well as more uniform light-sheets by employing two-photon excitation[38], non-diffracting Bessel[39] and Airy beams[40] and combinations thereof[41]. In 2014, Dean et al.[42] used a tunable acoustic gradient (TAG) lens to create an extended focus out of a Gaussian beam for more uniform axial resolution. In 2015, the same group combined the remote focusing system by Botcherby et al.[43] with the externally (controlled) rolling-shutter readout of a modern sCMOS camera to form an axially swept light sheet microscopy (ASLM) instrument[3]. Briefly, the waist motion through the sample was synchronized with the readout of the rolling shutter so that during the sweep the waist location tracked the active line on the camera (see **Supplementary Figure 1**). This instrument yielded isotropic 390-nm resolution throughout a $216\times162\times100$ $\mu m^3$ imaging volume. The major drawback is similar to earlier approaches, namely a longer acquisition time for the creation of 3D data because for each z-plane, additional time has to be spent sweeping the light-sheet across the FOV to collect a sufficient number of signal photons. With tunable optical elements, as employed in the mesoSPIM systems, however, there are no heavy mechanical parts that have to be moved,

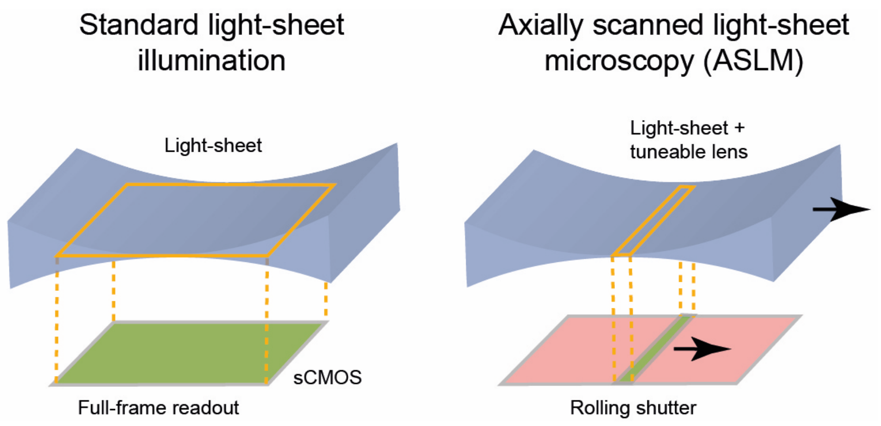

**Supplementary Figure 1: Comparison of standard light-sheet illumination and axially scanned light-sheet microscopy.** With standard illumination, the whole frame is read out at once, which leads to non-uniform axial resolution across the FOV. In ASLM mode, the readout of the camera is synchronized with the motion of the light-sheet waist through the sample. The final image contains only signal from the waist region which leads to uniform axial resolution across the FOV.

which leads to a considerable speedup. However, each plane has to be illuminated for a longer time to perform the axial sweep, which can lead to increased photobleaching.

In 2016, Hedde and Gratton[44] used an electrically tunable lens (ETL) as the remote focusing device for ASLM, a device which can also be used to tune the excitation light-sheet to the shape of the sample[45] by positioning the waist at regions of interest, for example to illuminate the outline of a Drosophila embryo as efficiently as possible. A comparable effect to ASLM can be achieved by tiling the excitation beam waist[24,25] using spatial light-modulators (SLM). A further option is to use lattice light-sheet illumination[46,47] which has not been implemented yet for cm-sized FOVs.

While testing early mesoSPIM prototypes (Version 1 to 3), which employed cylindrical lenses or scanned Gaussian beams to create the light-sheet, we noted that especially in large (cm-sized) samples, achieving uniform axial resolution is a key requirement. At 488 nm and at $n_D = 1.45$, even a light-sheet with a 5-µm (FWHM) waist in the center expands to approximately 400 µm at the edges of a 13.4 mm FOV (a size, which is for example needed to image a whole CLARITY-cleared mouse brain). As noted above, this loss of axial resolution means that most light-sheet instruments are used with extensive tiling acquisitions[10,35,48], yielding hundreds of GB per brain. We reasoned that implementing a light-sheet illumination mode with more uniform axial resolution would both lead to higher data quality and easier data analysis and allow users to screen samples at near-isotropic resolution before committing to generate TB-sized datasets. We noted that for many labs in the early stages of adopting a clearing method, sample quality was usually not sufficient to warrant generation of high-resolution data across the whole sample. However, visualizing at which locations inside a whole-brain sample clearing or labeling quality were insufficient was not straightforward at low magnification because a light-sheet with 40-100 µm thickness (with a Rayleigh range $z_R$ tuned to large FOVs) led to extensive axial blurring of features. Therefore, we reasoned that achieving a thinner and more uniform effective light-sheet thickness would allow better visualization of features at low magnification and possibly also improve data quality at higher magnification. Among the possible options (ASLM, tiled light-sheets, nonlinear excitation, Bessel or Airy-beams, or a lattice light-sheet approach) we selected ASLM. Given that our macro-light-sheet instrument was supposed to have FOVs ranging from 2-20 mm and be capable of exciting a wide variety of fluorophores, we deemed ASLM as the simplest single-photon approach to be most appropriate and cost-efficient. Given our positive previous experiences with electrically tunable lenses in microscopy[49,50], we decided to include ETLs as remote focusing units in the optical path as suggested by Hedde and Gratton[44].

To demonstrate that the resulting mesoSPIM configuration allows a substantial increase in axial resolution uniformity across the FOV, we cleared mouse brains, in which vasoactive intestinal peptide (VIP) interneurons express tdTomato, with a passive CLARITY protocol (**see Supplementary Note 9 – Sample preparation**). When scanning this sample with a Gaussian beam with the ASLM-mode switched off, labeled neurons could be distinguished in the axial direction in the center of the FOV whereas at the edges of the 13.29 mm FOV (zoom 1x) cells overlapped in Z (**Fig. 1b**; **Supplementary Figure 2a,c,e**). After switching the ASLM mode on, however, neurons could easily be distinguished in the XZ plane (**Fig. 1b**; **Supplementary Figure 2b,d,f**). In a volume rendering, the same effect is apparent (**Supplementary Videos 1 & 2**). These datasets were acquired within 8 minutes (2-µm z-step size). At 4x magnification and 1.6 µm x 1.6 µm x 2 µm sampling, switching the ASLM mode on allows for the visualization of axons in the XZ-plane across the entire 3.3-mm FOV (**Supplementary Figure 4**, **Supplementary Video 3**). Allowing for ASLM also means that unlike most other light-sheet microscopes, a mesoSPIM allows users to control the effective thickness of the light-sheet to a high degree. This freedom comes with a considerable learning curve as users have to learn how to optimize the ASLM sweep parameters from the live (XY) image. Sub-optimal settings of the ASLM parameters will result in a loss of optical sectioning quality (see **Supplementary Figure 5**).

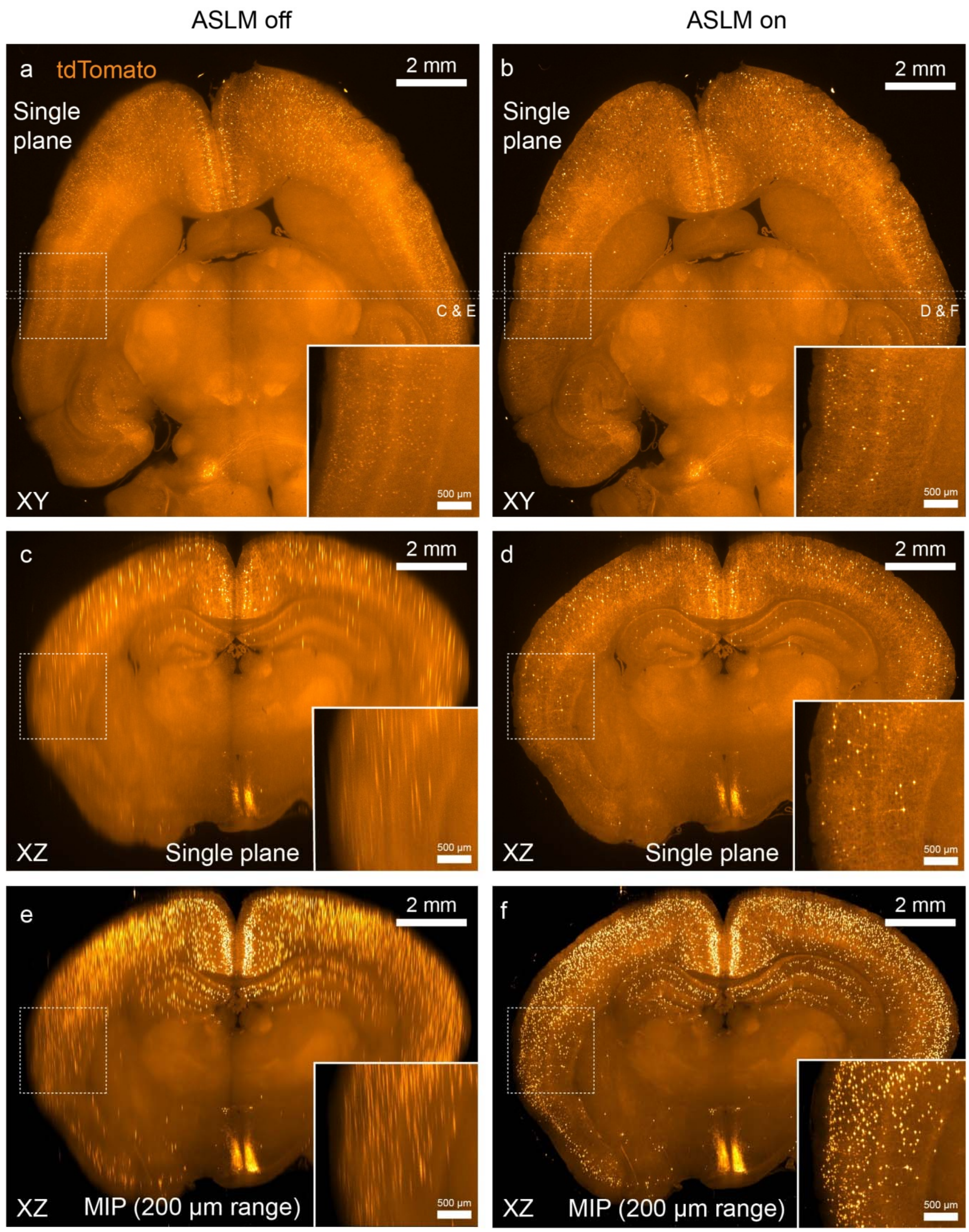

**Supplementary Figure 2: Comparison of whole-brain VIPCre-tdTomato datasets acquired with and without ASLM at 1× magnification.** a) & b) Single planes acquired from a passive CLARITY-cleared VIPCre-tdTomato mouse brain with ASLM switched off (left) and ASLM switched on (right). Whereas in the center, both images look similar, with ASLM on, far fewer cells are visible because the optical section is thinner. c) & d) Single resliced (XZ) plane at the location indicated in a) & b): Due to ASLM, the uniformity of the axial resolution is much better which renders single cells visible in the XZ plane at the edges of the FOV. e) & f): Comparison of maximum intensity projections at the same location.

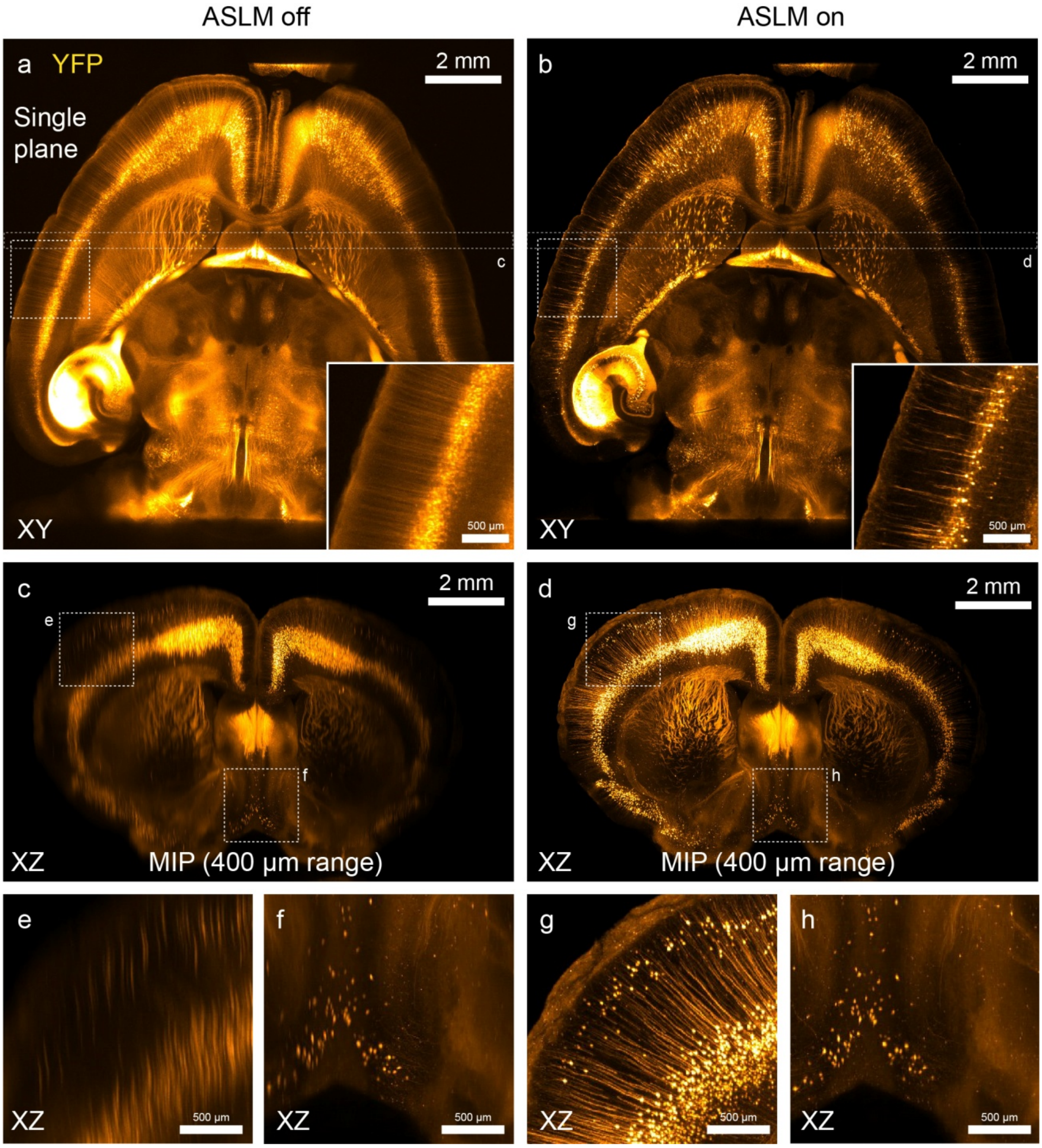

**Supplementary Figure 3: Comparison of whole-brain Thy1-YFP datasets acquired with and without ASLM at 1x magnification. a) & b)** Single planes acquired from an active CLARITY-cleared Thy1-H-YFP mouse brain with ASLM switched off (left) and ASLM switched on (right) at low magnification (Zoom 1x). Whereas in the center, both images look similar, with ASLM on, far fewer cells are visible because the optical section is thinner (insets). In addition, sections of single dendrites become visible. c) & d) Comparison of maximum intensity projections (MIP) in the XZ plane. The location of the coronal MIP is indicated in subpanels a) and b). e)-h) Details from subpanels c) & d): Whereas along the midline (subpanels f) and h)), the improvement in axial resolution is less pronounced, there is a drastic increase in axial resolution at the edges of the FOV (subpanel e) vs. g)) which renders single dendrites visible in the XZ plane.

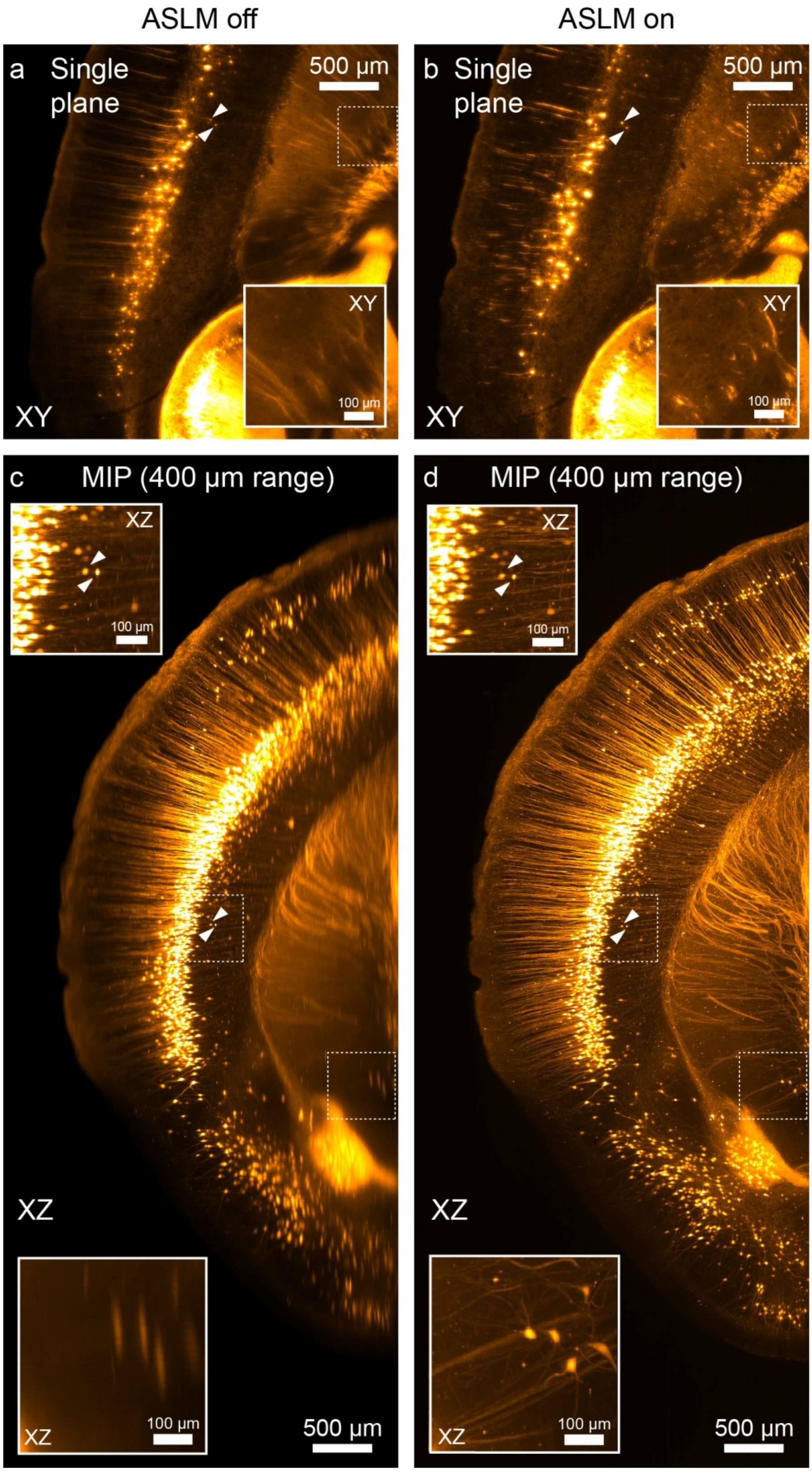

**Supplementary Figure 4: Comparison of Thy1-YFP datasets acquired with and without ASLM at 4x magnification:** a) & b) Comparison of single planes acquired with ASLM switched off (left) and ASLM switched on (right). The more uniform thickness of the optical section reduces the blur around features at the edges of the 3.3-mm FOV. c) & d): Comparison of XZ maximum intensity projections. Whereas cells close to the location of the light-sheet waist in the ASLM-off condition (marked by arrows) do not show a loss of axial resolution in ASLM mode (top insets), neurons at the edge of the FOV (bottom insets) show a massive improvement in axial resolution. Passing axons at that location are virtually invisible without ASLM.

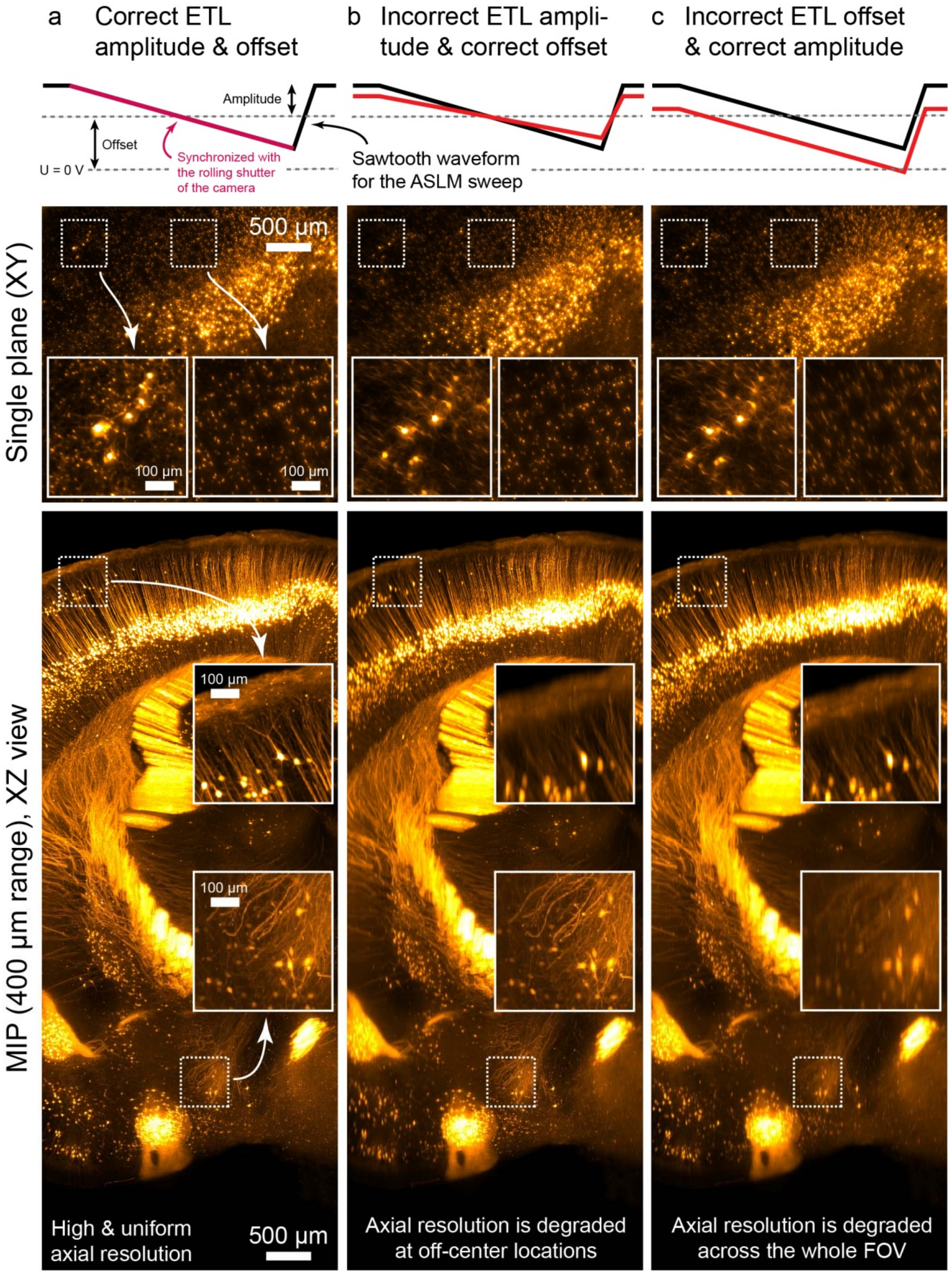

**Supplementary Figure 5: Choosing the correct ETL parameters is important to achieve uniform axial resolution.** Shown is a comparison of z-stacks in both the XY and XZ planes acquired at 4x magnification in a CLARITY-cleared Thy1-YFP mouse brain. The slow linear section of the ETL waveform is synchronized with the rolling shutter of the sCMOS camera. a) The optimal choice of the ETL parameters yields high and uniform axial resolution across the FOV. If either the ETL amplitude (b) or offset (c) are chosen incorrectly, axial resolution is degraded.

The ETL parameters (ETL amplitude and offset of the ramp, see **Supplementary Figure 12**) depend on a variety of imaging conditions:

- A change in zoom leads to a change in ETL amplitude: For example, smaller FOVs at higher magnification mean that the ASLM sweep should cover a smaller distance along the illumination direction which corresponds to a reduction in ETL amplitude.
- The ETL offset mostly depends on the bulk refractive properties of the imaging medium – switching to a different clearing medium (with different $n_D$) also leads to a required change in offset.
- Switching the excitation wavelength requires changing the ETL offset as the dispersion of the immersion medium is wavelength-dependent.

For all of the above, the mesoSPIM software provides reasonable presets that have to be calibrated for a certain immersion medium when setting up the instrument. However, the following should be noted:

- Both ETL amplitude and zoom can depend on the local refractive properties of the sample – we noticed that even extremely well cleared and index-matched samples still show some variation of optimal ETL parameters in mosaic acquisitions.

In our experience, despite extensive index-matching, every cleared sample still behaves like a lens. This is not surprising given that even very small index variations across slightly tilted interfaces can lead to detectable refraction across cm-sized samples. For an experienced user imaging a familiar sample, manually optimizing the thickness of the light-sheet locally is a process that takes only a few seconds. However, we also noted that in samples with insufficient signal intensity or unsatisfactory clearing quality, an optimum can be extremely hard to find. If a user has never seen a certain type of labeling before, understanding how to read the XY image as an indication of optical sectioning thickness requires some experience and interpretation. For example, in samples with a uniform nuclear staining or with fluorescent beads, the number of visible spots can be used as a simple indication of light-sheet thickness – the fewer spots, the better. This also means that lower bulk image intensity is better. In samples in which neurons are labeled in their entirety, the effective light-sheet thickness is the better the shorter the visible sections of dendrites and axons appear in the XY image. However, especially in regions of interest (ROIs) that contain background autofluorescence and a small subset of passing axons, it is important to optimize for the axons only and not for general image contrast – a process that can be complicated by the rapid bleaching of background autofluorescence in CLARITY-cleared samples. As we noted that especially difficult-to-optimize subregions of specific samples require quite some background knowledge from the user, we currently have not automated the choice of ETL parameters. Nonetheless, we foresee that automation should be possible in a subset of samples by using on-line optimization similar to the techniques implemented in the current generation of smart light-sheet microscopes for developmental biology[51]. In fact, an optimization step for the overlap of illumination and detection for tiled acquisitions in cleared samples was already implemented in the COLM setup software by Tomer et al.[10], but it comes with considerable overhead in acquisition time.

The local refractive properties of cleared samples we highlighted above also lead to an important consideration on how to acquire data with a mesoSPIM: We strongly recommend acquiring data from well-cleared samples with a single illumination direction only, as stacks taken with double-sided illumination in the ALSM mode can show axial doubling of cells (see **Supplementary Figure 6**), indicating that the light-sheets are not overlapping properly. Importantly, this happens even though the light-sheets were co-aligned in some other part of the sample along the z-axis (see **Supplementary Figure 6**). Especially when using light-sheets with an effective thickness below 10 µm such effects are highly prevalent, but tend to be hidden when thicker light-sheets are used. If two illumination directions are required, they should be acquired sequentially and merged during data processing.

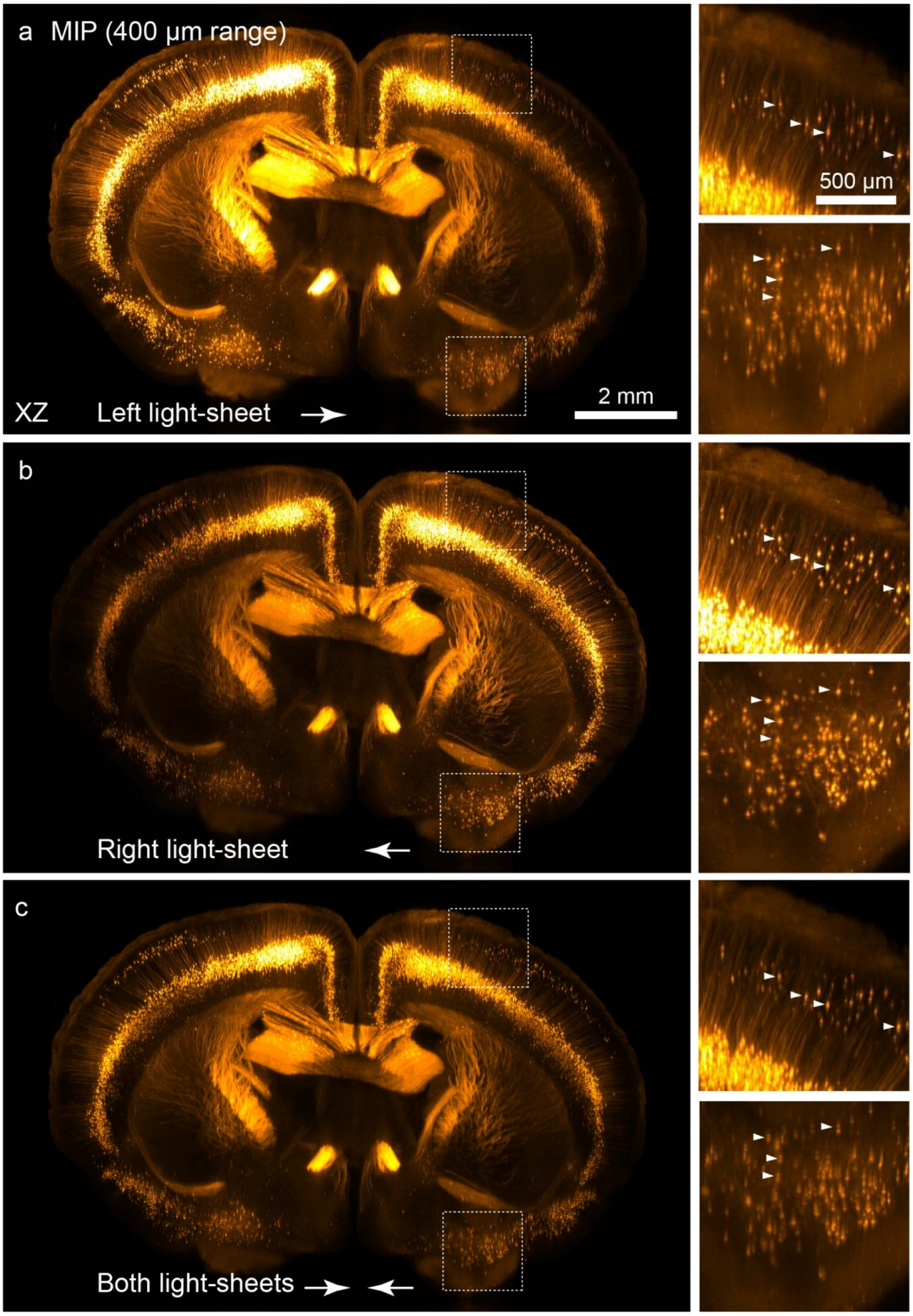

**Supplementary Figure 6: Double-sided illumination can lead to location-specific doubling of axial features.** Shown is a 1x overview stack taken from a CLARITY-cleared Thy1-YFP mouse brain. a) & b): Maximum-intensity projected (MIP) reslices (XZ plane) from datasets taken with illumination from the left or right. A direction-dependent intensity and resolution gradient is visible. c) When acquiring data with both light-sheets, the illumination is more uniform, however, cells which are located in ventral regions appear twice (bottom inset). This artifact is not visible in the dorsal part of the datasets (top inset) which indicates that in well-cleared specimens, light-sheet co-alignment is strongly dependent on the local refractive properties of the sample.

**Supplementary Note: Optical design of the mesoSPIM setup**

Key design decisions during the optical design for the mesoSPIM included:

- Olympus MVX-10 macroscope in the detection path with a MVPLAPO 1x objective for FOVs of 2-21.8 mm (inspired by the LaVision Ultramicroscope I).
- Horizontal detection path as in the original SPIM[52] to simplify sample exchange (from the top) and sample rotation (direction of gravity acting on the sample does not change during rotation).
- Use of a Hamamatsu Orca Flash 4.0 V2 or V3 to allow the usage of the rolling shutter ("light-sheet mode") for axially scanned light-sheet microscopy (ASLM)
- The light-sheet has to be at least as wide as the FOV of the microscope.
- The remote focusing system to translate the waist location trough the sample needs to have a tuning range >22 mm with typical immersion media (to allow ASLM even at the lowest detection zoom settings).
- To generate z-stacks, the light-sheet is stationary and the sample is translated through the sheet.
- The light-sheet illumination should be multidirectional (mSPIM[2]) to minimize the amount of shadowing artifacts.
- The microscope should support single-photon excitation wavelengths from 405 nm to 650 nm and into the NIR region, if possible.
- Co-alignment of both light-sheets should be easy.

The choice of the Olympus MVX-10 macroscope as the core component of the detection path led to the following decisions:

- As the MVPLAPO 1x is an air objective, the samples need to be immersed in an immersion cuvette of sufficient size to allow sample translation and rotation. We opted to use standard glass or quartz macro-fluorescence cuvettes from vendors such as

Hellma or Portmann Instruments which are available in sizes ranging from 30x30x35, 40x40x45 to 50x50x50 $mm^3$ and can be custom made to user requirements.

- As samples come in all kinds of sizes and shapes, the distance between the wall of the immersion cuvette needs to be adjustable to keep the amount of medium between the light-sheet and the objective to a minimum, but allow for enough travelling range to take a z-stack of the region of interest.
- This in turn requires two translation stages along the detection axis: One for translating the sample through the light-sheet to generate z-stacks and one for focusing the detection optics.
- In addition, the MVX-10 requires the use of 32-mm filters in the detection filter wheel to have sufficient free aperture (necessitating custom filters)

The parameters of the detection system thus dictated the design of the illumination optics to a high degree. Early tests with light-sheet illumination using cylindrical lenses showed that shadow artifacts can lead to many stripes in the images, which we deemed inacceptable. For example, in a modified openSPIM for cleared tissue[18], shadow artifacts lead to a severe degradation in image quality. Therefore, we concluded that shadow reduction – for example in the form of multidirectional SPIM (mSPIM)[2] – is necessary. In this solution, the light-sheet is rapidly pivoted by a resonant scanner so that the shadows cast by absorbing structures in the sample are averaged out. While designing a mSPIM for 2-20 mm FOVs is feasible, we adopted a design similar to an digitally scanned light-sheet microscope (DSLM[36]). In a DSLM, the light-sheet is created by rapidly scanning a Gaussian beam in a plane. Typically, in a DSLM the NA of the laser beam is chosen such that the Rayleigh length is on the order of half of the FOV of the microscope camera so that the resolution variation of the axial resolution across the FOV is not too severe. This leads to low excitation NAs for large FOVs. As noted by Fahrbach et al.[53], a scanned Gaussian beam leads to a reduction in shadow artifacts – though not as strongly as using Bessel beam excitation. The effectiveness of shadow reduction is related to the NA of the

Gaussian illumination beam: The higher the NA, the shorter the shadow cone behind an absorbing object. For large FOVs, the necessary low excitation NAs can thus lead to severe shadow artifacts. In an ASLM setup, however, there is no trade-off between NA and FOV: The higher the illumination NA, the smaller the beam waist which in turn is converted to a low effective light-sheet thickness by the ASLM sweep. A side effect of ASLM is thus that it allows using higher excitation NAs (compared to a standard DSLM with the same FOV) which in turn lead to decreased shadow artifacts.

Initial tests with mesoSPIM prototypes showed that a scanned Gaussian beam with an excitation NA of 0.14 - 0.15 yielded acceptable reduction of shadow artifacts over the FOV. The scan lens path has to be telecentric, otherwise the height of the light-sheet (in the Y direction) does not stay constant during remote focusing as required for ASLM. Any change in height will also be a nuisance as the light-sheet intensity will vary across the FOV: If the light-sheet height is not constant, the available laser power will be more concentrated in certain parts of the FOV (see **Supplementary Figure 7**). In typical DSLM instruments, the excitation path is similar to typical confocal excitation paths with a combination of scanners, a scan and tube lens and a microscope objective. As we needed more than 20-mm FOV due to our large detection FOV, we deemed standard microscope optics as inadequate as there are no low-magnification objectives with sufficient working distance (> 25 mm) and NA (>0.1). One of the closest approximations to the mesoSPIM requirements, the Olympus XLFLUOR 4x NA 0.28 with a working distance of 29.5 mm is designed for a field number of 22, corresponding only to a 5.5 mm FOV. However, the design of the scan path can be simplified by removing the tube lens and objective altogether: By combining a single-axis galvo scanner with a scan lens of sufficient NA and working distance, creating a light-sheet of more than 20-mm height after the scan lens should be feasible (**Supplementary Figure 7)**.

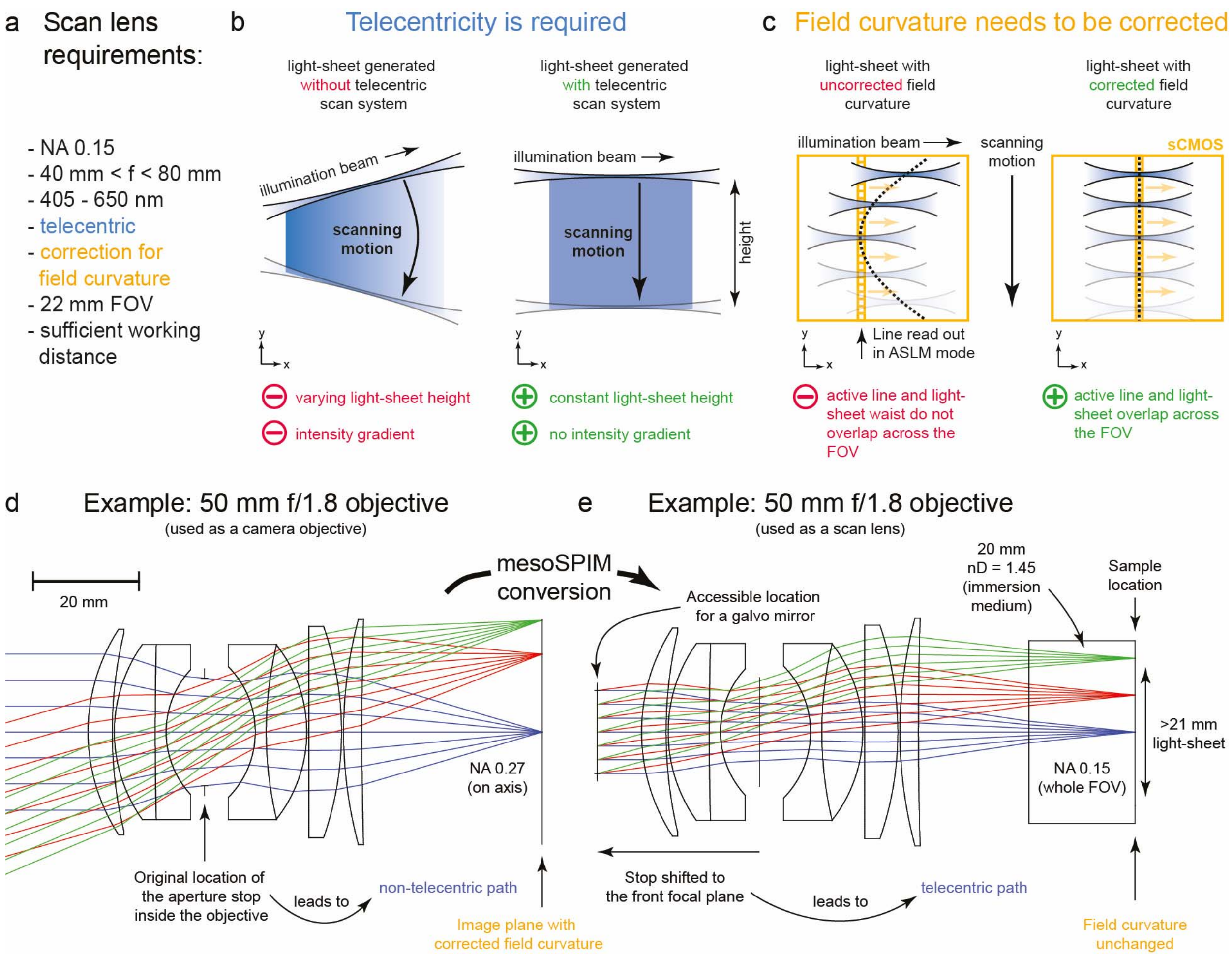

**Supplementary Figure 7: Challenges and solutions in the optical design of the mesoSPIM excitation scan lens.** a) List of scan lens requirements. b) The scan system has to be telecentric to avoid a varying height of the light-sheet and the associated intensity gradient. c) Field curvature has to be corrected allow for the ASLM mode to work properly for large FOVs. d) Example design of a 50 mm f/1.8 objective (from Laikin[54]) showing the original design conditions for this camera lens. e) Conversion of the lens into a scan lens. Please note this design is close to the Nikon 50 mm f/1.4 G objectives used in the mesoSPIM to show the concept of shifting a stop, but currently, the design of the lenses used in the mesoSPIM is unavailable to the authors.

Therefore, we were faced with the challenge of requiring a scan lens with the following parameters (**Supplementary Figure 7**):

- NA 0.15
- Focal length in the range from 40 mm to 80 mm (larger focal lengths lead to overly large galvo scanners).
- Telecentric (on the finite conjugate side) to generate a light-sheet with uniform height for ASLM.
- Achromatic from 405 to 650 nm

- Corrected for field curvature to have a straight waist for ASLM.
- Ideally, diffraction limited across a 22-mm FOV / image circle.
- Sufficient working distance

For comparison, standard scan lenses for confocal microscopy have specifications close to the Thorlabs SL50-CLS2:

- f = 50 mm
- NA 0.04 (4-mm input beam)
- Telecentric
- Field flattened
- Correction from 450 to 1100 nm
- 26.4 mm working distance
- 15.5 mm image circle at 587.6 nm

In optical design, scaling up such a field-flattened telecentric scan lens (to obtain 3x the NA and 25% larger FOV) is a formidable challenge that will result in a costly custom design. Apart from the upscaled NA, a key factor in the cost of such a design is the achromaticity requirement. Moreover, such a scan lens design cannot be carried out in a fully achromatic fashion without movable elements if it is supposed to work with varying immersion media: As the dispersion of clearing media and immersion oils varies significantly between different clearing media, the scan lens would have to contain a correction collar which in turn leads to cost increases. Upon closer inspection, however, we noted that in fixed and cleared tissue, it is unlikely that simultaneous illumination with multiple laser lines is required: We were planning to use only a single detection camera and in fixed tissue, different color channels can be acquired by successive z-stacks if the sample does not drift or change over time. In addition, we were planning to have a tunable lens in the path to allow for ASLM, which is basically a remote focusing arrangement. Hence, if there has to be a tunable lens anyway and if we plan to use only one excitation wavelength between 405-650 nm at a time, the axial chromatic aberration of the scan lens can be corrected by simply applying an offset to the ETL driving signal. This drastically simplifies the optical design.

In light of these modified requirements, the opportunity for an optics "hack" opened up: In optical design, custom designs tend to be costly and take long to manufacture, but things are much easier when mass-produced optical systems can be creatively repurposed for the required application. We noted that the requirement for working distance, field flattening, and FOV are basically covered by camera lenses: Typical consumer DSLR (digital single-lens reflex camera) lenses have long image-side working distances (to make space for the moving mirror, typically 44-46 mm) and a flattened 24x36 mm field to cover full-format CMOS detectors (**Supplementary Figure 7d**). In addition, there is a massive selection of focal lengths in the 40-70 mm range available as these are considered "standard" or "portrait" lenses. Typical 50-mm DSLR lenses have an f-Number (f/#) of 1.4 (or NA 0.35, $f/\# = 1/(2NA)$) but are not diffraction-limited under such conditions. However, such a lens can be "stopped down" to NA 0.15 (corresponding to approx. 15-mm beam diameter), which yields a reasonable size for a galvo mirror when using this lens as a scan lens. Therefore, such DSLR lenses were promising candidates for the mesoSPIM excitation path. The missing requirement from our initial list is telecentricity – DSLR lenses are usually not single- or double-telecentric (except some macroscopy lenses) and have an inaccessible aperture stop inside the lens. However, a lens can be forced to perform as a telecentric system by placing the aperture stop at a particular position. In the case of a laser scanning system, the galvo scanner creates a stop location (where ray bundles directed at different image locations originate). If the position of the scanner coincides with the front or back focal plane of the optical system, the system will perform telecentrically (**Supplementary Figure 7e**). Importantly, forcing an optical system to accept a different stop location will result in introducing additional off-axis aberrations such as coma or astigmatism, the degree of which can be calculated using stop-shift formulae[55]. We deemed this risk acceptable as NA 0.15 is much lower than the original design NA of DSLR lenses. From this point on, the challenge was to find a 50-mm DSLR lens, which had an external front focal plane (FFP) and a sufficient distance between FFP and front lens (at least 7.5 mm to place a 15-mm scan mirror there). We therefore screened a series of consumer DSLR lenses for accessible FFPs using an autocollimator and found a lens with approximately 12 mm distance between the FFP and the vertex of the front lens in the Nikon AF-S 50 mm f/1.4 G. In this lens, however, the FFP is located within the lens hood which means that several plastic parts have to be removed before a scan mirror can be placed close enough. The modification process is documented on the mesoSPIM wiki: https://github.com/mesoSPIM/mesoSPIM-hardware-documentation/wiki/50mm_lens_modification

Be aware that this modification is likely to void the warranty of the lenses and will render their autofocus unusable in a standard DSLR as the focus encoder has to be removed. Nonetheless, we consider this acceptable as each objective / scan lens costs only around 0.2% (or approx. 500 USD) of the total cost of a mesoSPIM with 6 laser lines.

Therefore, the combination of this 50-mm DSLR/"scan lens" and 15-mm beam diameter (NA 0.15) was chosen as basic components for designing the rest of the excitation path. 15-mm scanners from galvo suppliers such as Citizen Chiba, Cambridge Technologies or Scanlab are made-to-order items with long lead times and require custom power supplies and driver boxes. Therefore, the mesoSPIM excitation path is also compatible with 10-mm scanners such as the Thorlabs GVS211/M, which are off-the-shelf items but lead to a reduced excitation NA of 0.1. To ensure telecentric conditions during refocusing/translation of the waist of the excitation light-sheet, the ETL has to be placed in a conjugate pupil relative to the galvo scanner[50], which can be achieved by a 4f system. To allow for the maximum possible tuning range, the magnification of the relay should be as low as possible. Therefore, we opted to use a 1:1 relay and an ETL with 16-mm free aperture (EL-16-40-TC-VIS-5D-1C, Optotune AG). ETLs with large free apertures tend to have slow step responses and are

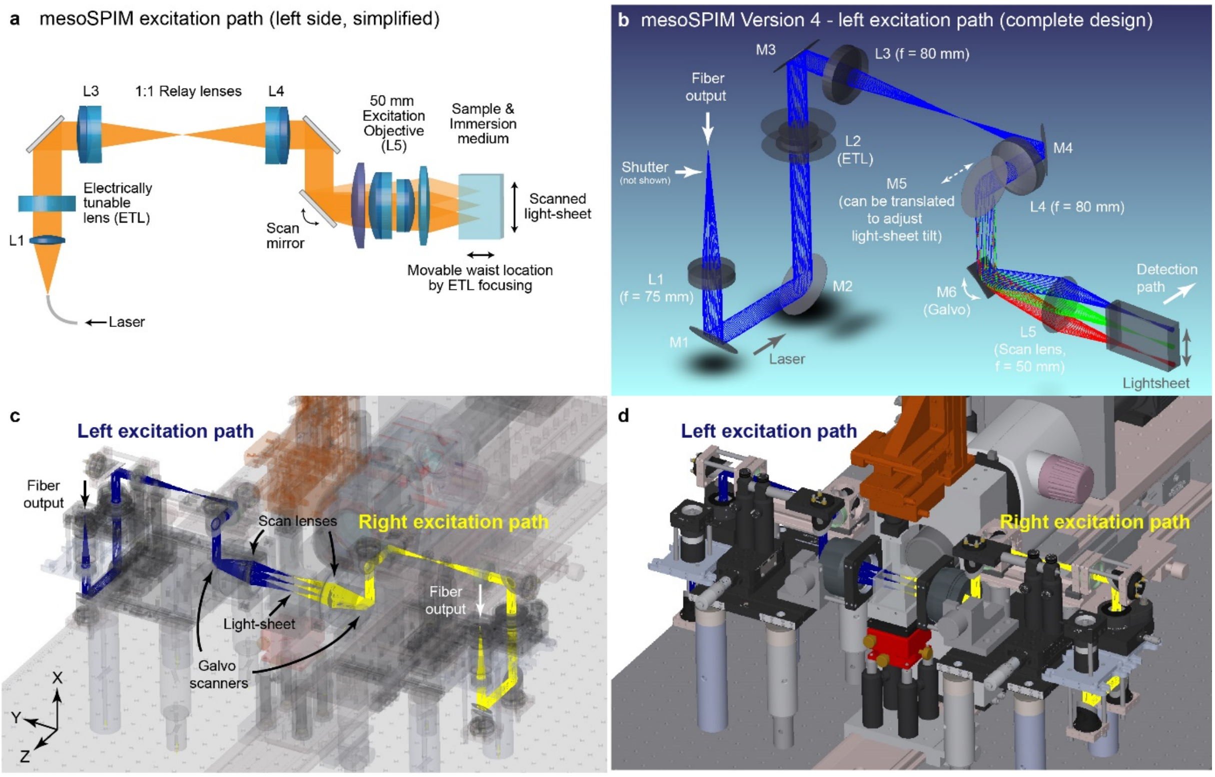

**Supplementary Figure 8: Optical design of the mesoSPIM excitation path.** a) Simplified optical path: Laser light from a single-mode fiber is collimated and then sent to an electrically tunable lens (ETL). The ETL is placed in the conjugate pupil of the galvo scan mirror via a 1:1 relay system (4f system) composed of two achromats (L3 and L4). The galvo scanner creates the light-sheet in combination with a f = 50 mm scan lens. The location of the light-sheet waist along the excitation direction can be changed by changing the ETL focus. b) Full design of the left excitation path showing additional fold mirrors which were required to place the ETL in a vertical section of the optical path to avoid gravity-induced sag and to allow for adjustment of the light-sheet tilt using the translation of mirror 5 (M5). c) & d) CAD renderings showing how the left and right optical paths are placed inside the mechanical design.

thus not suitable for very fast focusing, but we reasoned that the frame rate of the microscope would rarely exceed 10-20 Hz and thus, the 30-ms step response and settling time of the EL-16 does not constitute a major limitation. The ETLs were selected for low wavefront aberrations by the manufacturer. For optimum performance, ETLs should be placed in a vertical section of the optical path because otherwise gravity-induced sag of the liquid leads to non-rotationally symmetric deformation of the lens membrane, which in turn leads to additional aberrations such as coma. This necessitated the inclusion of additional fold mirrors between ETLs and galvo scanners. The final mesoSPIM V4 optical path is shown in **Supplementary Figure 8**. The optical path of the mesoSPIM V5 is identical.

In setting up most light-sheet microscopes with dual-sided illumination paths, co-alignment of both light-sheets is a challenge. This is caused by the need to include enough degrees of freedom in the excitation path to vary the position of the light-sheet along the detection axis and its tilt as independently as possible. While the tilt of the light-sheet can easily be adjusted using a fold mirror with kinematic mount upstream of the galvo scanners, adjusting the tilt angle requires translating the footprint of the excitation beam on the scan lens along the detection axis. We therefore included a fold mirror that can be manually translated and adjusted in angle to simplify this process (**Supplementary Figure 8b**). Additionally, it is aided by a light-sheet alignment mode in the microscope software that interleaves the illumination direction between subsequent frames in live mode.

**Supplementary Note: Description of the mesoSPIM setup**

Detailed parts lists, drawings and CAD files for mesoSPIM version 4 and 5 are available on the mesoSPIM hardware documentation repository: https//github.com/mesoSPIM/mesoSPIM-hardware-documentation

**Guidelines:**

For the optomechanical design, we adhered to the following guidelines:

- The microscope has to be as modular as possible to allow for future modifications.
- Extensive use of rail systems (Qioptiq X95, FLS95 and FLS40) and cage systems (Qioptiq Microbench & Thorlabs 30-mm cage) to aid in modularity and ease of alignment.
- The number and complexity of custom parts has to be kept to a minimum (a mesoSPIM V4 contains only 15 custom parts).
- The detection axis is located 225 mm above the plane of the optical table to allow for sufficient vertical movement of samples.
- The excitation system is located on a horizontal breadboard 185 mm above the plane of the optical table (40 mm below the detection for FLS40 compatibility).

**Optical table:**

With a weight in excess of 50 kg, we recommend placing a mesoSPIM on a dedicated optical table. The total footprint of a mesoSPIM is approximately 1.1 m x 0.75 m x 0.7 m (length x width x height). The detection axis of the microscope is defined by a central FLS-95 rail, which carries the detection system and immersion cuvette (see **Supplementary Figure 9**).

**Excitation lasers:**

The excitation path consists of a laser combiner unit coupled to two single-mode fibers, one for each excitation path. Two of the existing mesoSPIM instruments have Omicron SOLE-6 laser with 405, 488, 515, 561, 594 and 647 nm excitation lines and a 50:50 split between fibers. Three mesoSPIMs have two Toptica MLE laser combiners with 405, 488, 561, 640 nm lines, one for each excitation path. In general, it is recommended to use laser combiners with the highest available output powers, each line should have at least 30 mW on the sample to properly illuminate the full FOV (which can be up to 22 mm). The laser selection should take the following considerations into account:

- Relatively high laser powers are required for imaging endogenous labels in samples cleared with methods with loss of fluorescent proteins (in our experience especially passive and active CLARITY).
- In contrast, well-stained samples – for example, after amplification with secondary antibodies such as in iDISCO – generally leads to high signal levels and thus can be imaged at lower laser intensities.
- If it can be anticipated that only very little imaging will be done in samples requiring FOVs of more than 5 mm, lower laser power specifications will be sufficient as the available power will be concentrated onto a smaller light-sheet.
- We do not recommend having a 405-nm laser line unless it is absolutely needed – in general, the penetration depth even in cleared samples at this wavelength is very low[56] and it is recommended to use red labels instead of UV-excitable dyes.
- White-light laser sources are not recommended (though not tested) as it can be expected that the axial chromatic aberration of the mesoSPIM excitation path will lead to worse effective light-sheet thickness because ETL sweep parameters depend on the dispersion of the immersion medium.

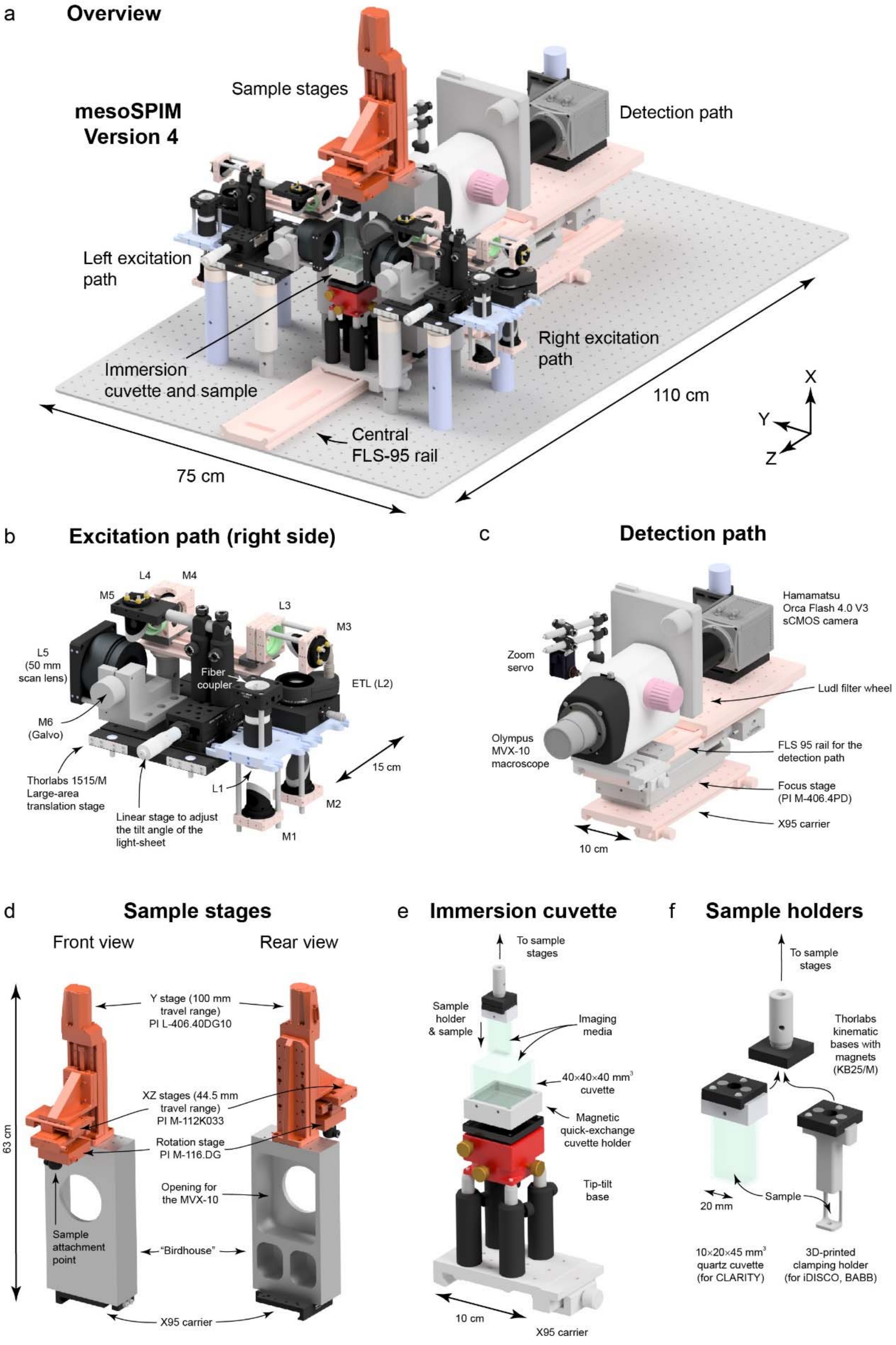

**Supplementary Figure 9: mesoSPIM mechanical design.** a) Overview rendering. b-f): Core mesoSPIM components ranging from the excitation path to the sample holders. For a mesoSPIM Version 4, a total of 15 custom parts are necessary.

Most existing mesoSPIMs are part of either research labs or imaging facilities with a wide range of clearing methods available. Having sufficient laser powers and a wide selection of laser lines available therefore is beneficial to support all kinds of imaging needs. Nonetheless, high-power laser combiners with 4 or 6 wavelengths are a major cost factor and account for 30-50% of the total cost of a mesoSPIM setup. If a mesoSPIM has to be built on a lower budget, we recommend starting with a single laser line and a single excitation path. If multiple lines are required, building an open-source laser combiner might be an option.[57]

**Excitation path:**

Light from the laser combiners is directed to each excitation path (left & right) via a single-mode fiber (**Supplementary Figure 8 and 9b**). After passing a shutter (Thorlabs SHB025), the laser light is collimated by an achromat (f = 75 mm; Thorlabs AC254-075-A-ML) to approximately 15-mm beam size. It is then directed to an electrically tunable lens (ETL, Optotune EL-16-40-TC-VIS-5D-1C), which can create both a converging and diverging beam in combination with a Optotune EL-E-4-i lens driver that is configured for external control via an analog input. The beam is then directed through a 1:1 relay comprising two visible achromats (f = 80 mm; Qioptiq G063200000). Via a series of elliptical 1-inch fold mirrors (Thorlabs BBE1-EO2), the beam is directed towards a galvo scanner for which the following options exist:

- Thorlabs GVS211/M scanner with 10 mm beam size (resulting in an excitation NA of 0.1) – these are standard Thorlabs items and have short lead times and are recommended for building a mesoSPIM quickly.
- Citizen Chiba GCM-2280-1500 scanner allowing a 15 mm beam (NA 0.15) – legacy option implemented in several mesoSPIM V4.
- Scanlab 14/1 dynaxis 3M scanner with a custom air-cooled galvo mount (NA 0.14) – this option has longer lead time and requires custom electronics and housing.

The galvo scanner creates the light-sheet by scanning the Gaussian input beam through a f=50 mm consumer single-lens reflex (DSLR) objective (Nikron AF-S 50 mm f/1.4 G) that was "hacked" into a telecentric scan lens (see **Supplementary Note: Optical design of the mesoSPIM setup**). Typically, the galvo scanners are driven with a symmetric sawtooth signal at f=99 or f=199.9 Hz. The amplitude of the galvo driver signal dictates the height of the light-sheet (**Supplementary Figure 12**). Each excitation path is located on a Thorlabs TB1515/M large-area translation stage (see **Supplementary Figure 9b**) to allow changing the distance between the detection axis and the excitation objectives to accommodate immersion cuvettes of varying sizes. In practice, the EL-16 tunable lenses in combination with a f = 50 mm scan lens have a tuning range in excess of >100 mm in air, so that all distance changes can be done by changing the driving signal to the ETLs.

**Immersion cuvettes:**

We typically keep the immersion medium in an quadratic glass or quartz macro-fluorescence cuvette which are 30×30×30 $mm^3$, 40×40×40 $mm^3$, or 50×50×50 $mm^3$ in size (Portmann Instruments or Hellma AG). The immersion cuvettes are mounted on plastic cuvette mounts milled from polyoxymethylene (POM). At the bottom, the cuvette mounts are attached to a KB25/M kinematic base (Thorlabs). This quick-detachable magnetic base with a ball and V-groove design allows quick exchange between immersion cuvettes filled with different immersion media (**Supplementary Figure 9e**). For example, for imaging in CLARITY, we fill the cuvette with an immersion oil (Cargille 50350, $n_D$=1.45) whereas for imaging in iDISCO-cleared and stained-samples, the cuvette is filled with dibenzyl ether ($n_D$=1.562). If a mesoSPIM is supposed to be used with many different immersion media within a day of imaging, we recommend having dedicated immersion cuvettes filled with each medium ready.

**Sample holders:**

The sample XYZ & rotation stages carry a KB25/M kinematic base (Thorlabs) as well. This allows quick exchange of samples – they can be "clicked" in and suspended below the stages within seconds. For mounting samples, we either use cuvettes or sample clamps: For example, CLARITY cleared whole-mouse brains are usually immersed in 10x20x45 $mm^3$ quartz cuvettes (Portmann Instruments) in a refractive-index matching solution (RIMS). The same approach is recommended for all samples that are swollen or fragile. Samples cleared with organic solvents (i.e. DISCO, iDISCO, or BABB) tend to shrink and harden in the clearing process and can be clamped in a holder (either milled from aluminum or 3D-printed). As immersion media such as BABB or DBE tend to dissolve plastic, we recommend using nylon (polyamide) screws and 3D-printed parts for this purpose – in our hands, these tend to be very resistant. An overview of the Pros and Cons of the different sample mounting strategies is shown in **Supplementary Figure 10**.

**Sample stages:**

The travel range of the sample stages (see **Supplementary Figure 9d**) is currently 44.5 mm × 100 mm × 44.5 mm (X/Y/Z, mesoSPIM V4). The stages are produced by Physik Instrumente, Germany: M-112K033 (for X & Z movement), L-406.40DG10 (Y movement), M-116.DG (sample rotation) and controlled by a C-884 controller. In the mesoSPIM Version 4, the sample XYZ & rotation stages are mounted on a massive aluminum block (150×65×300 $mm^3$) that has a central hole for the detection objective (termed the 'bird house'). As this aluminum block requires a large milling machine for manufacturing, it has been replaced by a gantry built from X95-profiles (Qioptiq) in mesoSPIM V5. Version 5 also features slightly larger travel ranges (52 mm × 52 mm × 102 mm) by using PI L-509 stages.

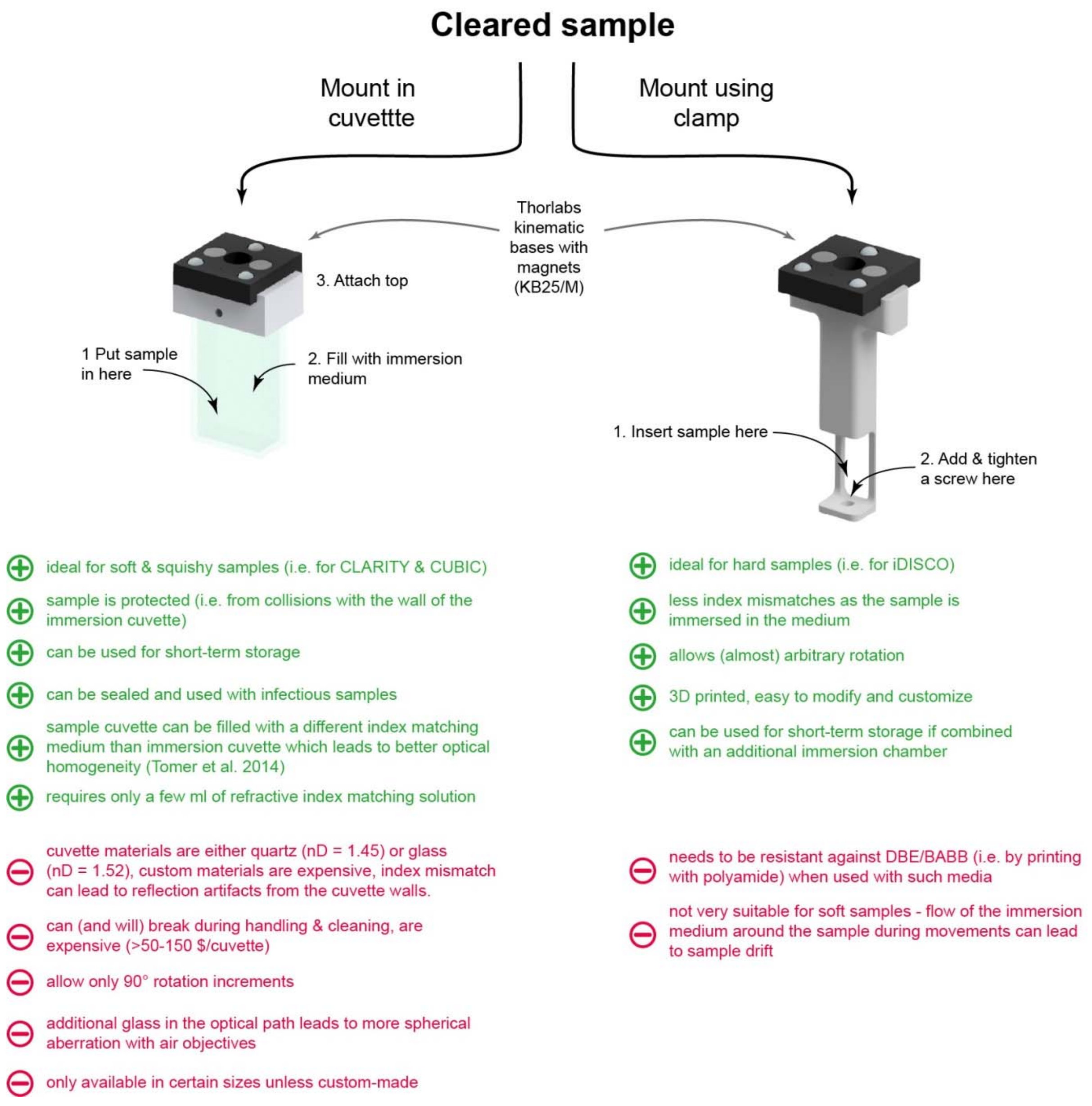

**Supplementary Figure 10: Comparison of mesoSPIM sample mounting approaches.** Samples are mounted using either an imaging cuvette (left) or a sample clamp (right), which both have a wide range of advantages and disadvantages. Beyond these standard sample holders, any custom holder can be fitted to a mesoSPIM as long as it interfaces with the Thorlabs KB25/M kinematic base with magnets.

**Detection path:**

The mesoSPIM detection path (**Supplementary Figure 9b**) consists of an Olympus MVX-10 macroscope with a MVPLAPO 1x objective, which allows FOVs of 2-20 mm in combination with a Hamamatsu Orca Flash 4.0 V3 camera. Due to its zoom system, the Olympus MVX-10 does not have a classical infinity-corrected space between objective and tube lens/zoom body that could be varied for focusing. Therefore, we opted to translate the entire detection path (MVX10 + filter wheel + camera assembly) for focusing by mounting it on a heavy-duty linear stage (PI M-406.4PD). The focusing stage carries an additional FLS-95 rail so that the detection system can be highly modular. The stage also aids in refocusing between different zoom settings as the MVX-10 has a large focus drift with magnification (> 150 µm for a change from 1× to 4×). The MVX-10 has a manual zoom wheel which we motorized by attaching a Robotis Dynamixel MX-28R servo to it. The MX-28R is computer-controlled via a serial RS-485 connection and has an internal rotation encoder with a resolution of 4096 steps/revolution, which we deemed to be adequate for the approximately 160 degrees of rotation necessary to change the zoom setting from 0.63× to 6.3×. For higher magnification, the Olympus MVPLAPO 2x objective can be used, but it has large distortion in the corners of the FOV. At the rear port of the MVX-10, we attached a Ludl 96A350 filter wheel with ten 32-mm filter positions. The filter wheel is controlled by a MAC-6000 controller which is connected via RS-232 to the imaging computer. The rear port of the filter wheel housing interfaces with a Olympus MVX-TLU tube lens and a MVX-TV1xC C-mount adapter to which the camera is attached. The camera is rotated by 90° around the detection axis because in an ALSM instrument, the read-out direction of the rolling shutter and the translation of the waist have to be parallel.

**Waveform generation and electronics:**

The control signals for the galvo scanners and tunable lenses are generated by a National Instruments (NI) PXI-6259 card housed in a NI PXIe-1073 chassis connected to the imaging computer by a MXI-bus. An additional PXI-6733 card controls the laser intensities (up to 8 lines) via the analog inputs of the laser engines. All waveforms are generated by the mesoSPIM-control software (see corresponding Supplementary note), buffered in advance and triggered by a master trigger routed to all NI cards. The camera is triggered via a counter output on the PXI-6259. A detailed overview of the electronic mesoSPIM components is given in **Supplementary Figure 11** whereas the timing and shape of the mesoSPIM waveforms is shown in **Supplementary Figure 12**.

**Installation instructions:**

Detailed instructions for setting up a mesoSPIM can be found on the mesoSPIM wiki: https//github.com/mesoSPIM/mesoSPIM-hardware-documentation.wiki

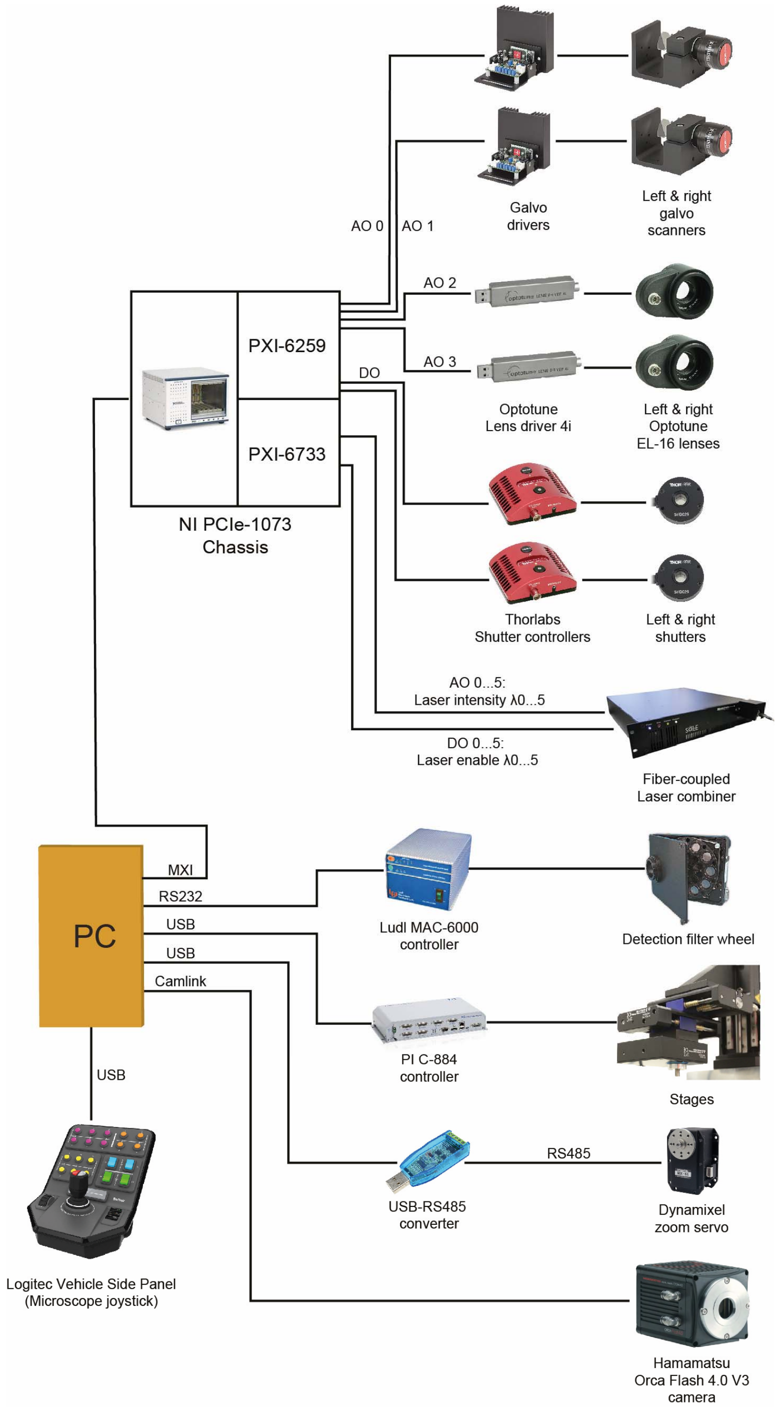

**Supplementary Figure 11: Block diagram of the mesoSPIM electronics.** The sCMOS camera, zoom servo, mechanical stages and filter wheel all have dedicated connections to the imaging computer (PC) whereas the waveform generation (ETL ramps, galvo signals, shutter control, and laser intensity control) is done via a National Instruments PCIe-1073 PXI chassis.

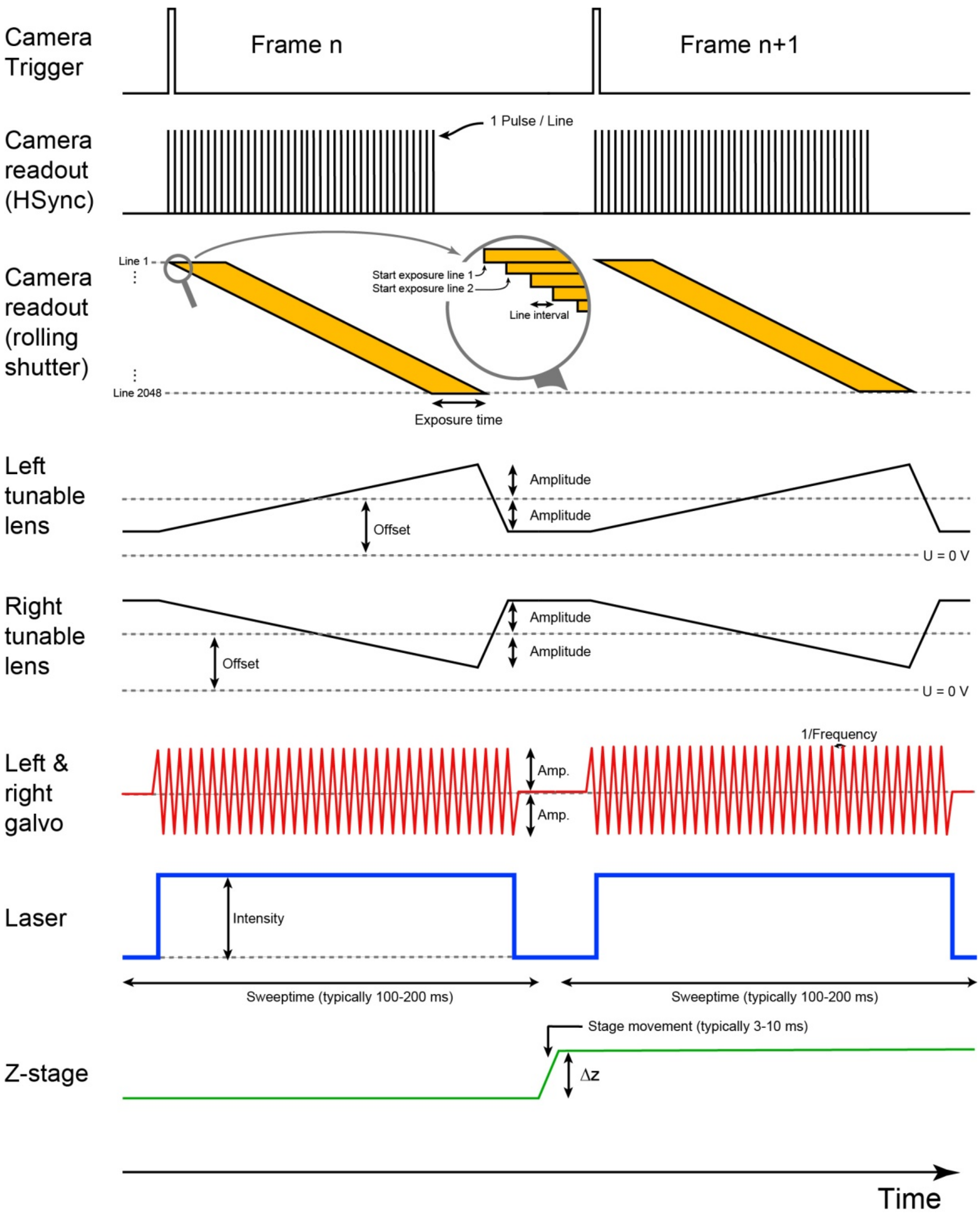

**Supplementary Figure 12: Timing diagram of the mesoSPIM waveforms.** The camera trigger starts the rolling shutter, which is synchronized to a sweep of the tunable lenses. At the same time, the light-sheet is created by rapidly scanning a Gaussian beam using the galvo scanners. While the exposure is running, the laser is set to a constant intensity and switched off between frames. During the same time, the camera frame is read out and the z-stage can advance.

**Supplementary Note: Microscope software: mesoSPIM-control**

A wide range of custom light-sheet microscopes (e.g. the openSPIM instrument) are controlled by Micromanager[58], an open source microscope control software based on ImageJ (imagej.net)[59]. As a significant fraction of early mesoSPIM hardware components were not supported by Micromanager, we opted to control the instrument with custom-written software in Python. The mesoSPIM-control software (https://github.com/mesoSPIM/mesoSPIM-control) software runs on Python >3.6 and was tested both on Windows 7 and Windows 10. It utilizes the PyQt5 bindings for Qt V5, a set of cross-platform C++ libraries that implement high-level APIs for accessing many GUI functions. In addition, we are using the QThread class of Qt5 for multithreading to allow the user interface to stay responsive while the microscope is acquiring images. The Camera Window and histogram controls are using the ImageView class of the pyqtgraph graphics and user interface library (www.pyqtgraph.org). Installation instructions are documented in the software wiki (https://github.com/mesoSPIM/mesoSPIM-control).

After startup, users are prompted to select a microscope configuration file. This file contains assignments of digital and analog output channels to microscope actuators (i.e. stages, galvo scanners, tunable lenses etc.) and configuration data for all major microscope components (i.e. filter assignments to filter wheel positions). The configuration files are user-editable and contain all parameters in the form of Python dictionaries. If necessary, the microscope then carries out referencing movements of the translation stages. The microscope software consists of three windows (**Supplementary Figure 13**): “Main Window”, “Camera Window” and the “Acquisition Manager”.

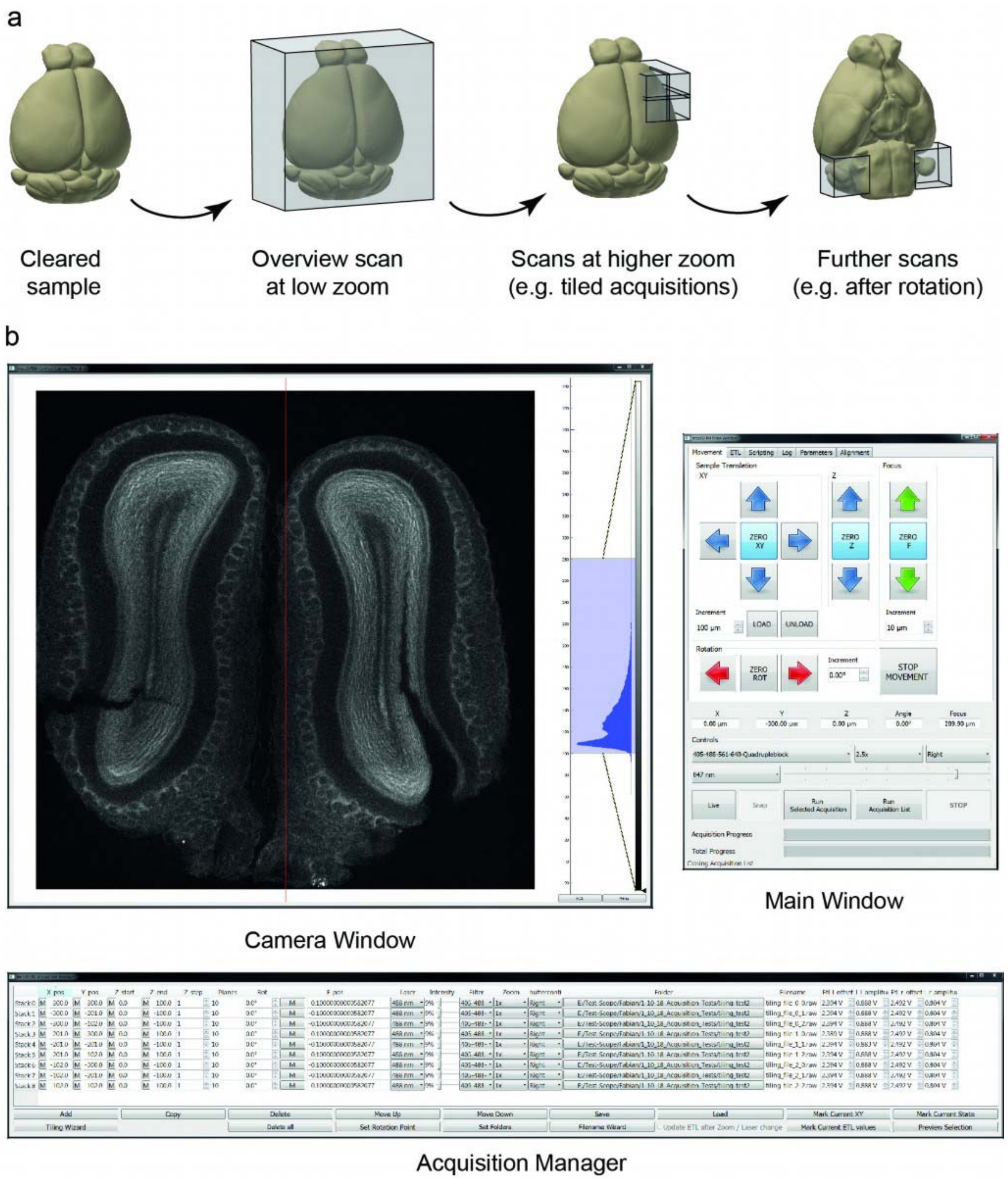

**Supplementary Figure 13: mesoSPIM-control.** a) Possible user-defined series of data acquisitions: Starting from a cleared sample, users typically perform overview scans to evaluate the quality of the labeling and clearing. Subregions (or the entire sample) can then be scanned with tiled acquisitions. By rotating the sample, multi-view datasets can be generated. b) The mesoSPIM-control user interface consists of three windows: A Main Window with microscope controls (i.e. for sample position, laser selection, laser intensity etc.), a Camera Window with histogram controls and an Acquisition Manager, a table-based tool to plan a series of z-stacks.

The “Main Window” contains all user controls for sample movement, microscope focus, and user-adjustable parameters such as emission filters and laser intensity. After clicking the “Live” button, the “Camera Window” displays images acquired with the current set of parameters without saving them. As most mesoSPIM users aim for the thinnest possible optical sections using the ASLM mode, the “Main Window” provides extensive control over ETL parameters in the ETL tab: Firstly, sets of predefined ETL parameters (“ETL configuration files”) for each combination of zoom, excitation wavelength, and light-sheet direction (left or right) can be saved and loaded as .csv-files. As ETL parameters (offset and amplitude) depend on the refractive index of the medium inside the immersion cuvette and its size, a dedicated ETL configuration file should be generated for each combination of clearing method and immersion cuvette the setup is supposed to be used with. In addition, individual clearing methods might require more fine-grained presets: As RIMS[60] refractive indices can vary with preparation and temperature, it is recommended for each user/project to have their individual ETL parameter file. In contrast, clearing methods based on organic solvents such as BABB or iDISCO tend to have much more uniform refractive indices across different batches of samples.

To optimize the effective light-sheet thickness, we recommend the following procedure: After setting up the desired zoom and excitation wavelength, users can deactivate the ASLM mode by toggling the “Amp=0” buttons in the ETL tab. This sets the ETL amplitude to zero and saves the current amplitude (which can be reloaded by de-toggling the button). Effectively, this turns the mesoSPIM in a standard light-sheet microscope with a scanned Gaussian beam (DSLM) for light-sheet generation. Users can then manually place the waist location (visible as the vertical stripe with the lowest signal or the least amount of sample features) in the center of the camera FOV (aided by crosshairs in the camera window). In samples with insufficient clearing, visually localizing this region can be a challenge. As soon as the waist is placed in the center of the FOV, users can switch the ASLM mode back on and optimize the ETL amplitude.

In the ideal case, the amplitude is just right for the ETL-actuated waist and the rolling shutter of the camera to track each other perfectly across the FOV of the microscope. If this is not the case, for example, when the ETL amplitude is too small or too large for a given zoom/FOV setting, the center of the image will display good axial confinement of the light-sheet whereas the left and right edges will appear more blurry or contain more sample features as the effective light-sheet thickness is larger in these regions (see **Supplementary Figure 5**). By observing the left and right edges of the image while changing the ETL amplitude, an optimum can be found.

After finding the optimum ETL parameters for a certain region of the sample, users can generate complex multidimensional acquisition sequences using the "Acquisition Manager" window, which contains a table of z-stacks. Each stack (or "acquisition") is represented by a row in this table and can have a user-defined startpoint (defined by its XYZ and rotation coordinate) and an endpoint at a different z-location. The software then automatically calculates the number of planes for a selected z-spacing. In addition, users can define the laser line, laser intensity, emission filter, zoom and shutter configuration (left/right/both light-sheets). In addition, a filename and a folder location can be specified. The tunable lens offset and amplitude values can be specified as well for each row. A series of "Mark" buttons allows the user to copy elements of the current microscope state (i.e. filter and laser intensity or ETL parameters) into a table row to simplify setting up acquisitions.

Tables for mosaic acquisitions can be generated using a "Tiling Wizard" which guides users through a series of steps (starting from the definition of a bounding box for the list of acquisitions). A preview button allows users to visually inspect a single selected acquisition in "Live mode" by moving the sample to the starting position and setting up the desired excitation wavelength, laser intensity, filter, shutter configuration and ETL parameters. This aids in checking whether the imaging parameters are correct for each individual stack. By selecting a table row and clicking the "Run selected acquisition" button in the main GUI, a single z-stack can be acquired. By clicking “Run Acquisition List”, the microscope acquires all acquisitions specified in the table from top to bottom. Acquisition manager tables can also be saved and reloaded, i.e. to apply the same tiling pattern to a different sample.

If the user specified a sample rotation between subsequent z-stacks, the microscope will move in X,Y, and Z to a safe rotation position, rotate the sample, move back, and continue with the next z-stack. This is necessary as large samples can collide with the immersion cuvette during rotation. The rotation position can be inspected and specified in the main user interface. By specifying multiple rotation positions, multi-view acquisitions can be generated (see **Supplementary Figure 25** ). Currently, the microscope saves acquisitions in a raw-format with an additional metadata file. A “Parameter” tab in the main GUI allows access to other microscope configuration parameters such as the delays, rise and fall times for the ETL ramps, camera exposure time and line interval. In addition, the Main Window has a “Alignment” tab that allows running the microscope in a mode that simplifies co-alignment of both light-sheets by interleaving the illumination directions frame-by-frame.

**Supplementary Note: Resolution measurements and microscope characterization**

To generate test-samples, we embedded Fluoresbrite YG Microspheres (1-µm diameter; Polysciences) in 1% agarose (Sigma-Aldrich 9012-36-6). For resolution measurements at low zoom settings (1×), we used 1:10000 dilution, for 4× 1:1000. We cured the agarose/bead mixture in molds made out of 10 ml syringes with cut-off tips. Using the plunger, removing the agarose from the mold was straightforward. The agarose cylinders were then transferred to a 20 ml RIMS[60] solution to begin index matching. After 24 hours, the samples were transferred into 10×20×45 $mm^3$ quartz cuvettes and the RIMS was exchanged. We noted that full index-matching could take up to a week. It is recommended to keep such test-samples for long time periods in the imaging cuvette to periodically check the microscope performance. To allow this, the cuvettes should be covered to avoid evaporation and can be placed in a fridge at 4°C. The RIMS was prepared at an index of nD=1.45 using a refractometer (Krüss DR301-95). We then placed the samples in the immersion cuvette filled with a fused silica immersion oil (Cargille 50350). This allowed resolution measurements under conditions for imaging CLARITY samples. We acquired z-stacks over a range of 200 µm at 1-µm z-spacing after manually optimizing the ETL parameters. The datasets were then converted to .tif and analyzed using the Jupyter notebook for mesoSPIM PSFanalysis (github.com/mesoSPIM/mesoSPIM-PSFanalysis), which are based on the PSF analysis code developed by Nick Sofroniew for the Thorlabs two-photon mesoscope (github.com/sofroniewn/psf). Briefly, after smoothing using a Gaussian filter, maxima (corresponding to beads) are detected by the scikit-image (https://scikit-image.org/ ) "peak_local_max" function. Maxima locations that are too close in 3D than a predefined bounding box (window size) in X x Y x Z (20 × 20 × 40 $\mu m^3$ at zoom 1×; and 5 × 5 × 20 $\mu m^3$ at zoom 4×) are discarded to allow for measurements on single beads. A Gaussian function is then fitted to the lateral and axial intensity profiles. For lateral PSFs, the X and Y full-width-at-half-maximum (FWHM) are averaged. The resulting data are shown in **Supplementary Figure 14 and 15**.

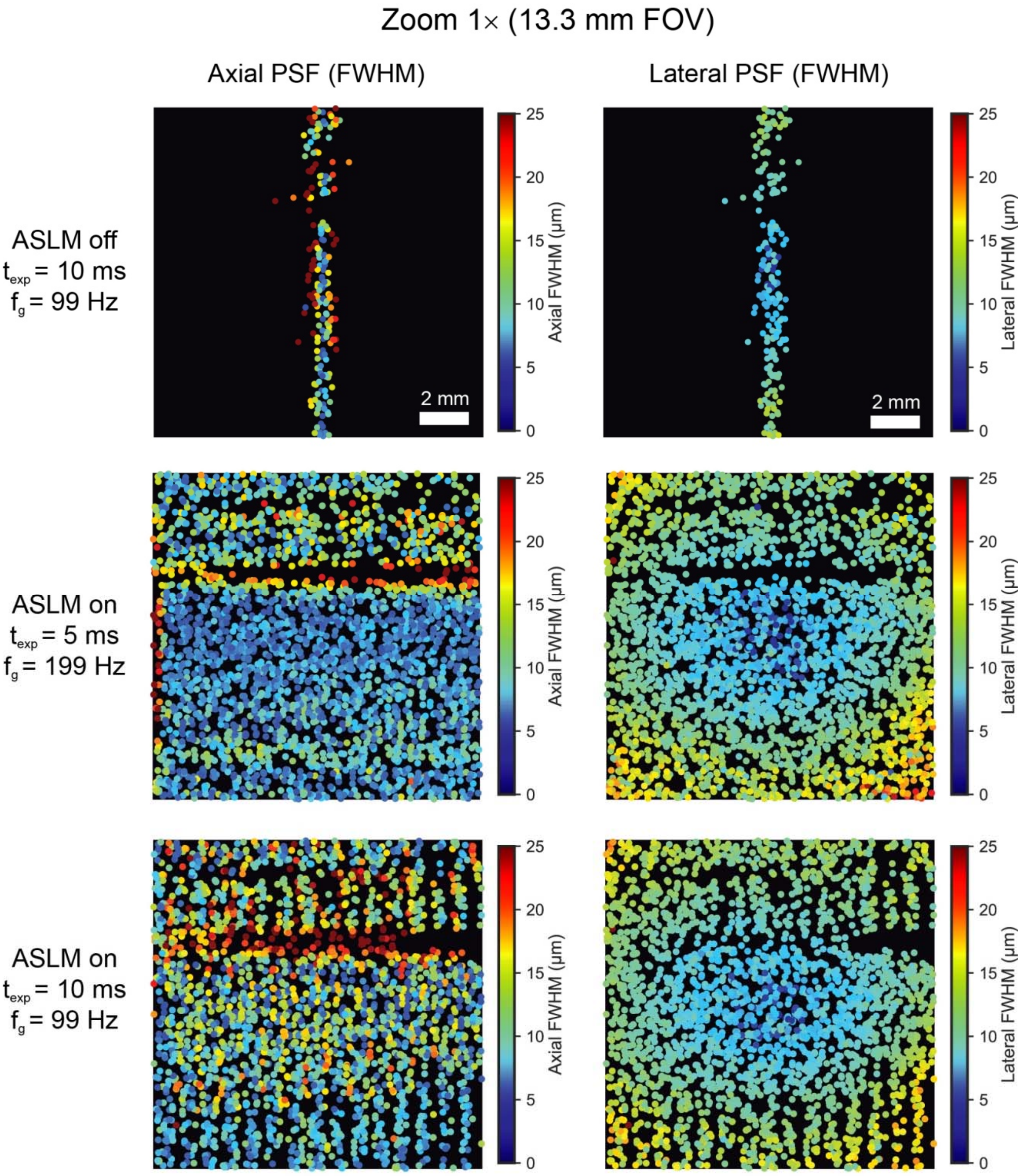

**Supplementary Figure 14: Axial and lateral resolution at low zoom (1×).** Color-coded FOV-maps of the axial (left) and lateral (right) point-spread-function (PSF) full-width at half-maximum (FWHM) at zoom 1×. With the ASLM mode switched off, beads were detected only along the light-sheet waist as the axial FWHM increased rapidly beyond the window size. The beads were diluted 1:10000 in 1% agarose and index matched to a final index of nD = 1.45. The excitation wavelength was 488 nm, the emission light was filtered using a 520/35 band pass filter. Cracks in the agarose block deteriorate the light-sheet confinement and lead to shadow regions, along which beads could not be fitted. Note that compared to the ASLM-off condition, running the ASLM mode at a galvo scanner frequency $\boldsymbol{f_g}$= 199 Hz and 5-ms exposure time allowed a uniform axial resolution <7 µm across the whole lateral FOV in the central region of the sample. As the Olympus MVX-10 was originally designed for visual use, an inside-out gradient of the lateral resolution is visible. In addition, the lateral resolution measurements are sampling-limited due to a pixel size of 6.55 µm. At lower $\boldsymbol{f_g}$, the pattern of detected beads shows stripes due to detoriation of the axial resolution, leading to fewer detected beads.

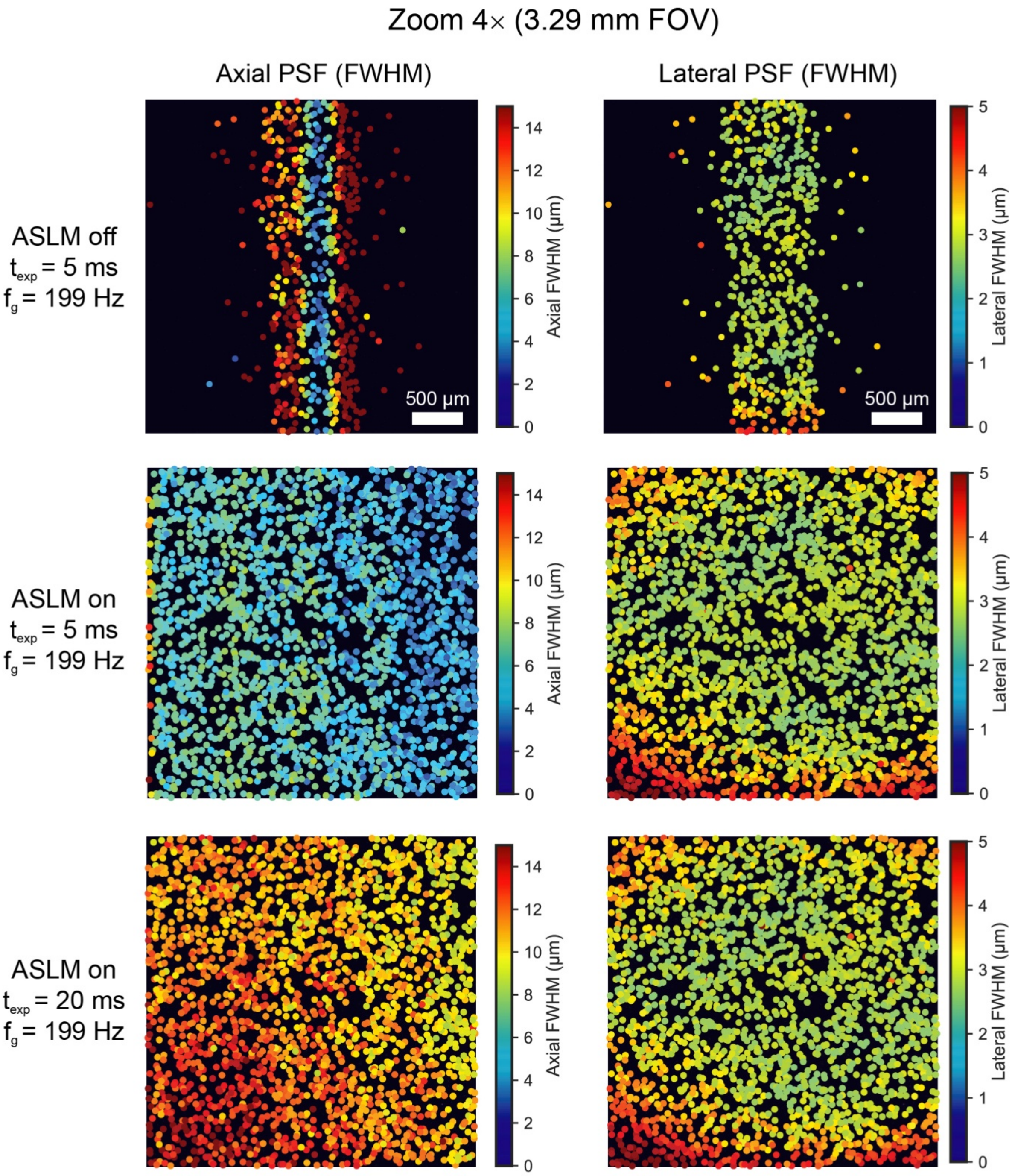

**Supplementary Figure 15: Axial and lateral resolution at high zoom (4×).** Color-coded FOV-maps of the axial (left) and lateral (right) point-spread-function (PSF) full-width at half-maximum (FWHM) at zoom 4x. Each dot represents a single detected 1 µm bead of which the axial and lateral intensity profiles were extracted and fitted with a Gaussian. To allow proper fitting of the PSF, only a single bead was allowed within a volume of 5 × 5 × 20 µm³. With the ASLM mode switched off, beads were detected only along the light-sheet waist as the axial FWHM increased rapidly beyond the window size. The beads were diluted 1:1000 in 1% agarose and index matched to a final index of nD = 1.45. The excitation wavelength was 488 nm, the emission light was filtered using a 520/35 band pass filter. Note that compared to the ASLM-off condition, running the ASLM mode at a galvo frequency $f_g$= 199 Hz and 5 ms exposure time allowed a uniform axial resolution of 4.097 ± 0.013 µm across the whole FOV. The axial resolution degrades with increasing exposure time as each active line on the sCMOS sensor then samples more axial variation of the passing light-sheet waist region. At this magnification, the lateral resolution is more uniform compared to 1× zoom, but shows an inside-out gradient as well.

By switching the ASLM mode off the resolution of the microscope could be estimated in a scanned Gaussian beam mode. At 488 nm, we estimated the FWHM of the non-scanned light-sheet as 3.514 ± 0.013 µm (n = 198 beads), which corresponds to a Rayleigh length of approximately 83 µm. When switching on the ASLM mode at zoom 4×, the average axial FWHM was 5.568 ± 0.027 µm (n = 2170 beads) across a 3.29-mm FOV (**Supplementary Figure 15**). To achieve these values, the lateral galvo sweep frequency was set to 199 Hz and the exposure time to 5 ms. While these values are higher than the non-ASLM z-PSF, it should be noted that even a Gaussian beam with a 5.5 µm FWHM waist has a Rayleigh range of 203 µm; the usable FOV of the light-sheet microscope ($\mathrm{FWHM}_{\mathrm{axial}} < \sqrt{2} \cdot \mathrm{FWHM}_{\mathrm{center}}$) was thus increased at least 8-fold. At lower zooms, it was also possible to achieve a considerable increase in uniformity of the axial PSF across the FOV: In the central region (13.29 mm in X × 1.3 mm in Y or 2048 × 200 pixels) of the 1× FOV, a resolution of 6.519 ± 0.074 µm (n = 322 beads) across a FOV of 13.29 mm was measured (**Supplementary Figure 14**). A Gaussian light-sheet with a 6.5 µm waist would have a Rayleigh range of 284 µm – enabling the ASLM mode thus corresponds to a 23.4-fold increase in FOV. However, such resolution measurements at low zoom suffer from inhomogeneities and cracks in the agarose, which make generation of stable test samples for large FOVs difficult.

The effective detection NA of the MVX-10/MVPLAPO 1x combination varies with zoom and can be estimated as NA = 0.075 at 1× and 0.22 at 4× (Olympus, personal communication). Based on these values, the diffraction-limited lateral resolution would be 3.5 µm and 1.18 µm, respectively. With 6.55 µm/pixel and 1.6 µm/pixel sampling, the mesoSPIM is undersampled at both zooms which results in much larger measured resolution values. In addition, at both zooms, a radial gradient of the lateral FWHM values is visible.

With longer and longer exposure times at constant ETL sweeptime (see **Supplementary Figure 12**), the effective light-sheet thickness (or axial FWHM) across the FOV increases: As each line of the detector collects photons for a longer time, it samples a wider stripe of the axial light-sheet profile. This means that in practice, a trade-off has to be found between exposure time (collecting sufficient signal) and light-sheet thickness. For example, for imaging antibody-stained samples in BABB (**Fig. 1e**; **Supplementary Figure 24**) or samples

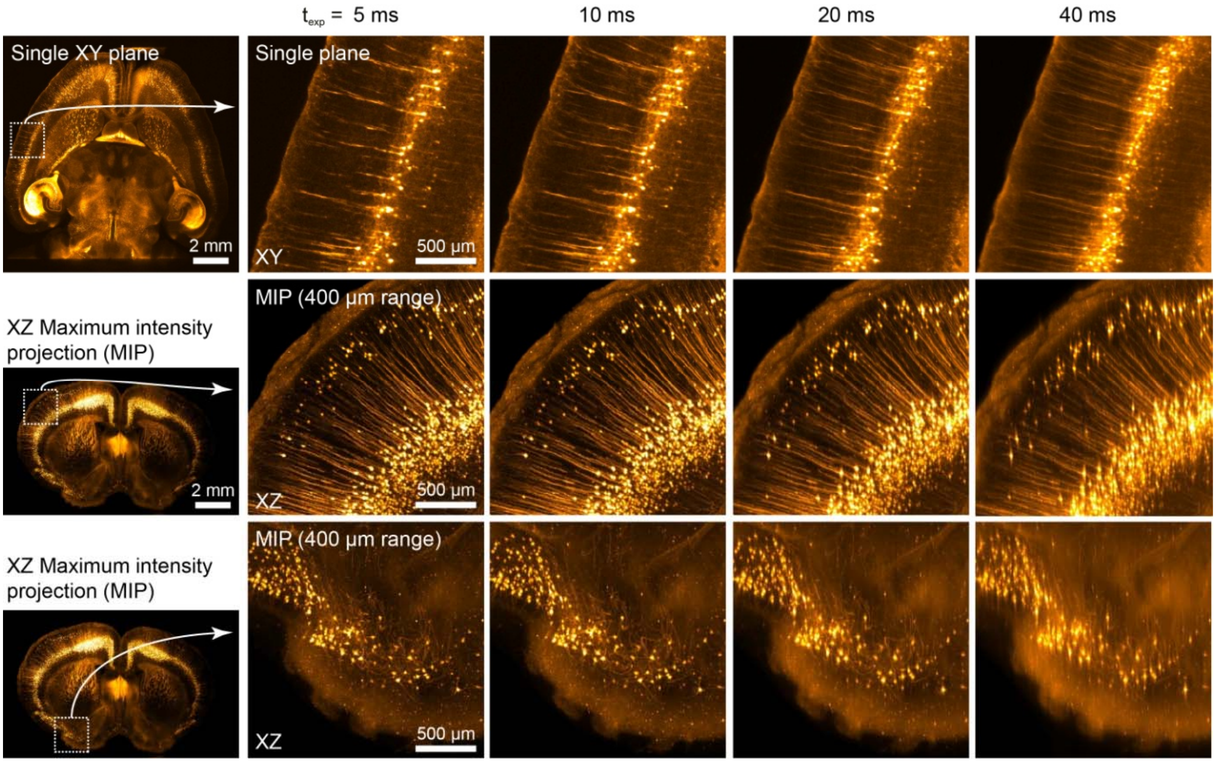

**Supplementary Figure 16: In ASLM, axial resolution depends on exposure time at low zoom (1x).** Comparison of example stacks taken in a CLARITY-cleared Thy1-YFP whole mouse brain. The galvo frequency $f_g$ was set to 199 Hz and the sweeptime to 200 ms. Increasing the exposure time yielded worsening of axial resolution. Note that despite the loss of axial resolution, it is uniform across the FOV; even 40-ms exposure time (which is 1/5 of the sweeptime) leads to a better axial resolution at the edges of the FOV compared to switching off the ASLM mode entirely (compare to Supplementary Figure 4). As the microscope collects more light with increasing exposure time, users have to select a tradeoff between exposure time and axial resolution depending on their requirements.

cleared and stained with iDISCO (**Supplementary Figure 27**), exposure times could be reduced to 10 ms or even 5 ms because the secondary amplification in the staining procedure yields high signal levels. When imaging endogenous fluorescence, however (**Fig. 1b**;**Supplementary Figure 18-20**), typical exposure times were 20 ms. To demonstrate this trade-off, we imaged the same CLARITY-cleared Thy1-H-mouse brain under a variety of imaging conditions: At zoom 1x (**Supplementary Figure 16**), increasing the exposure time from 5 to 40 ms led to a drastic reduction in axial resolution. At 4x magnification

(**Supplementary Figure 17**), the loss in axial resolution was less severe. In these experiments, the sweeptime was kept constant at 200 ms which led to an effective frame rate of 4.8 Hz.

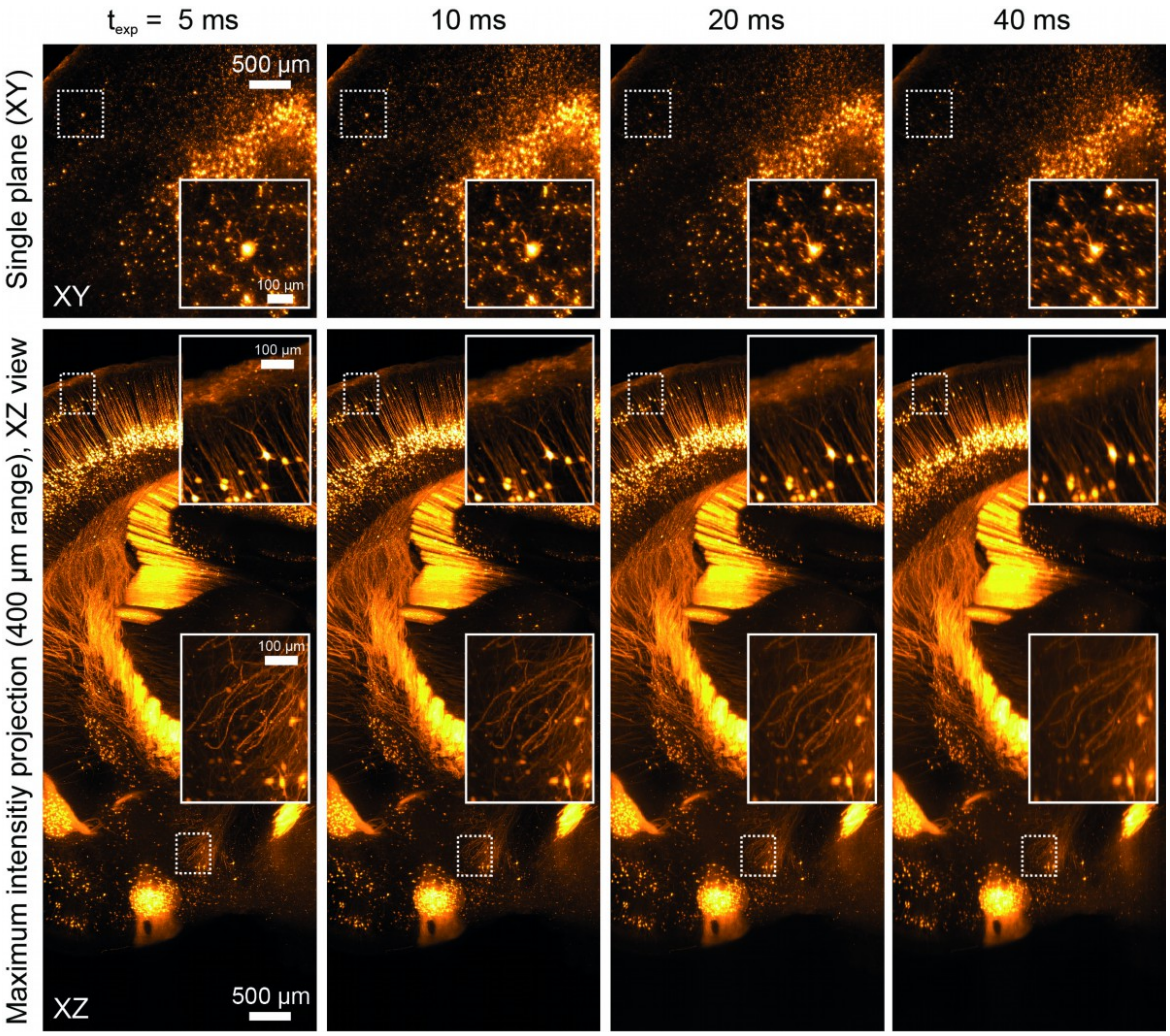

**Supplementary Figure 17: In ASLM, axial resolution depends on exposure time at high zoom (4x).** Comparison of example stacks taken in a CLARITY-cleared Thy1-YFP whole mouse brain. The galvo frequency $f_g$ was set to 199 Hz and the sweeptime to 200 ms. Increasing the exposure time yielded worsening of axial resolution. Note that despite the loss of axial resolution, it is uniform across the FOV; even 40-ms exposure time (which is 1/5 of the sweeptime) leads to better axial resolution at the edges of the FOV compared to switching off the ASLM mode entirely (compare to Supplementary Figure 4).

**Supplementary Note: Imaging examples: Compatibility with different clearing techniques**

A key advantage of the modular and open design of the mesoSPIM is the compatibility with all major clearing techniques. As different clearing techniques require immersion media with a wide range of refractive indices and different physical properties, typical light-sheet microscopes have to be modified extensively to allow for imaging across a wide range of media. For example, many CLARITY immersion media such as RIMS[60] are not very homogenous in refractive index (evident by visible striae in the medium). This is not surprising given that the RIMS by Yang et al. is based on Histodenz, which is originally a gradient density medium. Other media such as BABB (benzyl alcohol-benzyl benzoate) can lead to damage of the front lens of microscope objectives which is why very few objectives capable of imaging in BABB exist. An overview of all tested clearing techniques is given in **Supplementary Table 2**. As a reference, **Supplementary Table 3** lists all mouse lines used in this study. In addition, **Supplementary Table 4** provides an overview of the imaging parameters.

| Clearing technique | Sample | Figure | Supplementary Video |
|---|---|---|---|
| CLARITY (passive) | Mouse brain (VIPCre-tdTomato) | Figure 1b, Supplementary Figure 2 | Supplementary Video 1 |
| CLARITY (passive) | Mouse brain (TPH2Cre-tdTomato) | Figure 1c-e | Supplementary Videos 4 - 6 |
| CLARITY (passive) | Mouse brain (Rbp4Cre-YCX2.60) | Supplementary Figures 18-20 | Supplementary Video 7 |
| CLARITY (active) | Mouse brain (Thy1-YFP) | Supplementary Figures 3-6, 16-17 | Supplementary Videos 2 - 3 |
| X-CLARITY (active) | Mouse brain (GlyT2-EGFP) | Supplementary Figure 21 | Supplementary Video 8 |
| CUBIC-X | Mouse brain (Ntsr1Cre-tdTomato) | Supplementary Figures 22-23 | |
| BABB | Chicken embryo (Neurofilament) | Figure 1f, Supplementary Figures 24-25 | Supplementary Video 9 |
| BABB | Drosophila melanogaster | Supplementary Figure 26 | Supplementary Video 10 |
| iDISCO | Mouse brain (IgG) | Supplementary Figures 27-28 | Supplementary Video 11 |
| iDISCO | Rat brain (To-Pro) | Supplementary Figure 29 | |
| MASH (iDISCO/ECi) | Human cortex | Supplementary Figure 30 | Supplementary Video 12 - 13 |

**Supplementary Table 2: Overview of clearing techniques tested with the mesoSPIM.** References to the figures and supplementary videos are provided.

**The mesoSPIM is compatible with hydrogel-based clearing techniques such as active and passive CLARITY:**

The CLARITY[61] family of clearing techniques is based on crosslinking proteins in a hydrogel and using sodium dodecyl sulfate (SDS) as detergent to remove lipids from the sample. Lipid removal can be done 'actively', using electrophoretic assistance in a custom chamber[61], or using 'passive' or flow-assisted immersion in SDS[10,62]. Samples cleared with CLARITY tend to be slightly expanded after the index-matching step in the refractive index matching solution (RIMS) we employed (based on the Histodenz approach by Yang et al.[60]). As noted by Tomer et al[10], the refractive index of cleared samples is close to the index of quartz, which means that samples can be placed in quartz cuvettes for imaging. In addition, as many immersion media for CLARITY samples such as glycerol-water solutions and RIMS tend to be inhomogeneous in refractive index, placing the cuvette in an imaging chamber filled with an immersion oil at the index of quartz/fused silica (such as Cargille 50350) means that the light-sheet has to pass through a minimum of RIMS, which reduces the impact of local index variations in the quality of the light-sheet. For imaging CLARITY-cleared samples, we adopted the same technique and immersed whole mouse brains in a 10×20×45 $mm^3$ quartz cuvette filled with RIMS and then immersed this imaging cuvette in turn in an immersion cuvette (typically a 40x40x45 $mm^3$ quartz macro fluorescence cuvette made by Portmann Instruments). As CLARITY-cleared samples tend to float, we glued a small weight to the brain (typically a M4 nut) using quick glue.

To showcase the imaging quality achievable with the mesoSPIM in a mouse brain processed using active CLARITY, we imaged an Thy-1 YFP sample[63] (**Supplementary Figure 3-5**). In this sample, the mesoSPIM can resolve single neurons in sparse subregions at 1x zoom, which corresponds to a 13.29-mm FOV and 6.55-µm pixel size (**Supplementary Figure 3-4**, **Supplementary Video 2**). At higher magnification (zoom 4x, 3.3-mm FOV, 1.6-µm pixel size) axons can be imaged and are well resolved in the XZ plane (**Supplementary Figure 4**,

**Supplementary Video 3**). The preparation of this sample is described in a dedicated supplementary note (**Supplementary Note on Sample Preparation**).

A simplified CLARITY protocol utilizes either passive immersion or pumping of SDS through the clearing chamber without electrophoresis[10]. This "passive" CLARITY approach takes much longer (up to 4-6 weeks for a whole mouse brain), but many samples can be processed in parallel. As an example, we processed and imaged a whole mouse brain expressing tdTomato in VIP neurons, a subclass of inhibitory interneurons (**Fig. 1b**). Owing to the sparse expression pattern of this mouse line, single neurons can be visualized across the whole brain in the axial direction even at low magnifications such as 0.8x if the ASLM mode of the mesoSPIM is engaged (**Fig. 1b**, **Supplementary Figure 2**, **Supplementary Video 1**).

In addition, we applied the passive CLARITY protocol to a mouse brain expressing tdTomato in serotonergic neurons driven by a TPH2Cre line (Tryptophan Hydroxylase 2). In this sample, expression is not restricted to structures such as the Raphe nuclei, but widespread (**Fig. 1c,d, Supplementary Video 4**) – for example, both the hippocampi and the optic nerves between the optical chiasm and the lateral geniculate nuclei are strongly labeled. In addition, a small subset of Purkinje neurons showed labeling in dendrites, soma and axons **(Fig. 1d,e, Supplementary Videos 5 & 6).**

A key advantage of CLARITY, compared to clearing techniques that quench fluorescence within a few days (such as 3DISCO[7]), is that no additional staining is required to visualize expression patterns. For example, in two-photon microscopy, genetically encoded calcium indicators are often used to measure neuronal activity[64]. In such experiments, it is of considerable interest to link the functional response properties of neurons imaged *in vivo* to their anatomical identity and projection profiles. A key step in a cleared whole-mount sample is to achieve sufficient resolution of indicator expression so that single neurons can be identified. To demonstrate this, we processed mouse brains expressing the calcium indicator Yellow Cameleon YCX2.60[65] in excitatory neurons in layer 5 (L5) of the neocortex (Rbp4Cre-

YCX2.60) using passive CLARITY (details on the sample processing are provided in Supplementary Note 9). YCX2.60 is a calcium indicator based on fluorescence resonance energy transfer (FRET) and contains a ECFP/EYFP pair. For anatomical imaging, we excited EYFP at 515 nm and recorded background autofluorescence with 647-nm excitation (**Supplementary Figure 18**).

Overview images at low magnification (zoom 1×) reveal that in this mouse line, a wide range of long-range projections are labeled (**Supplementary Figure 18a-c**). In addition, in certain subregions, for example in the hippocampus, single neurons and their dendrites are readily visible in the XZ **(Supplementary Figure 18d)**. At 4× magnification, it is possible to detect single neurons in L5 in both the XY and XZ plane (**Supplementary Figure 19a-c**). In addition, the dense neuropil labeling by the calcium indicator leads to non-labeled cells visible due to counterstaining (**Supplementary Figure 19d**). As the ability to visualize such non-expressing cells is critically dependent on the effective light-sheet thickness, ETL parameters can be tuned in the sample by optimizing the contrast of these negatively stained cells. In addition, it is possible to visualize singe neurons and their dendrites in deeper regions of the brain such as the hippocampus (**Supplementary Figure 20a,b**). Interestingly, the Rbp4Cre line shows a sparse expression confined to the hippocampal region CA2 as well as strong expression in granule cells of the dentate gyrus (DG) and their axons forming the mossy fiber pathway (MF). Most importantly, the ASLM mode allows visualizing the same structures in the XZ plane with ease (**Supplementary Figure 20c, Supplementary Video 7**).

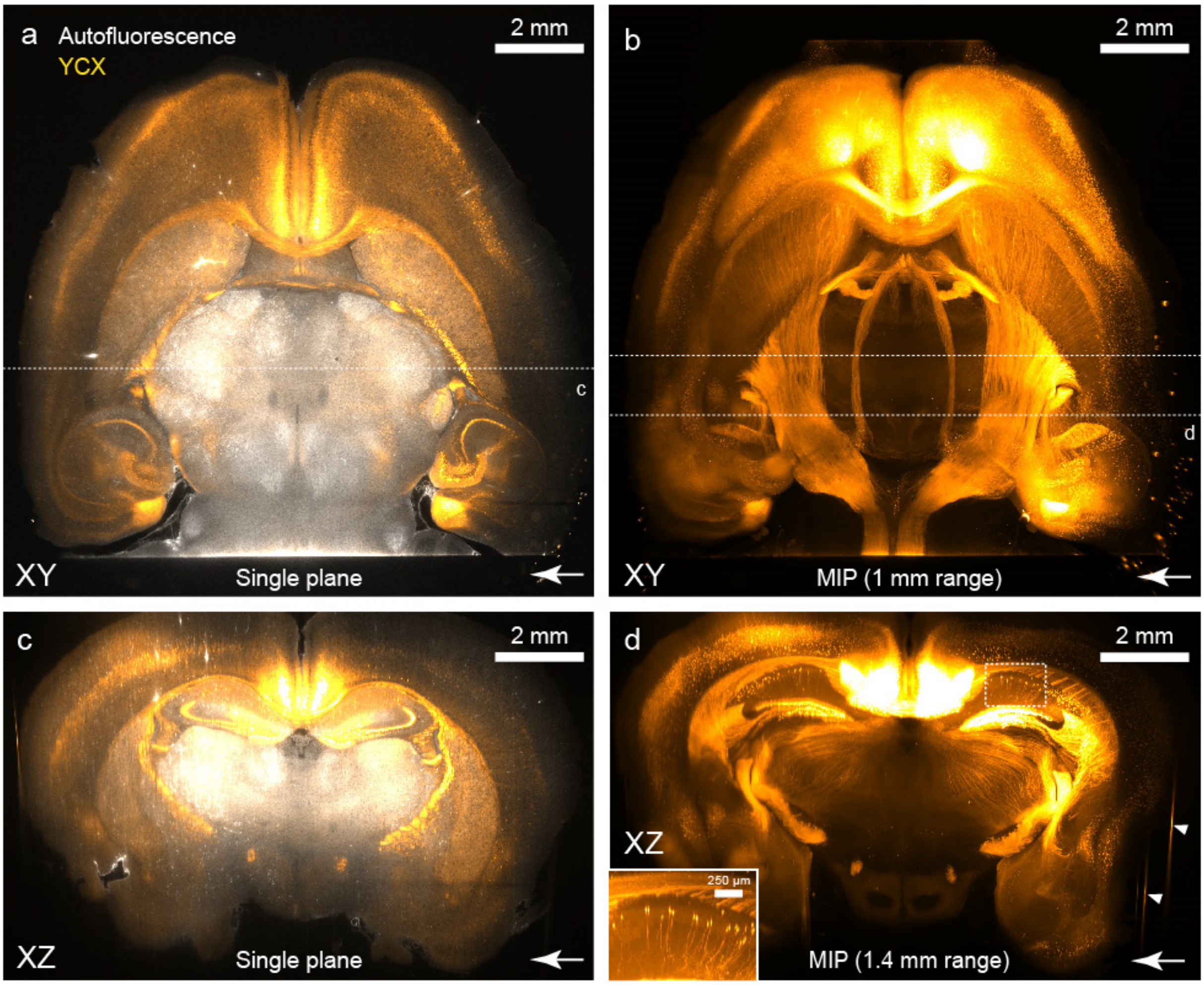

**Supplementary Figure 18: Low-resolution overview images (Zoom 1x) of a passive CLARITY-cleared mouse brain.** a) Single horizontal plane taken from an overview dual-color stack of a mouse expressing the calcium indicator Yellow Cameleon X 2.60 (YCX) in excitatory neurons in layer 5 (Rbp4Cre-YCX2.60) of the neocortex. b) Maximum intensity projection (MIP) over the basal half-section of the imaging volume in a). Long-range projections from the cortex to the spinal cord are visible. c) Single resliced plane (XZ view) from the dataset (location indicated in a). d) MIP over a range of 1.4 mm. Despite the large pixel size (6.55 µm), dendrites from YCX-expressing neurons in the hippocampus are visible in the XZ view (inset). The bottom right arrow indicates the light-sheet direction (illumination from the right side). Degradation of the light-sheet inside the tissue leads to the lower resolution in the left hemisphere. The arrow heads indicate Z-streaks caused by bubbles on the surface of the sample which scatter the excitation laser light.

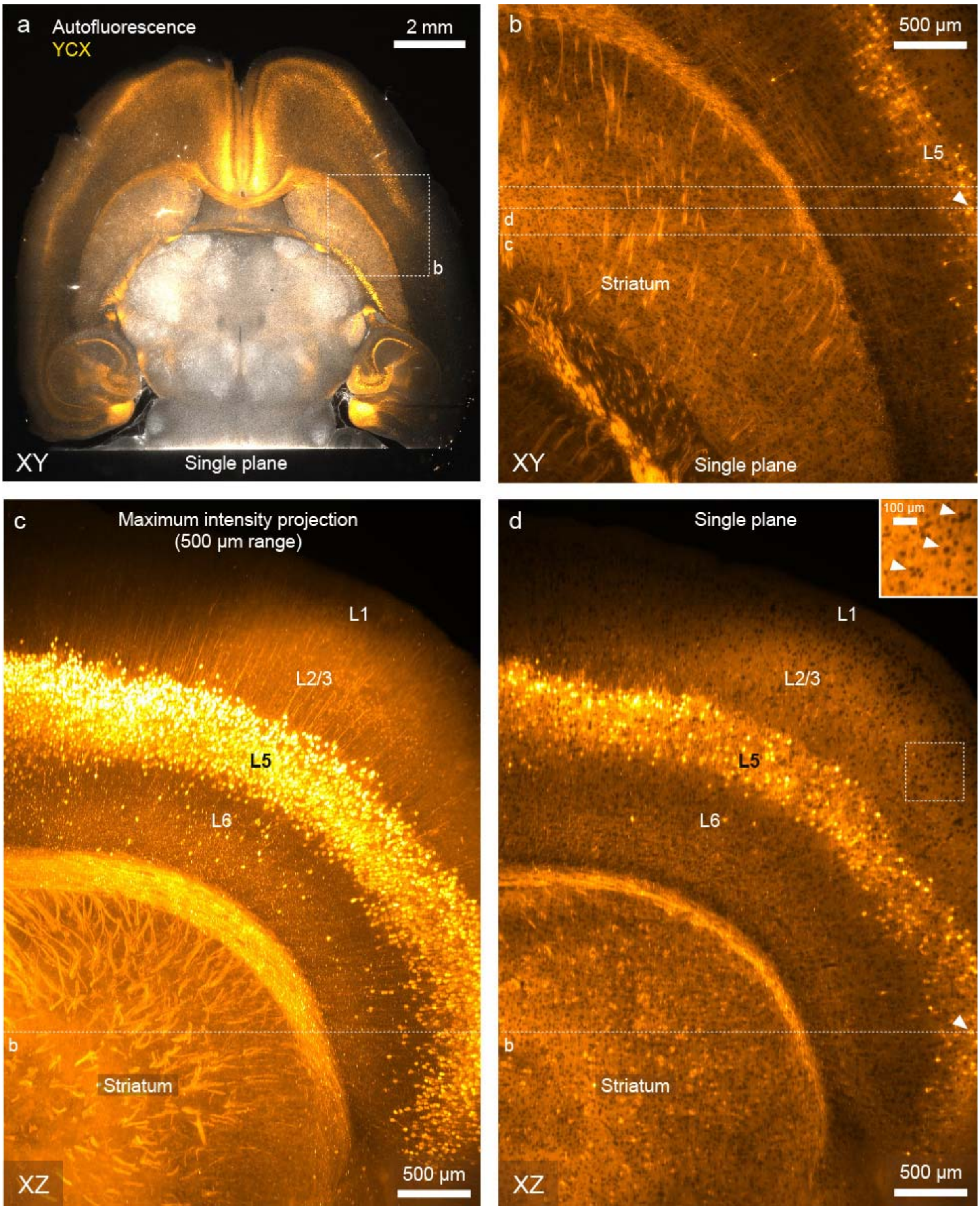

**Supplementary Figure 19: Images of a passive CLARITY-cleared mouse brain at higher resolution (Zoom 4×).** A mouse brain expressing the calcium indicator Yellow Cameleon X 2.60 (Rbp4Cre-YCX2.60) was cleared using passive CLARITY. a) Single plane taken from an overview dual-color stack. b) Single plane acquired at 4x zoom of the subregion shown in a). The arrow head indicates the same cell in subpanels b) and d) c) MIP in the XZ-plane (reslice) of the subvolume indicated in b). d) Single plane (location indicated in b) of the resliced dataset. The expression occurs predominantly in cortical layer 5 (L5). Long-range projections are visible in the striatum. A wide variety of counterstained cell bodies is visible throughout the volume and can be identified even in the XZ plane (inset).

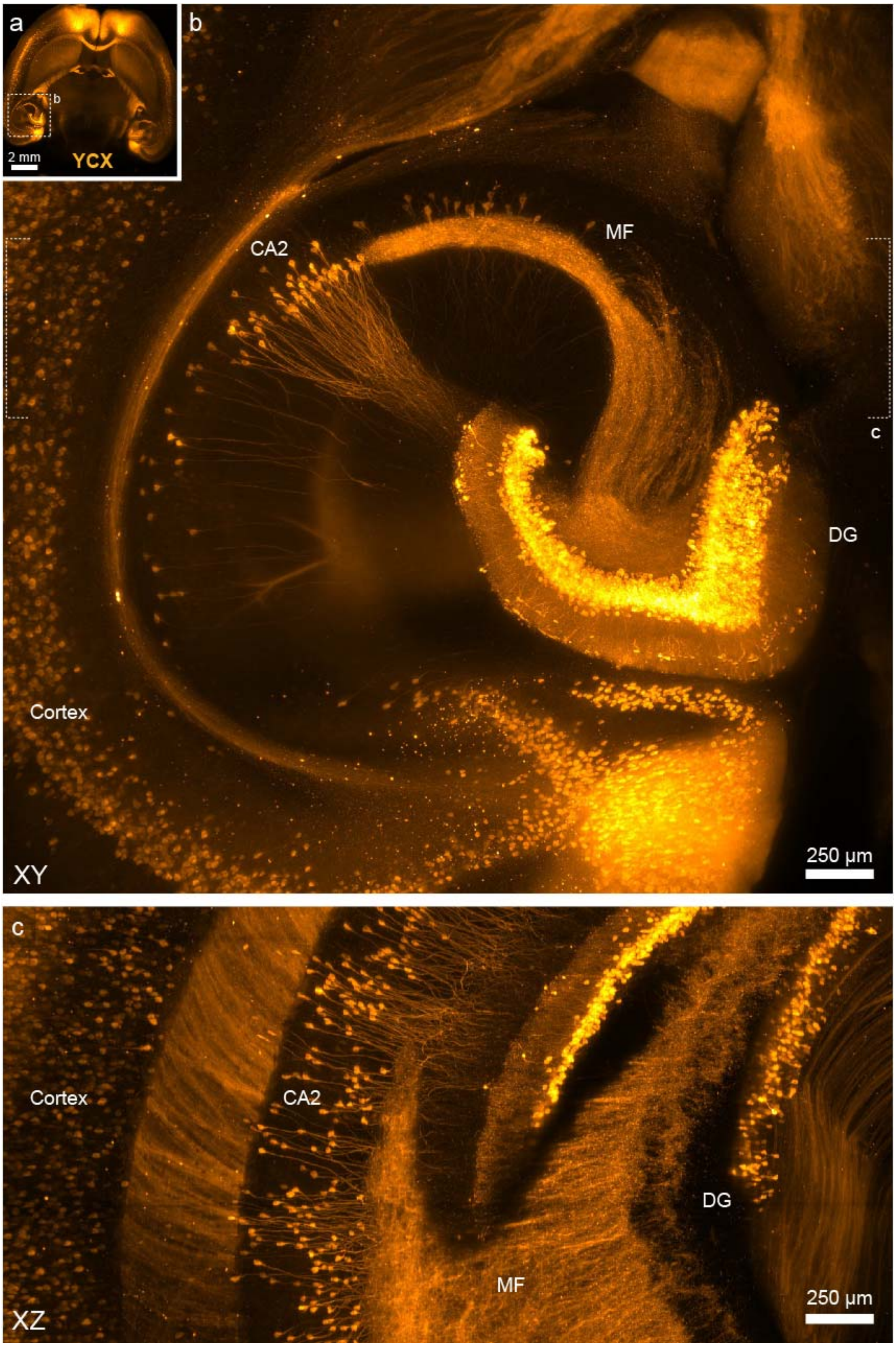

**Supplementary Figure 20: Multi-scale imaging with subcellular resolution (Zoom 4x) in a mouse brain cleared with a passive CLARITY protocol.** a) Overview image of a mouse brain expressing the calcium indicator Yellow Cameleon X 2.60 (Rbp4Cre-YCX2.60). b) MIP of a 600-µm thick virtual slice. In addition to the cortical expression, this mouse line shows sparse labeling in CA2 region of the hippocampus and strong labeling of granule cells in the dentate gyrus (DG). Furthermore, the mossy fiber pathway (MF) is labeled. c) MIP of a subvolume (indicated in subpanel a) of b). Single CA2 neurons and their dendrites are well resolved in the XZ plane.

**The mesoSPIM is capable of imaging a whole mouse central nervous system**

A key advantage of the mesoSPIM compared to existing commercial setups is that much larger cleared samples can be accommodated. To demonstrate this capability, we dissected a whole central nervous system (CNS) from a GlyT2-EGFP mouse[66] which expresses EGFP in glycinergic neurons in the spinal cord, brainstem, cerebellum, and thalamus. We processed the sample using the X-CLARITY clearing machine (Bucher Biotec AG). For mounting, we transferred the sample in RIMS in a custom 10×20×120 $mm^3$ quartz cuvette (Portmann Instruments) filled with RIMS and immersed it in a 40×40×120 $mm^3$ quartz cuvette filled with an immersion oil at nD = 1.45 (Cargille 50350). The large travel range of the sample stages (44.5 × 44.5 × 100 $mm^3$) then allowed scanning the entire sample without remounting or cutting (**Supplementary Figure 21**). Using this approach, single interneurons expressing the glycine transporter 2 (GlyT2) could be visualized in the spinal cord and hindbrain (**Supplementary Video 8**).

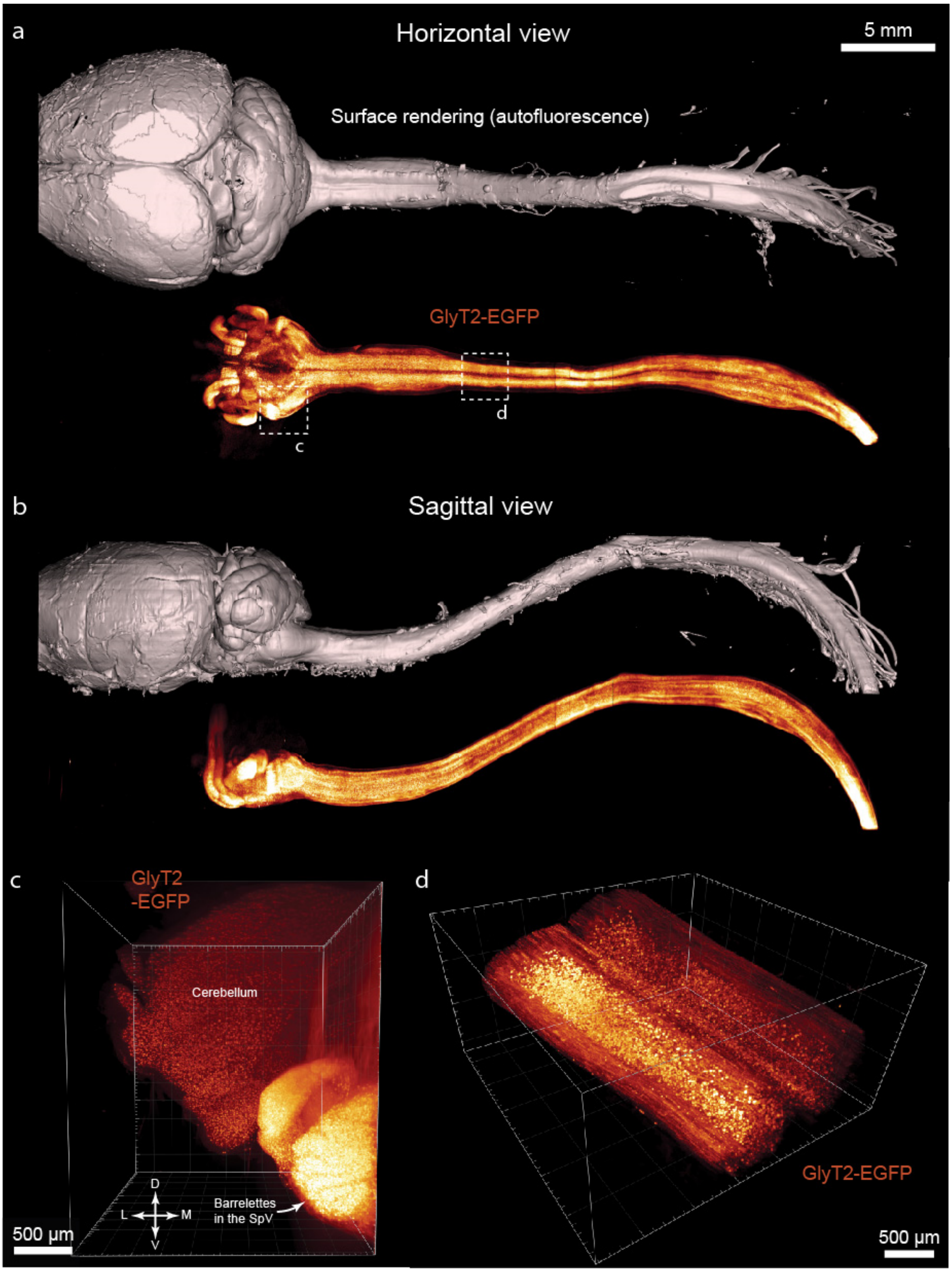

**Supplementary Figure 21: Whole-CNS imaging with the mesoSPIM.** a) A whole central nervous system was dissected from a Glycine Transporter-2 EGFP (GlyT2-EGFP) mouse and cleared using the X-CLARITY protocol. Shown is a stitched overview dataset taken at 1x zoom (13.29-mm FOV). As GlyT2 expression is restricted to the hindbrain and spinal cord, the forebrain does not show much signal. b) Sagittal view of the data in a). c) Volume rendering of a subvolume acquired using 4x magnification in the ventral hindbrain and cerebellum. Barrelettes in the trigeminal nucleus (SpV) are visible. d) Volume rendering of a volume acquired at 4x magnification in the spinal cord. Individual Glyt2-positive neurons are visible.

### The mesoSPIM is compatible with CUBIC-cleared samples

Apart from CLARITY, another family of clearing methods that retain endogenous fluorescence is a combination of "clear, unobstructed brain imaging cocktails and computational analysis" based on amino alcohols (CUBIC[5]). To demonstrate the compatibility of the mesoSPIM with CUBIC-cleared samples, we processed a Ntsr1Cre-tdTomato mouse brain with the CUBIC-X protocol[67], a recent improvement to the original CUBIC which can also be tuned for sample expansion. After clearing, the sample was transferred to a 10×20×45 $mm^3$ quartz cuvette similar to the ones used for imaging CLARITY samples. The imaging cuvette was then filled with the CUBIC-X2 index matching medium, a combination of 5% (w/v) imidazole and 55% (w/v) antipyrine. To keep the sample from floating to the surface of the imaging medium, a 10×20×10 $mm^3$ plug made out of 2% agarose was then inserted above the sample. The imaging cuvette was then submerged in a 40×40×45 $mm^3$ quartz cuvette filled with the same CUBIC-X2 solution. While the original approach of embedding the samples in 2% agarose for imaging can also be utilized with a mesoSPIM, we noted that the approach using cuvettes tends to be much simpler and safer as the sample was very fragile. As the Ntsr1Cre-tdTomato line has an expression pattern restricted to a subset of layer 6 (L6) excitatory neurons, overview images at low zoom show strong labeling in this region and in thalamus (**Supplementary Figure 22**). At 4x magnification, it is possible to visualize commissural axons in the hindbrain (**Supplementary Figure 23**).

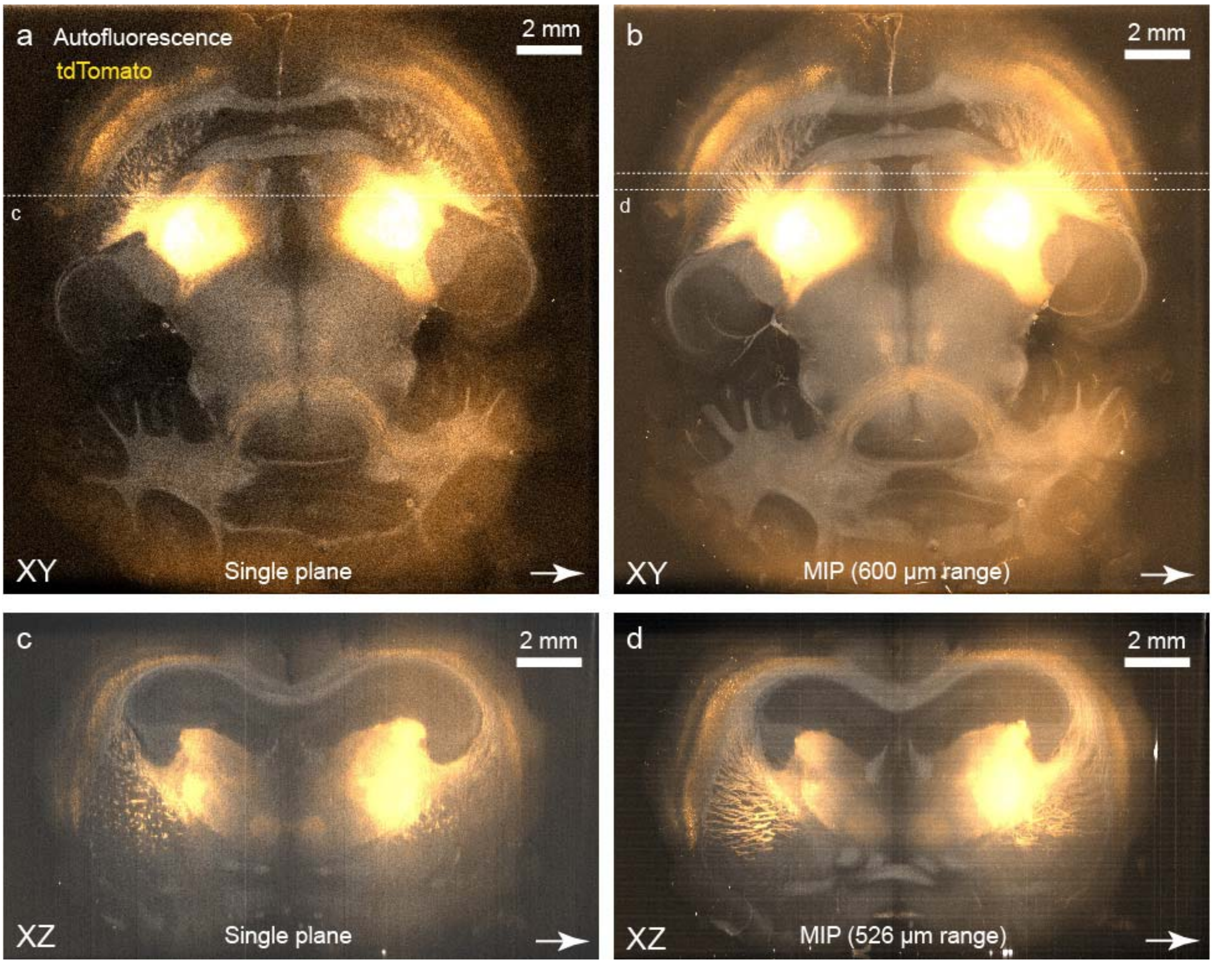

**Supplementary Figure 22: Low-resolution overview images (Zoom 0.63x) of a cleared mouse brain cleared using CUBIC-X.** A mouse expressing tdTomato in a subset of layer 6 (L6) neurons (Ntsr1-Cre;CAG-tdTomato) was cleared using the CUBIC-X protocol and imaged with the mesoSPIM. a) Single horizontal plane taken from an overview dual-color stack. b) Maximum intensity projection (MIP) over a 600-µm range of the imaging volume in a). Long-range projections from the cortex to the thalmus are visible. c) Single resliced plane (XZ view) from the dataset (location indicated in a). D) MIP over a range of 526 µm. The bottom right arrow indicates the light-sheet direction (illumination from the left side). Degradation of the light-sheet inside the tissue leads to the lower image resolution on the right half of the brain. Due to low signal levels, the images are very noisy and stripes from single camera pixels are visible.

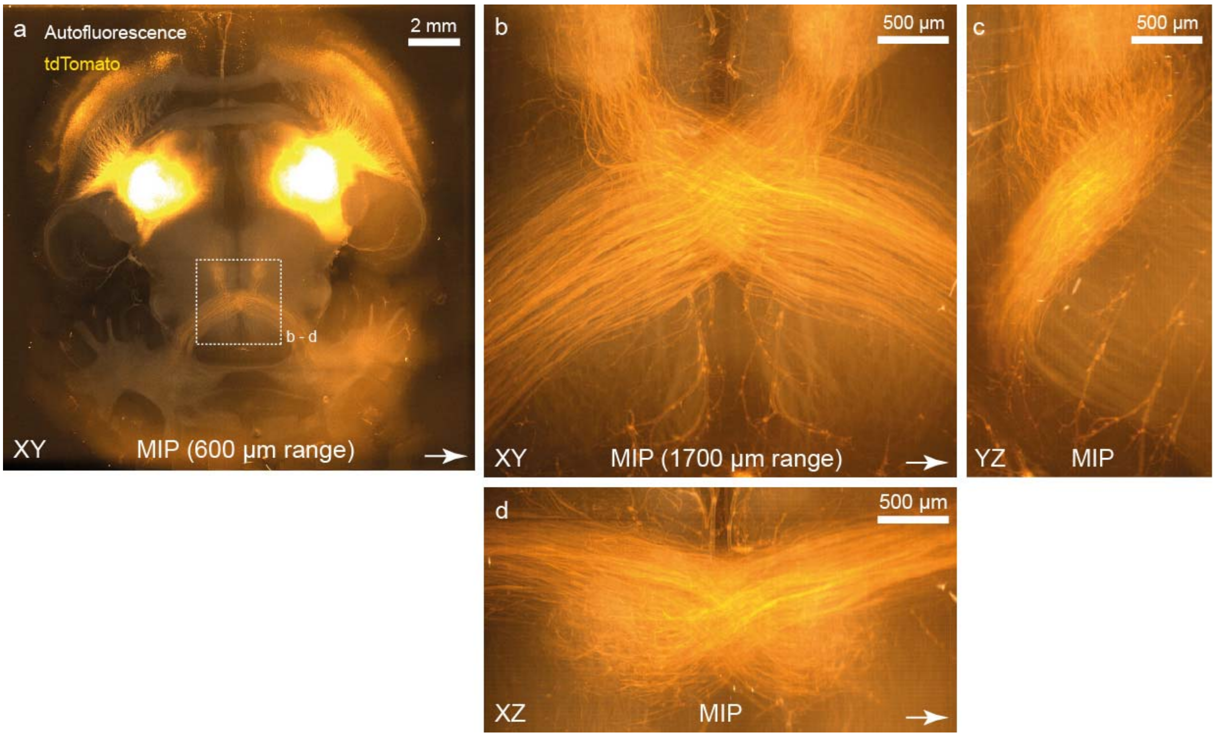

**Supplementary Figure 23: Imaging long-range projections in a CUBIC-X-cleared mouse brain.** a) Low-resolution overview image (Zoom 0.63x) of a cleared mouse brain (using CUBIC-X2) expressing tdTomato in a subset of layer 6 (L6) neurons (Ntsr1-Cre;CAG-tdTomato) b) Maximum intensity projection (MIP) over a 1700-µm z-range of the midline-crossing axons in the sample (location indicated in a). c & d: YZ and XZ projections of the same volume. Due to low signal levels, the images are very noisy and stripe artifacts from single camera pixels are visible. The arrow in the lower right corner indicates the direction of the light-sheet illumination.

### The mesoSPIM is compatible with BABB-cleared samples

The combination of benzyl alcohol and benzyl benzoate is among the oldest clearing techniques and was introduced in the early 20$^{th}$ century by Spalteholz[8,68]. It forms the ancestor of a wide variety of clearing techniques based on dehydration, delipidation and index matching using organic solvents – including 3DISCO[6,7,69], iDISCO[11], iDISCO+[70,71], uDISCO[72], vDISCO[56] and PEGASOS[73]. Compared to CLARITY and CUBIC, the refractive index of the employed organic solvents is usually >1.5, for example $n_D$ = 1.56 for BABB. As an example for imaging results achievable with the mesoSPIM, we stained a 7-day old chicken embryo (Hamburger Hamilton stage HH31) for neurofilament with secondary antibodies conjugated to Cy3 and mounted the sample in a 10×10×45 mm$^3$ glass cuvette (Portmann Instruments) filled with BABB. The imaging cuvette was then immersed in a 40×40×45 mm$^3$ immersion cuvette filled with BABB. With 561-nm excitation, the developing nervous system could be visualized in its entirety (**Supplementary Figure 24**). Using 405-nm excitation and a quadruple-band emission filter (QuadLine Rejectionband ZET405/488/561/640, AHF), it is possible to visualize the anatomy of the embryo using autofluorescence (**Supplementary Figure 25**). As the mesoSPIM contains a rotation stage, it is possible to perform multi-view acquisitions by specifying rotation increments in the Acquisition Manager Window of the mesoSPIM-control software. Especially for sample features that are highly absorbing and therefore cast shadows—such as the melanin-rich eyes—multidirectional acquisitions can allow "filling in" of missing information. In addition, multidirectional datasets can be fused to achieve more isotropic resolution, a technique common in developmental light-sheet microscopy[74] that has recently been applied to cleared samples[75]. As the sample was mounted in a quadratic cuvette, we recorded overview stacks at 90° intervals to avoid imaging through the corners of the cuvette (**Supplementary Figure 25**). If the sample is mounted in a clamping holder, more rotation angles (views) can be collected. By zooming in 4x and tiling the embryo in the 180° orientation, fine neurites in the developing nervous system can be discerned across the whole embryo (**Fig. 1f**; **Supplementary Figure 24**). For this, a 5x7 mosaic of stacks at zoom 4× was taken using 2-µm z-step size and fused into a single 670 GB dataset using Bigstitcher[76]. Single long-range axons can be discerned both in the developing brain (**Supplementary Figure 24b**) and the trunk (**Supplementary Figure 24c-e**). Reslicing the datasets of a single substack in the XZ direction reveals that fine processes can also be discerned in the axial direction (**Supplementary Figure 24d-e**), highlighting that the mesoSPIM allows near-isotropic imaging quality in such a sample (see also **Supplementary Video 9**).

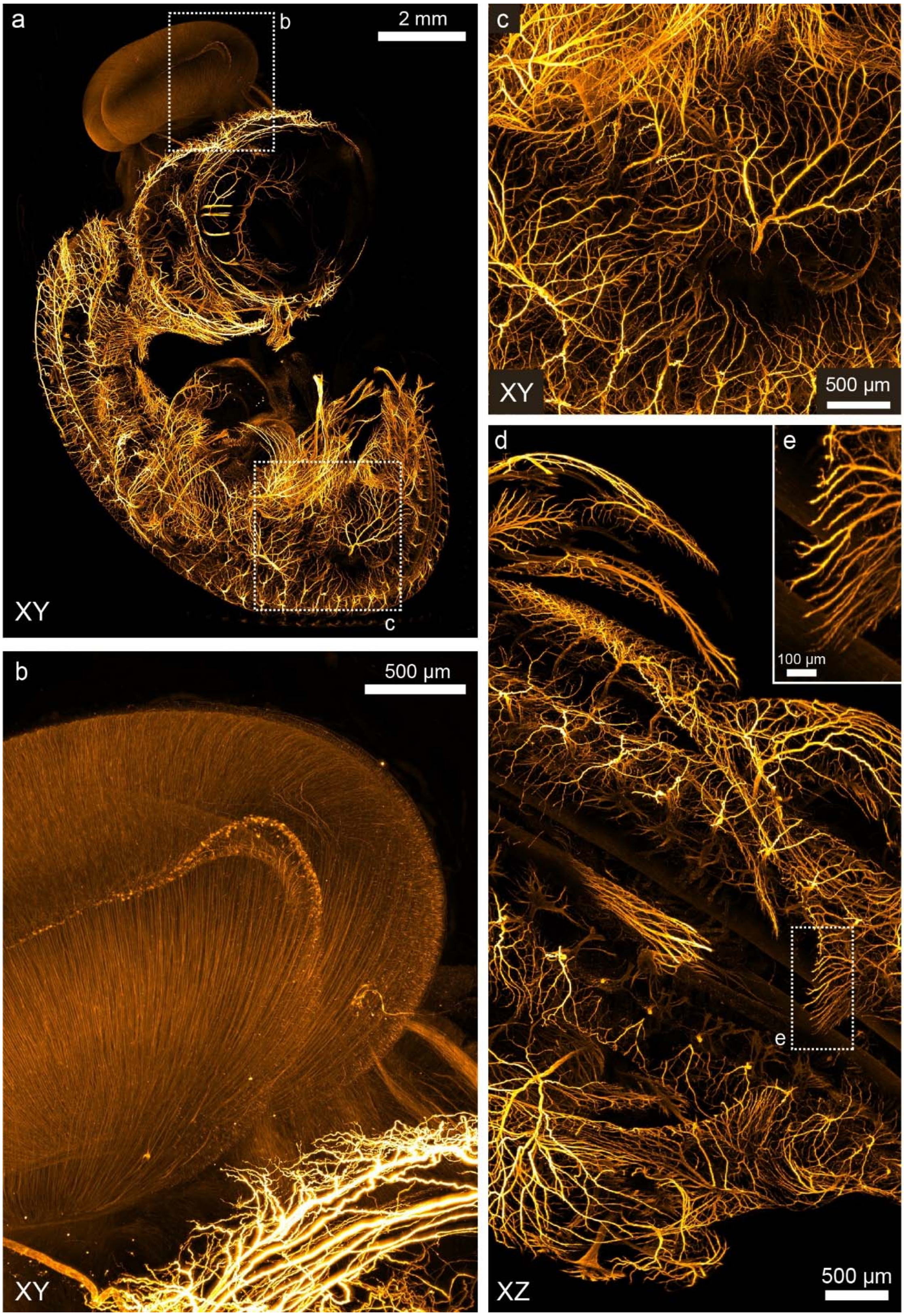

**Supplementary Figure 24: Multi-scale imaging in a BABB-cleared and neurofilament-stained 7-day old chicken embryo.** a) Overview image of the whole sample (Maximum intensity projection) taken at zoom 0.8x (Same sample as in Figure 1). b) & c) Higher magnification images (MIPs) taken at 4x. d) & e) Reslice (XZ view, MIP) of the substack shown c).

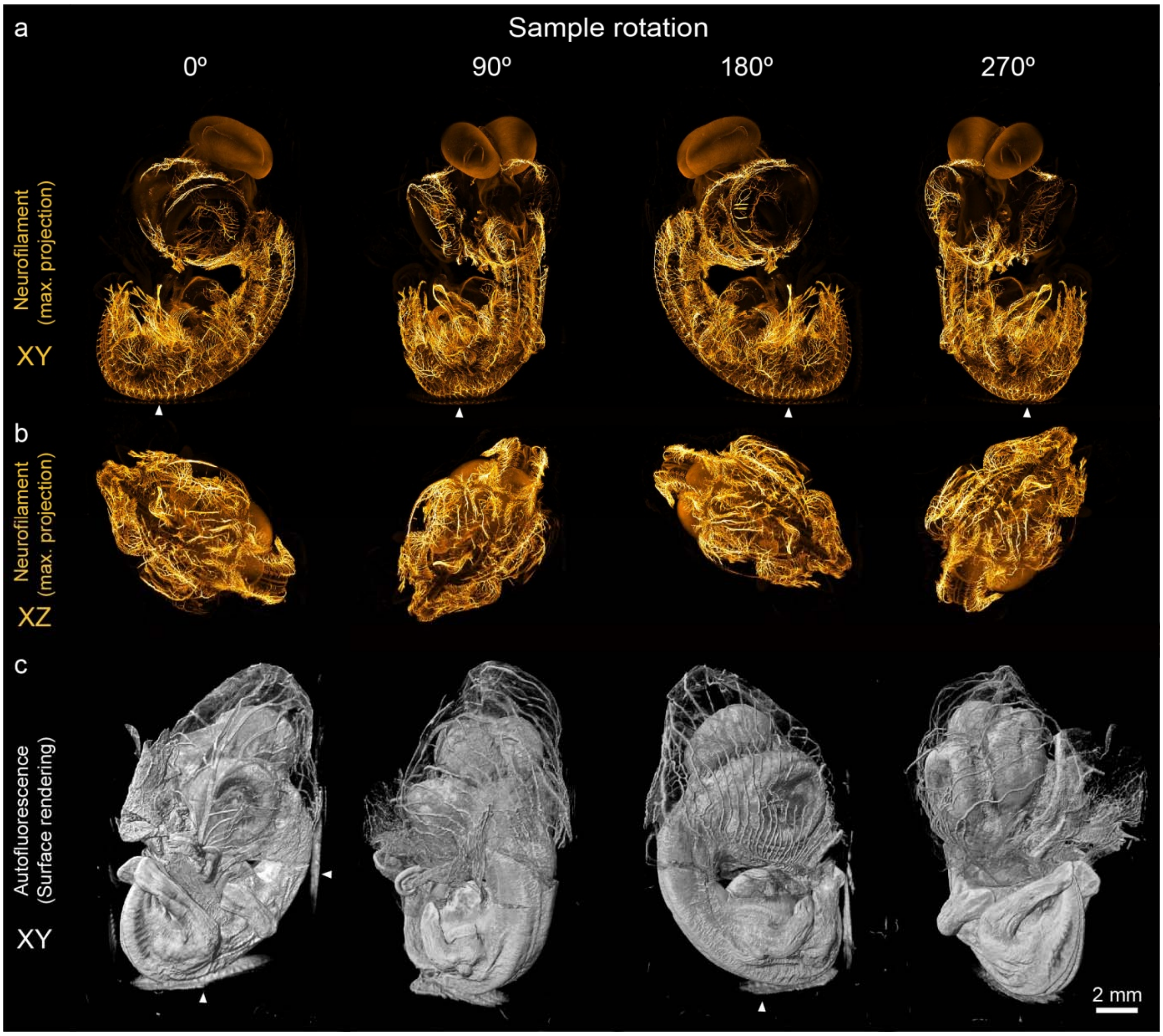

**Supplementary Figure 25: Multi-view imaging in a BABB-cleared and neurofilament-stained 7-day old chicken embryo.** a) MIPs of z-stacks (0.8x magnification) taken at different rotation angles of the sample (Cy3), excited at 561 nm. b) MIPs of the XZ (resliced) view of the datasets in a). c) Surface rendering of the autofluorescence signal excited at 405 nm. Due to the index mismatch between BABB and the quartz material of the cuvettes, reflections off the cuvette walls can occur (indicated by arrows).

As a further example for the imaging quality achievable with a mesoSPIM in a BABB-cleared sample, we cleared a white-eyed *Drosophila melanogaster* according to the protocol by Dodt et al.[9]. We then placed the sample in a 10×10×45 $mm^3$ glass imaging cuvette (Portmann Instruments) filled with BABB and immersed this cuvette in a 40x40x45 $mm^3$ immersion cuvette filled with BABB as well. Using the 1x objective, we scanned the head of the fly using 488-nm excitation (**Supplementary Figure 26Error! Reference source not found.** and **Supplementary Video 10**). In a surface rendering of the resulting dataset (**Supplementary Figure 26a**), a wide range of external anatomical features can be discerned, ranging from single ommatidia in the compound eyes to antennae and sensiliae around the head. Both in the XY and the XZ plane, a wide variety of internal anatomical features can be visualized as well – ranging from the internal structure of the mouthparts to subdivisions of the brain such as the layers of the optic lobe (**Supplementary Figure 26 b,c**). Despite its comparatively small size, it is thus possible to visualize details of the anatomy of *D. melanogaster* with a mesoSPIM using a 1x objective.

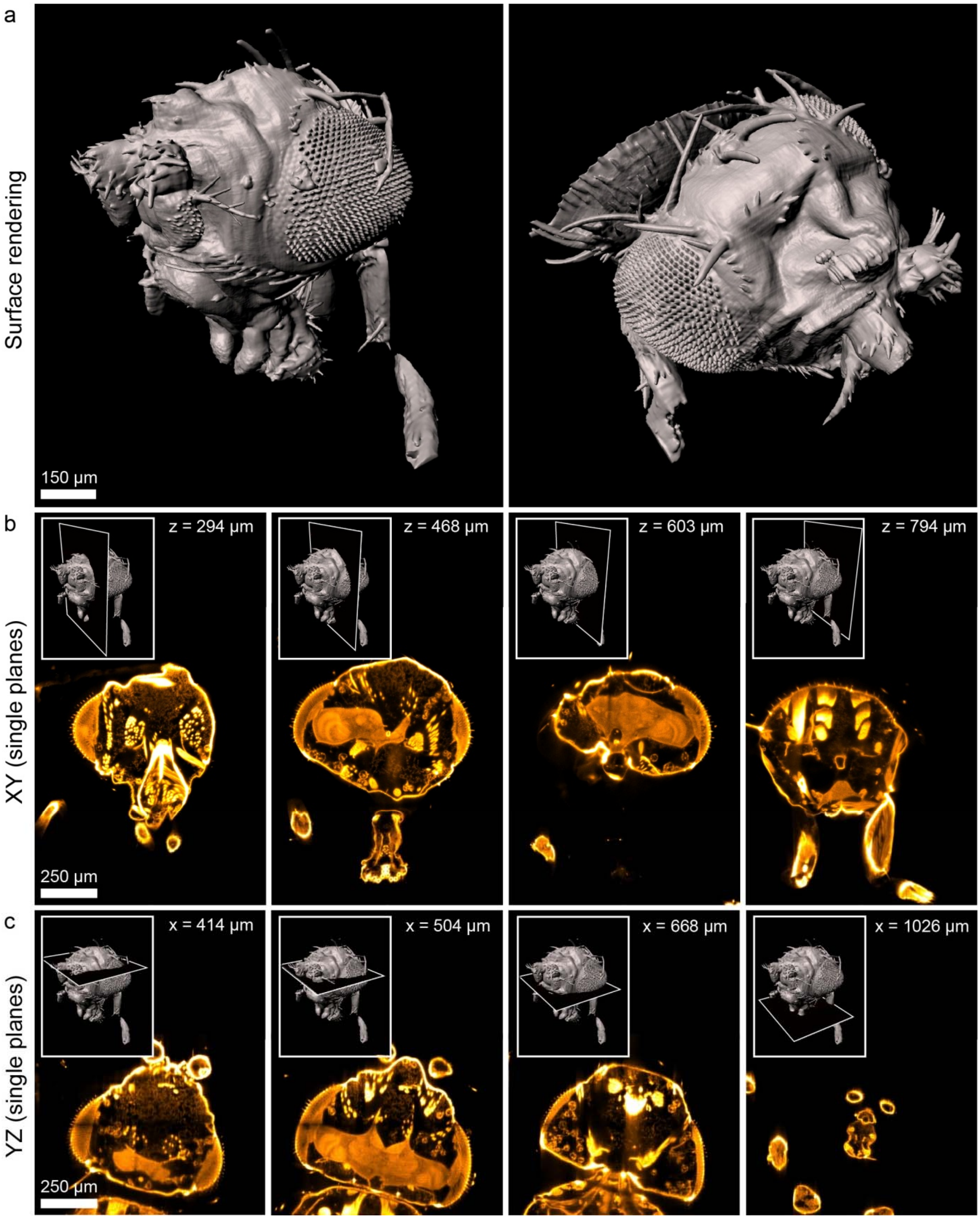

**Supplementary Figure 26: Anatomical imaging in a BABB-cleared *Drosophila melanogaster*.** A white-eyed fly was dehydrated and cleared with the BABB protocol. a) Surface renderings from a dataset acquired at 6.3× zoom using the 1× objective (1-µm pixel size). Cuticle and tissue autofluorescence was excited at 488 nm. b) Single XY planes at different axial locations (indicated by the insets). c) Single planes from the resliced view (YZ) view of the same dataset. A wide variety of fine anatomical features can be distinguished.

### The mesoSPIM is compatible with iDISCO-cleared samples

In recent years, the combination of 3DISCO clearing[6,7,69] and immunolabeling led to the iDISCO technique[11], which allows immunostaining in whole mouse brains for a wide range of antibodies. iDISCO samples are commonly imaged in dibenzylether (DBE), a high-index medium (nD = 1.562), which can dissolve plastics. As DISCO-samples tend to be small and hard, we clamp such samples in a 3D-printed custom sample holder made from polyamide (nylon) with nylon screws. In our hands, nylon is stable even during prolonged immersion in DBE. To demonstrate the compatibility of the mesoSPIM with iDISCO samples, we performed whole-brain immunostaining against immunoglobulin G (IgG) in a non-perfused mouse brain (**Supplementary Figure 27a-b**). Without perfusion, the vasculature retains IgG in the serum which can be stained using secondary antibodies. Even in low-magnification overviews taken at zoom 1.25×, details of the vasculature can be discerned in the XY and XZ views (**Supplementary Figure 27c-e**). We noted that given the high transparency of the tissue and excellent labeling quality, imaging with a single light-sheet was sufficient as there was no intensity gradient visible across the sample. In addition, optimizing the tunable lens parameters was straightforward as the length of the blood vessel sections visible in every XY plane is a good indicator for the effective thickness of the light-sheet. As the staining is done using passive diffusion of the antibody, there is a labeling gradient from outside to inside, but even in the central parts of the sample, the labeling quality is sufficient to discern small capillaries in higher resolution datasets taken at 4× magnification (3.29 mm FOV; **Supplementary Figure 28**, **Supplementary Video 11**).

To demonstrate that the mesoSPIM is capable both of imaging rat brain tissue and doing multi-color acquisitions in iDISCO samples, we performed a nuclear stain on a wildtype rat using To-Pro (**Supplementary Figure 29**).

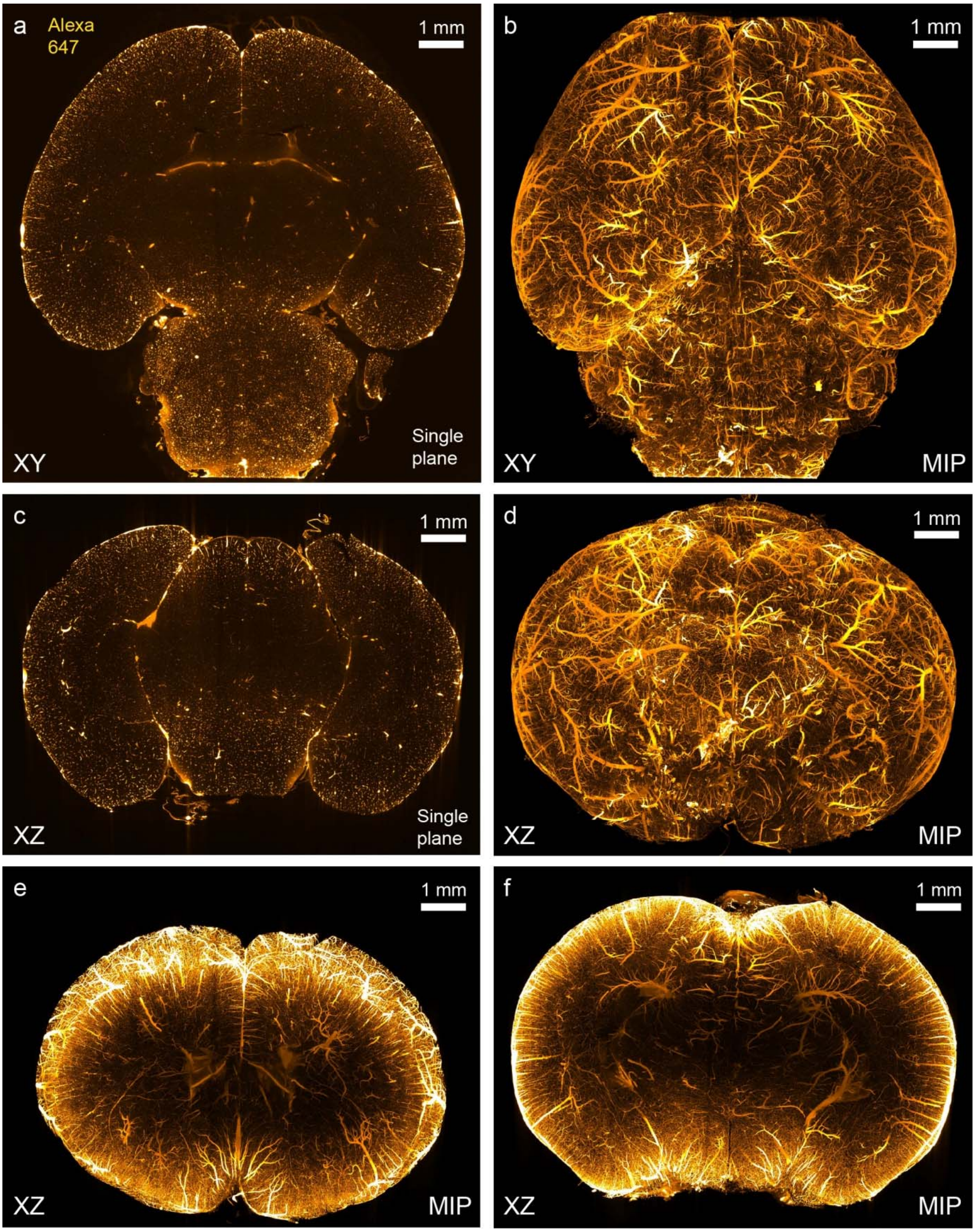

**Supplementary Figure 27: The mesoSPIM is compatible with iDISCO-cleared samples.** A whole mouse brain was stained for immunoglobulin G (IgG) conjugated to Alexa 647 antibodies and cleared using the iDISCO protocol. a) Single plane taken from a low-resolution overview stack (Zoom 1.25×). b) Maximum intensity projection (MIP) covering the dorsal aspect of the brain. Large vessels on the surface are prominent. c) Single resliced plane (XZ view) from the dataset. d) MIP of the frontal aspect of the sample. e) and f) MIP (500-µm range) in the coronal plane. As antibody penetration is limited by diffusion, an inside-out intensity gradient is visible.

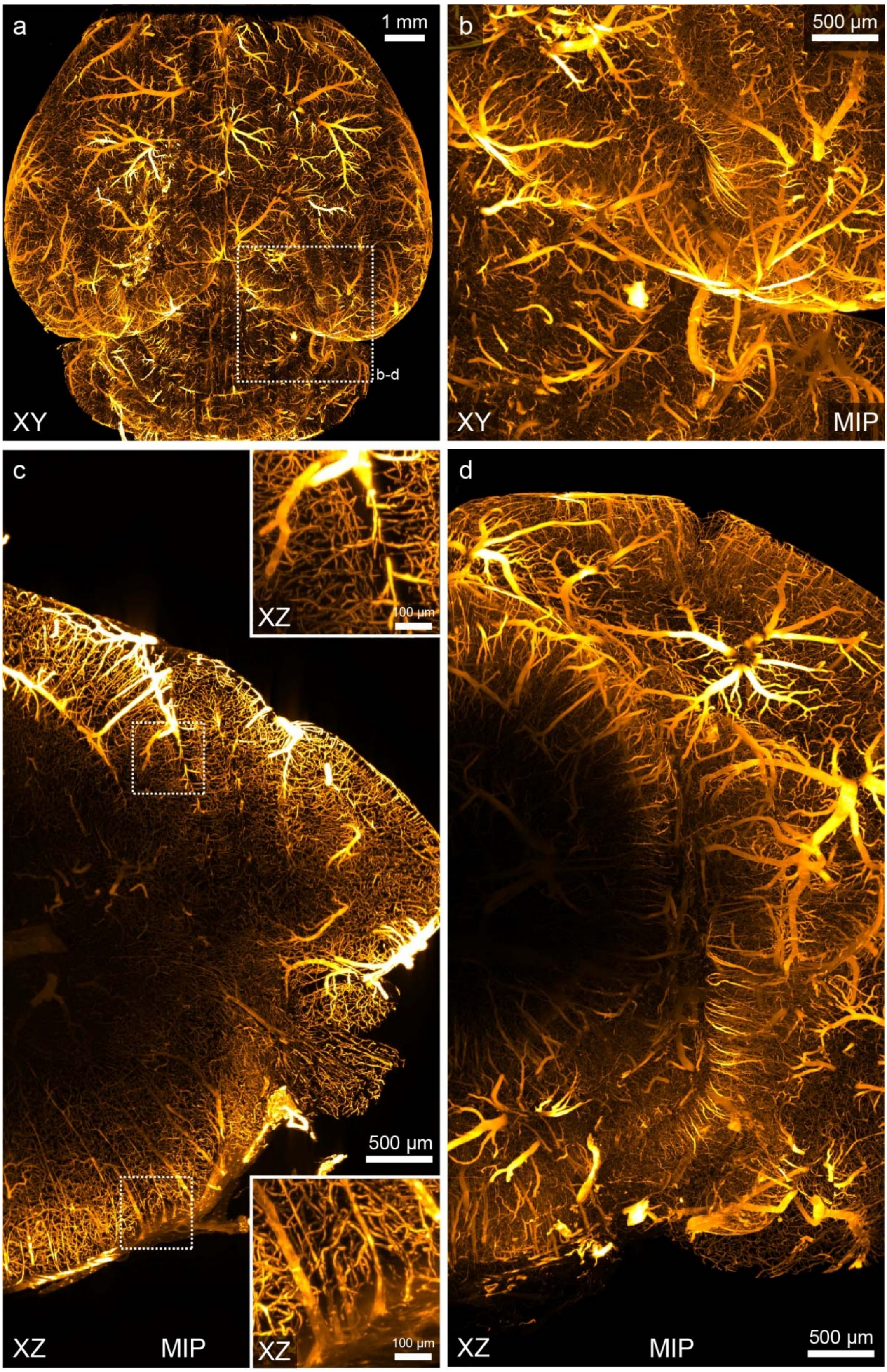

**Supplementary Figure 28: Near-isotropic imaging in an iDISCO-cleared mouse brain stained for vasculature.** a) The same sample as in Supplementary Figure 27 was scanned at zoom 4× (3.29-mm FOV) at the indicated location. b) Maximum projection (MIP) of the whole stack. c) and d) MIPs (250-µm range) of the acquired dataset. The insets in c) demonstrate that even in the XZ view, fine capillaries are well separated from each other.

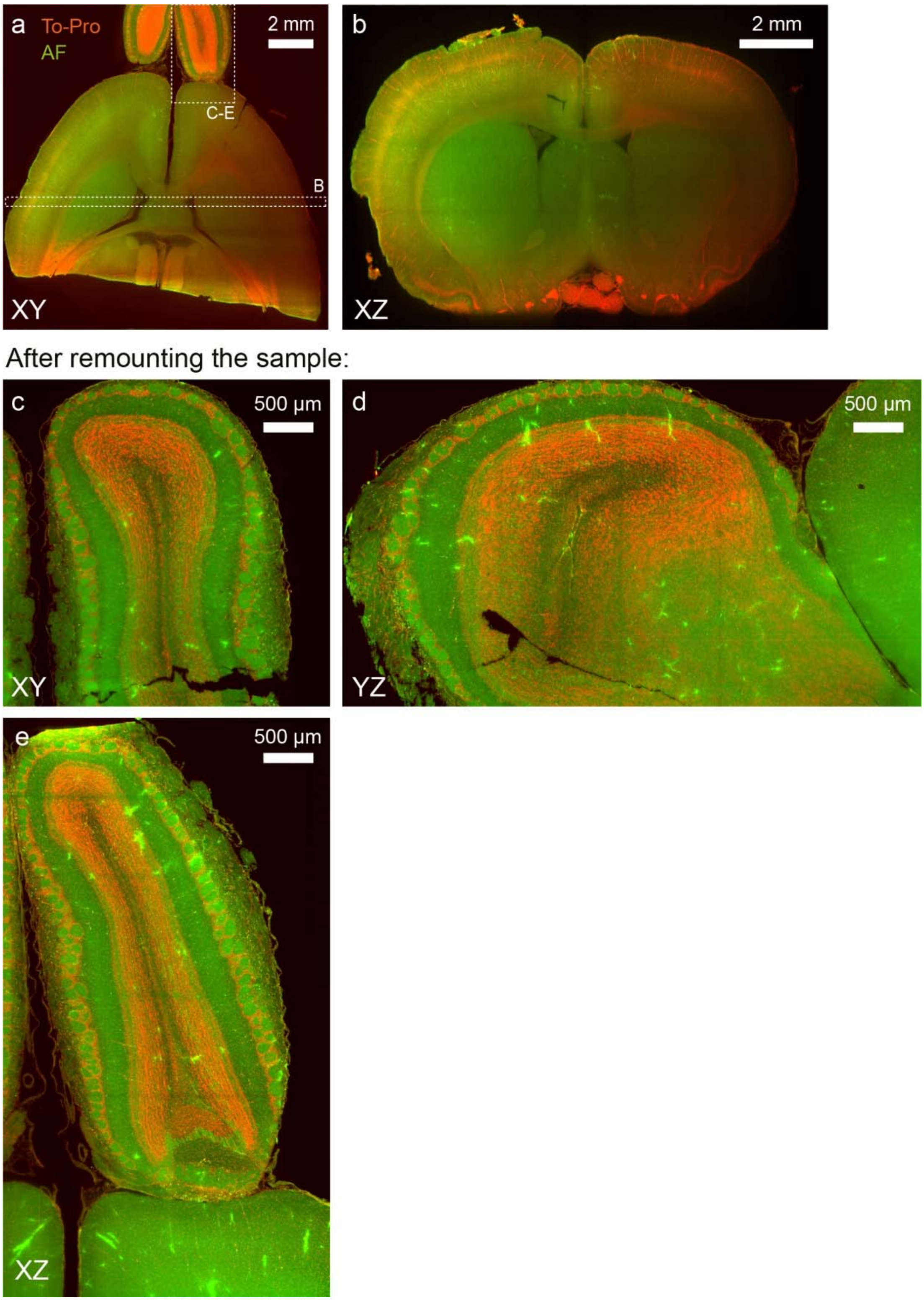

**Supplementary Figure 29: Multicolor imaging in an iDISCO-cleared and To-Pro-stained rat brain.** a) Overview image (maximum projection, 0.5 mm range) of a rat forebrain**.** The nuclear stain (To-Pro) is shown in red and the autofluorescence (AF) in green. b) Reslice (XZ view) of a subvolume of the sample in a). After remounting the sample (coronal planes parallel to the light-sheet), the olfactory bulb was imaged again (c-e). In all sectioning planes, single glomeruli can be seen.

**The mesoSPIM is compatible with MASH (iDISCO/ECi)-processed samples**

Given the size and complexity of the human brain, clearing techniques capable of rendering human samples sufficiently transparent for large-scale studies are of considerable interest for neuroscience and pathology. However, owing to its myelination and sheer size, clearing human brain tissue is challenging. In addition, samples are usually much older than the mouse brains used in the demonstration of clearing techniques, which leads to increased levels of autofluorescence (for example due to an increase in lipofuscin). However, a variety of clearing methods was shown to be compatible with human tissue, for example OPTIClear[77]. Recently it was shown that 'multiscale architectonic staining of human cortex' (MASH)[78] is possible by a modified iDISCO+ clearing protocol and with the possibility of using ethyl cinnamate (ECi) for index matching. Compared to other index matching solutions, the non-toxicity of ECi, which was established in 2017[79], makes it very user-friendly. Using this technique in combination with cell nucleus label MASH-MG (based on known protocols with Methyl Green) and cell body label MASH-NR (based on standard protocols for Nissl staining with Neutral Red) staining, we were able visualize the structure of the human neocortex (see **Supplementary Figure 30**). The 10×20×5 $mm^3$ slab of MASH-processed tissue was placed inside a 10×20×45 $mm^3$ glass cuvette filled with ECi and submerged in an immersion cuvette filled with ECi. Imaging at low zoom allowed localizing cortical folds (**Supplementary Video 12**) while datasets taken at 4× zoom reveal the distribution of) neurons across the layers of the neocortex (**Supplementary Video 13**).

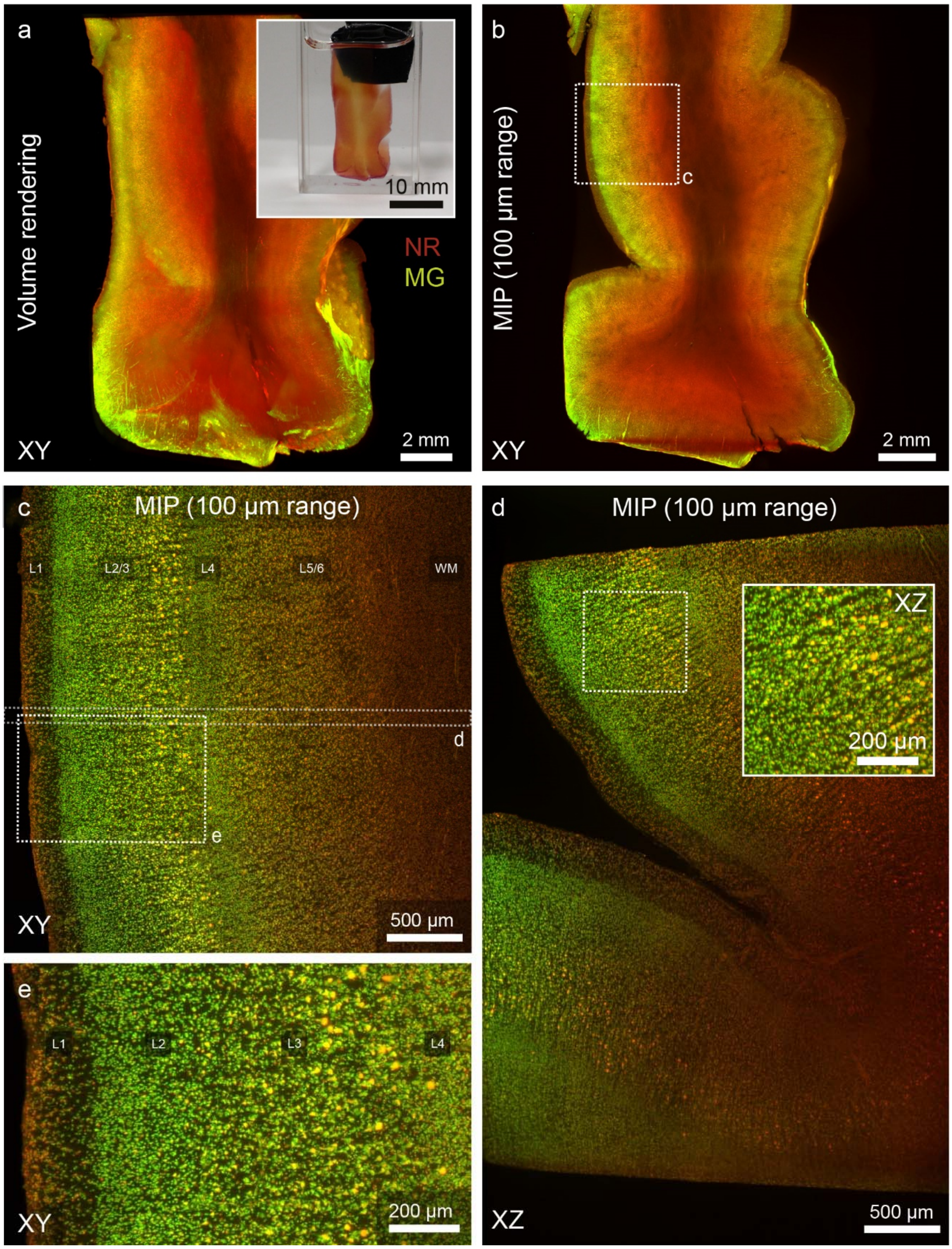

**Supplementary Figure 30: The mesoSPIM is compatible with MASH (iDISCO/ECi)**-processed **human neocortex samples.** a) Volume rendering of data acquired from a slab of human cortex cleared using ethyl cinnamate (ECi) and stained with Neutral Red (NR) and Methyl Green (MG) using the multiscale architectonic staining of tissue protocol (MASH)[78]. The inset shows the sample immersed in ECi in the imaging cuvette. b) Maximum intensity projection (MIP) of a part of the dataset in a) c) MIP of a dataset acquired at 4x magnification (location shown in b). d) Reslice of a part of the dataset (location) indicated in c). Even in the axial direction, cells can be resolved easily. In the XZ view, the better penetration of the red wavelengths is visible. e) Detail of c). Cortical layers 1 to 6 (L1 to L6) and white matter (WM) are indicated.

| Figure | Panel | Obj. | Zoom | Sample | Clearing method | Ill. Dir. | Excitation | Filter | $t_{sweep}$ | $t_{exp}$ | $f_{galvo}$ | Pixel size (X×Y×Z) | Tiles/Conditions × X × Y × Z | $t_s$ [min] | $t_{tot}$ [min] | File size | File size (total) | Comment |
|---|---|---|---|---|---|---|---|---|---|---|---|---|---|---|---|---|---|---|
| 1 | b | 1× | 1× | VIP-tdTomato mouse brain | Passive CLARITY | Right | 561 nm | 561LP | 200 ms | 20 ms | 99.9 Hz | 6.55 × 6.55 × 5 µm$^3$ | 2 × 2048 × 2048 × 1937 | 7.37 | 14.85 | 11.9 GB | 23.8 GB | |
| | c | 1× | 0.63× | TPH2::Cre;Rosa26$^{tdTomato}$ mouse brain | Passive CLARITY | Right | 561 nm | 561LP | 200 ms | 20 ms | 99.9 Hz | 10.5 × 10.5 × 5 µm$^3$ | 1 × 2048 × 2048 × 2107 | 8.03 | 8.03 | 15.58 GB | 15.58 GB | |
| | d/e | 1× | 4× | TPH2::Cre;Rosa26$^{tdTomato}$ mouse brain | Passive CLARITY | Right | 561 nm | 561LP | 200 ms | 20 ms | 199 Hz | 1.6 × 1.6 × 2 µm$^3$ | 5 × 2048 × 2048 × 3100 | 12.54 | 62.7 | 17.62 GB | 88.1 GB | |
| | f | 1× | 4× | Chick embryo (7d) | BABB | Right | 561 nm | 561LP | 200 ms | 5 ms | 199 Hz | 1.5 × 1.5 × 2 µm$^3$ | 24 × 2048 × 2048 × 4700 | 17.82 | 429 | 38.5 GB | 880 GB | |
| **Supplementary Figures** | | | | | | | | | | | | | | | | | | |
| 2 | a-f | 1× | 1× | VIP-tdTomato mouse brain | Passive CLARITY | Right | 561 nm | 561LP | 200 ms | 20 ms | 99.9 Hz | 6.55 × 6.55 × 5 µm$^3$ | 2 × 2048 × 2048 × 1937 | 7.37 | 14.85 | 11.9 GB | 23.8 GB | Same dataset as in 1b |
| 3 | a-h | 1× | 1× | Thy1-YFP mouse brain | Active CLARTIY | Right | 488 nm | 542/27 | 200 ms | 20 ms | 99.9 Hz | 6.55 × 6.55 × 2 µm$^3$ | 2 × 2048 × 2048 × 4200 | 15.95 | 31.9 | 34.4 GB | 68.8 GB | |
| 4 | a-d | 1× | 4× | Thy1-YFP mouse brain | Active CLARTIY | Right | 488 nm | 542/27 | 200 ms | 10 ms | 199 Hz | 1.6 × 1.6 × 2 µm$^3$ | 2 × 2048 × 2048 × 4200 | 15.95 | 31.9 | 34.4 GB | 68.8 GB | |
| 5 | a | 1× | 4× | Thy1-YFP mouse brain | Active CLARTIY | Right | 488 nm | 542/27 | 200 ms | 10 ms | 199 Hz | 1.6 × 1.6 × 2 µm$^3$ | 3 × 2048 × 2048 × 4200 | 15.95 | 47.85 | 34.4 GB | 103.2 GB | |
| 6 | a-c | 1× | 1× | Thy1-YFP mouse brain | Active CLARTIY | Left/Right/Both | 488 nm | 542/27 | 200 ms | 20 ms | 99.9 Hz | 6.55 × 6.55 × 3 µm$^3$ | 3 × 2048 × 2048 × 2800 | 13.42 | 40.26 | 22.9 GB | 68.7 GB | |
| 14 | | 1× | 1× | 1 µm bead test sample at nD=1.45 | n/a | Right | 488 nm | 520/35 | 200 ms | 5 & 10 ms | 99 & 199 Hz | 6.55 × 6.55 × 1 µm$^3$ | 3 × 2048 × 2048 × 200 | 0.77 | 2.75 | 1.6 GB | 4.8 GB | |
| 15 | | 1× | 4× | 1 µm bead test sample at nD=1.45 | n/a | Right | 488 nm | 520/35 | 200 ms | 5 & 10 ms | 99 & 199 Hz | 1.6 × 1.6 × 1 µm$^3$ | 3 × 2048 × 2048 × 200 | 0.77 | 2.75 | 1.6 GB | 4.8 GB | |
| 16 | | 1× | 1× | Thy1-YFP mouse brain | Active CLARTIY | Right | 488 nm | 520/35 | 200 ms | 5-40 ms | 199 Hz | 6.55 × 6.55 × 2 µm$^3$ | 4 × 2048 × 2048 × 4200 | 15.95 | 64.9 | 34.4 GB | 137.6 GB | |
| 17 | | 1× | 4× | Thy1-YFP mouse brain | Active CLARTIY | Right | 488 nm | 520/35 | 200 ms | 5-40 ms | 199 Hz | 1.6 × 1.6 × 2 µm$^3$ | 4 × 2048 × 2048 × 4200 | 15.95 | 64.9 | 34.4 GB | 137.6 GB | |
| 18 | a-d | 1× | 1× | YCX 2.60 mouse brain | Passive CLARITY | Right | 515 nm & 647 nm | 515LP & QB | 200 ms | 20 ms | 99.9 Hz | 6.55 × 6.55 × 5 µm$^3$ | 2 × 2048 × 2048 × 1674 | 6.38 | 12.65 | 13 Gb | 26 GB | |
| 19 | a | 1× | 1× | YCX 2.60 mouse brain | Passive CLARITY | Right | 515 nm & 647 nm | 515LP & QB | 200 ms | 20 ms | 99.9 Hz | 6.55 × 6.55 × 5 µm$^3$ | 2 × 2048 × 2048 × 1674 | 6.38 | 12.65 | 13 Gb | 26 GB | |
| | b-d | 1× | 4× | YCX 2.60 mouse brain | Passive CLARITY | Right | 515 nm | 515LP | 200 ms | 20 ms | 199 Hz | 1.6 × 1.6 × 5 µm$^3$ | 1 × 2048 × 2048 × 1310 | 5.06 | 5.06 | 10.2 GB | 10.2 GB | |
| 20 | b-c | 1× | 4× | YCX 2.60 mouse brain | Passive CLARITY | Right | 515 nm | 515LP | 200 ms | 20 ms | 199 Hz | 1.6 × 1.6 × 3 µm$^3$ | 1 × 2048 × 2048 × 766 | 2.97 | 2.97 | 6 GB | 6 GB | |
| 21 | a-b | 1x | 1x | GlyT2-EGFP mouse CNS | Active CLARTIY | Left/Right | 488 nm &561 nm | 520/35 @ 561LP | 200 ms | 20 ms | 199 Hz | 6.55 × 6.55 × 5 µm$^3$ | 16 × 2048 × 2048 × 1679 | 7.5 | 165 | 13.771 GB | 220 GB | |
| 21 | c | 1x | 4x | GlyT2-EGFP mouse CNS | Active CLARTIY | Right | 488 nm | 520/35 | 200 ms | 20 ms | 199 Hz | 1.6 × 1.6 × 2 µm$^3$ | 1 × 2048 × 2048 × 2469 | 10.24 | 10.24 | 19.2 GB | 19.2 GB | |
| 21 | d | 1x | 4x | GlyT2-EGFP mouse CNS | Active CLARTIY | Left/Right | 488 nm | 520/35 | 200 ms | 20 ms | 199 Hz | 1.6 × 1.6 × 2 µm$^3$ | 2 × 2048 × 2048 × 994 | 4.1 | 8.2 | 8.14 GB | 15.5 GB | |
| 22 | a-d | 1× | 0.63× | Ntsr1-Cre;CAG-tdTomato) mouse brain | CUBIC-X | Left | 561 nm & 488 nm | 561LP | 200 ms | 20 ms | 99.9 Hz | 10.5 × 10.5 × 3 µm$^3$ | 2 × 2048 × 2048 × 3566 | 13.53 | 29.48 | 29.2 GB | 58.4 GB | |
| 23 | a-d | 1× | 4× | Ntsr1-Cre;CAG-tdTomato) mouse brain | CUBIC-X | Left | 561 nm & 488 nm | 561LP | 200 ms | 20 ms | 199 Hz | 1.6 × 1.6 × 2 µm$^3$ | 2 × 2048 × 2048 × 4700 | 17.93 | 36.3 | 38.5 GB | 77 GB | |
| 24 | a-d | 1× | 4× | Chick embryo (7d) | BABB | Right | 561 nm | 561LP | 200 ms | 5 ms | 199 Hz | 1.5 × 1.5 × 2 µm$^3$ | 24 × 2048 × 2048 × 4700 | 17.82 | 429 | 38.5 GB | 880 GB | Same dataset as in 1f |
| 25 | | 1× | 0.8× | Chick embryo (7d) | BABB | Right & Both | 405 nm & 561 nm | QB & 561LP | 200 ms | 20 ms | 99.9 Hz | 7.8 × 7.8 × 2 µm$^3$ | 8 × 2048 × 2048 × 5200 | 19.8 | 2.64 | 42 GB | 319 GB | 4 rotation angles |
| 26 | | 1× | 4× | D. melanogaster | BABB | Right | 488 nm | QB | 200 ms | 10 ms | 99.7 Hz | 1 × 1 × 1 µm$^3$ | 1 × 2048 × 2048 × 1700 | 6.49 | 6.49 | 13.4 GB | 13.4 GB | |
| 27 | | 1× | 1.25 × | Mouse brain (vasculature) | iDISCO | Right | 647 nm | QB | 200 ms | 10 ms | 199 Hz | 5 × 5 × 2 µm$^3$ | 1 × 2048 × 2048 × 4072 | 15.4 | 15.4 | 31.8 GB | 31.8 GB | |
| 28 | | 1× | 4× | Mouse brain (vasculature) | iDISCO | Right | 647 nm | QB | 200 ms | 10 ms | 199 Hz | 1.55 × 1.55 × 2 µm$^3$ | 16 × 2048 × 2048 × 3186 | 12.1 | 191.4 | 24.8 GB | 398 GB | |
| 29 | a-b | 1× | 1× | Rat brain (TO-PRO) | iDISCO | Right | 488 nm & 647 nm | QB & QB | 200 ms | 20 ms | 99.9 Hz | 6 × 6 × 3 µm$^3$ | 2 × 2048 × 2048 × 2966 | 14.19 | 28.38 | 24.3 GB | 48.6 GB | |
| | c-d | 1× | 4× | Rat brain (TO-PRO) | iDISCO | Right | 488 nm & 647 nm | QB & QB | 200 ms | 10 ms | 199 Hz | 1.55 × 1.55 × 1 µm$^3$ | 2 × 2048 × 2048 × 5800 | 22 | 44 | 47.5 GB | 95 GB | |
| 30 | a-b | 1× | 0.8× | Human cortex | MASH (iDISCO/ECi) | Both | 561 nm & 647 nm | 561LP & QB | 200 ms | 20 ms | 99.9 Hz | 7.8 × 7.8 × 2 µm$^3$ | 2 × 2048 × 2048 × 2500 | 9.46 | 19.14 | 20.5 GB | 41 GB | |
| | c-e | 1× | 4× | Human cortex | MASH (iDISCO/ECi) | Left | 561 nm & 647 nm | 561LP & QB | 200 ms | 10 ms | 199 Hz | 1.6 × 1.6 × 1 µm$^3$ | 2 × 2048 × 2048 × 5000 | 18.92 | 37.95 | 41 GB | 82 GB | |

**Supplementary Table 4: Overview of all imaging parameters.** All datasets were taken with the MVPLAPO1x objective (Obj.). In addition, illumination direction (Ill.dir.), sweeptime (total waveform generation time, $t_{sweep}$), exposure time ($t_{exp}$), galvo frequency ($f_{galvo}$), and the acquisition time for a single stack in a multi-color or tiled acquisition ($t_s$) and for the entire acquisition ($t_{tot}$) are indicated. In addition, the size of the dataset for a single stack and for the entire acquired dataset are given. QB refers to a quadrupleband blocking filter (QuadLine Rejectionband ZET405/488/561/640, AHF). 561LP refers to a 561 nm longpass.

**Supplementary Note: Sample preparation**

**Overview of mouse lines**

An overview of all mouse lines used for this study is given in Supplementary Table 2. Crosses were performed as indicated in the text.

| Short name | Official strain name | Repository stock number |
|---|---|---|
| VIPCre | Viptm1(cre)Zjh/J | JAX 010908 |
| Ai14 (tdTomato) | B6.Cg-Gt(ROSA)26Sortm14(CAG-tdTomato)Hze/J | JAX 007914 |
| TPH2Cre | Tg(Tph2-cre)RH35Gsat/Mmucd | MMRRC 036634-UCD |
| Camk2a-tTA | B6.Cg-Tg(Camk2a-tTA)1Mmay/DboJ | JAX 007004 |
| Ai92(YCX2.60) | B6.Cg-Igs7tm92.1(tetO-ECFP*/Venus*)Hze/J | JAX 026262 |
| Rbp4Cre | Tg(Rbp4-cre)KL100Gsat/Mmucd | MMRRC 031125-UCD |
| Thy1-YFP | B6.Cg-Tg(Thy1-YFP)HJrs/J | JAX 003782 |
| GlyT2-EGFP | Tg(Slc6a5-EGFP)1Uze | MGI 3835459 |
| Ntsr1Cre | B6.FVB(Cg)-Tg(Ntsr1-cre)GN220Gsat/Mmucd | MMRRC 030648-UCD |
| Ai9 (tdTomato) | B6;129S6-Gt(ROSA)26Sortm9(CAG-tdTomato)Hze/J | JAX 007905 |

**Supplementary Table 3: Overview of mouse lines used in this study.**

**Passive CLARITY clearing of mouse brains**

To demonstrate the compatibility of the mesoSPIM with passive CLARITY clearing, we cleared mouse brains from VIPCre-tdTomato ((Viptm1(cre)Zjh/J); B6.Cg-Gt(ROSA)26Sortm14(CAG-tdTomato)Hze/J)[80], TPH2Cre-tdTomato (Tg(Tph2-cre)RH35Gsat/Mmucd; B6.Cg-Gt(ROSA)26Sortm14(CAG-tdTomato)Hze/J), and Yellow-

Cameleon YCX2.60 (Camk2a-tTA;Rbp4-Cre;TITL-YCX2.60)[65]. The method used for hydrogel-based tissue clearing is described in detail elsewhere[10,60,61]. In short, the animals were transcardially perfused first with PBS followed by an ice-cold hydrogel solution (1% PFA, 4% acrylamide, 0.05% bis-acrylamide). The brains were dissected and post-fixed in the same hydrogel solution for 24h at 4°C. The samples were then degassed using a dessicator before hydrogel polymerization was induced at 37°C for 2-3hours. Following the polymerization, excess hydrogel was removed from the brains and they were immersed in 40mL of 8% SDS and kept shaking at room temperature until the tissue was cleared sufficiently (30+ days for an adult animal). Finally, after 2-4 washes in PBS, the brains were put into a self-made refractive index matching solution (RIMS)[60] for the last clearing step. They were left to euqilibrate in 5mL of RIMS for at least 4 days before being imaged. After clearing, brains were attached to a small weight and loaded into a 10×20×45 $mm^3$ quartz cuvette (UQ-205, Portmann Instruments), then submerged in RIMS and imaged using the mesoSPIM. The sample cuvette was immersed in a 40×40×40 $mm^3$ quartz cuvette (UQ-753, Portmann Instruments) filled with index-matching oil (19569, Code 50350, Cargille, nD=1.45), which allows sample XYZ movements and rotations without refocusing the detection path. This set of animal experiments and procedures were performed in accordance with standard ethical guidelines and were approved by the Cantonal Veterinary Office of the Canton of Zurich.

**Active CLARITY clearing of Thy1-YFP mouse brains**

Nine-weeks-old Thy1-YFP mice[63] were deeply anaesthetized with intraperitoneal injection of a mixture of 150 µl Ketamine (Ketalar, Bayer AG), 75 µl Xylazine (Rompun, Parke-Davis) and 75 µl sterile water. When mice seized to breath and no toe reflex was present, mice were transcardially perfused with 20 ml ice cold phosphate buffered saline (PBS) after which 20 ml ice cold hydrogel monomer-fixative (4% acrylamide, 1% paraformaldehyde, 0.05% bis-acrylamide, 1% VA-044 initiator in phosphate buffered saline)[61] was infused. Harvested brains were fixed in 20 ml ice-cold hydrogel monomer fixative overnight. Brains were degassed in a vacuum exicator for 20 minutes at ~-0.8 bar, followed by purging with nitrogen gas. The hydrogel monomer was polymerized at 37°C for 2.5 hours in tightly closed tubes. Samples were extracted from the hydrogel and transferred into clearing solution (8% sodium-dodecyl-sulphate, 200 mM boric acid, pH 8.5). Brains were optically cleared with clearing solution based on the CLARITY method[61] in a custom-built electrophoretic setup in 5 hours. Samples were washed in PBS three times and then transferred into RIMS., RIMS was replaced once to reach RI 1.46. Samples were stored light-protected at 4°C until imaged. After curing, the sample was immersed in a 10×20×45 $mm^3$ quartz cuvette for imaging. This set of animal experiments and procedures were performed in accordance with standard ethical guidelines and were approved by the Cantonal Veterinary Office of the Canton of Zurich.

**Whole-CNS imaging of X-CLARITY cleared samples**

Mice (GlyT2::eGFP (Tg(Scl6a5-EGFP)1Uze)[66], were perfused transcardially with 10 ml of artificial cerebrospinal fluid (ACSF: 125 mM NaCl, 2.5 mM KCl, 1.25 mM $NaH_2PO_4$, 25 mM $NaHCO_3$, 1 mM $MgCl_2$, 2 mM $CaCl_2$, 20 mM glucose equilibrated with 95% $O_2$, 5% $CO_2$) at room temperature (RT) followed by 20 ml of RT 4% paraformaldehyde (PFA, in 0.1 M sodium phosphate buffer, pH 7.4). The perfusion was performed using a gravity perfusion setup. Brain and spinal cord attached were dissected and put in 4% PFA overnight. They were then put in 4% acrylamide (161–0140; Bio-Rad) and 0.25% VA-044 (017–19362; Novachem) in PBS at 4°C. They were then incubated for 3 hours at 37°C for acrylamide polymerization, washed overnight at 37°C in clearing solution (200 mM SDS (L3371; Sigma-Aldrich) and 200 mM boric acid (L185094; Sigma-Aldrich), pH 8.5), and electrophoresed in clearing solution using an X-CLARITY Tissue Clearing System (Logos Biosystems) for 8 hours at 1.2 A constant current, temperature <37°C, and 100 rpm pump speed. The samples were incubated in approximately 88% Histodenz (D2158; Sigma-Aldrich) solution in PBS (refractive index adjusted to 1.457) overnight, and mounted for imaging in the same solution. To accommodate the whole CNS, the sample was placed in a custom quartz imaging cuvette (10×20×120 $mm^3$, Portmann Instruments AG) and then placed in a custom 40×40×120 $mm^3$ quartz immersion cuvette (Portmann Instruments AG). To ease mounting inside the sample cuvette, the sample was attached to a 1×13 cm strip of black aluminium foil (Thorlabs BKF12) using quick glue. This set of animal experiments and procedures were performed in accordance with standard ethical guidelines and were approved by the Cantonal Veterinary Office of the Canton of Zurich.

**CUBIC-X clearing and imaging**

The clearing process followed the protocol described in Murakami et al.[67] Adult animals (Ntsr1-cre, strain: B6.FVB(Cg)-Tg(Ntsr1-cre)GN220Gsat/Mmcd,; crossed with LSL-tdTomato, strain: B6;129S6-Gt(ROSA)26Sortm9(CAG-tdTomato)Hze/; aged P50) were perfused for five minutes with cold PBS, then for five minutes with cold 4% PFA. Brains were dissected after perfusion and incubated in 4% PFA overnight. Brains were delipidated for 14 days. After delipidation, brains were washed in PBS overnight then incubated in 30 mL of 20% imidazole (VWR, AAA10221-36) at 4°C for 2.5 days. An expanded brain was immersed in 40 ml of CUBIC-X2 (5% (w/v) imidazole and 55% (w/v) antipyrine (VWR AAA11089-36) cocktail at room temperature for 1.5 days. Cleared swollen brains were embedded in CUBIC-X2 with 2% agarose. This set of experiments and procedures were performed in accordance with standard ethical guidelines (European Communities Guidelines on the Care and Use of Laboratory Animals, 86/609/EEC) and were approved by the Cantonal Veterinary Office of the Canton of Basel-Stadt.

**Neurofilament staining of a whole-mount chicken embryo.**

The embryo was sacrificed at day 7 of development and incubated in 4% paraformaldehyde for 2 hours at room temperature. For best results, the embryo was kept in constant, gentle motion throughout the staining procedure. Incubation was at 4°C. The tissue was permeabilized in 1% Triton X-100/PBS for 15 hours, followed by an overnight incubation in 20 mM lysine in 0.1 M sodium phosphate, pH 7.3. Then the embryo was rinsed with five changes of PBS. Non-specific binding was blocked using 10% FCS (fetal calf serum) in PBS for 48 hours. The primary antibody mouse anti-neurofilament (1:1500, RMO270, Invitrogen 13-0700) was added for 60 hours. The primary antibody was removed and the tissue rinsed with ten changes of PBS and an additional incubation overnight. After re-blocking in FCS/PBS for 15 hours, the embryo was incubated with the secondary antibody goat anti-mouse IgG-Cy3 (1:500, Jackson ImmunoResearch 115-165-003) for 48 hours. In a next step, the embryo was washed ten times with PBS followed by incubation overnight in PBS. For imaging the tissue was dehydrated in a methanol gradient (25%, 50%, 75% in $H_2O$ and 2x 100%, 2 hours each step) and cleared using 1:2 benzyl alcohol: benzyl benzoate (BABB) solution overnight (again gentle shaking is recommended for dehydration and clearing). The tissue and staining are stable for months when kept at 4°C in the dark. This set of animal experiments and procedures were performed in accordance with standard ethical guidelines and were approved by the Cantonal Veterinary Office of the Canton of Zurich.

**BABB-clearing of Drosophila Melanogaster:**

We sacrificed a white-eyed *D. Melanogaster* after ether anesthesia and fixed the tissue in 4% PFA. Sample preparation followed the protocol by Dodt et al[9]. Briefly, after washing in PBS, the sample was dehydrated in an ethanol series (50%, 70%, 80%, 2x 100%) for 2h each and then transferred to the clearing solution (1:2 benzyl alcohol: benzyl benzoate, BABB). Imaging was performed in a 10 × 10 × 45 mm glass cuvette (filled with BABB) which was submerged in a 40 × 40 mm quartz immersion cuvette filled with BABB.

### iDISCO clearing and staining of mouse vasculature

For vasculature staining, brain tissue from 1 month old C57bl6/N males was fixed in 4% PFA without perfusion to preserve the endogenous immunoglobulins. Dye-conjugated secondary antibodies (1:100 dilution) against the tissue species were then used without need of primary antibodies. The protocol followed the one by Liebmann et al[70]. This experiment was approved by the Institutional Animal Care and Use Committee of ICM Brain and Spine Institute, Paris.

### iDISCO clearing and staining of rat samples

The brain from a female Wistar rat (age 16 weeks) was processed for enhanced autofluorescence and nuclear staining using iDISCO clearing protocol (adapted from Renier et al.[11]). The brain was fixed transcardially and postfixed overnight (4% PFA in PBS). After cryoprotection (30% sucrose in PBS), in order to enhance autofluorescence, the brain sample was additionally postfixed with 10% PFA in PBS for one week before clearing.

Clearing was performed following the protocol on http://idisco.info, with n=4 days. Although no immunolabeling was performed, the sample underwent all incubation steps indicated in the protocol, simply omitting antibodies in the solutions. On the beginning of day 4 of the "secondary Ab" incubation, TO-PRO (Thermo Fisher) was added at 1:2500 dilution. After 24 hours, the sample was washed twice with PTwH (30 min) to ensure removal of non-bound TO-PRO, and then the protocol resumed as specified on http://idisco.info.

### Human brain tissue preparation

Brain tissue samples were taken from one human body donor (no known neuropathological diseases) of the body donation program of the Department of Anatomy and Embryology, Maastricht University. The tissue donor gave informed and written consent to the donation of their body for teaching and research purposes as regulated by the Dutch law for the use of human remains for scientific research and education ("Wet op de Lijkbezorging"). Accordingly, a handwritten and signed codicil from the donor posed when still alive and well, is kept at the Department of Anatomy and Embryology Faculty of Health, Medicine and Life Sciences, Maastricht University, Maastricht, The Netherlands.

The brain was first fixed *in situ* by full body perfusion via the femoral artery. Under a pressure of 0.2 bar the body was perfused by 10 l fixation fluid (1.8 vol % formaldehyde, 20 % ethanol, 8.4 % glycerine in water) within 1.5-2 hours. Thereafter the body was preserved at least 4 weeks for post-fixation submersed in the same fluid. Subsequently, the brain was recovered by calvarian dissection and stored in 4% paraformaldehyde in 0.1 M phosphate buffered saline (PBS) for 25 months. All tissue was manually blocked with anatomical trimming blades, then cut into 5 mm thick slices in coronal orientation and immediately processed.

**Clearing and labelling of human brain tissue**

All human tissue samples were prepared following the MASH protocol[78]. As long as not indicated otherwise, samples were incubated in a volume of 5 ml in 6 well cell culture plates respectively. Samples were dehydrated in 20 %, 40 %, 60 %, 80 % and 100 % methanol (MeOH) for 1h each at room temperature (RT). After that samples were incubated for 1h in 100 % MeOH at $4^0$C and bleached in chilled, freshly prepared 5 % $H_2O_2$ in MeOH overnight at $4^0$C in the dark. Samples were then rehydrated 1h each in 80 %, 60 %, 40 %, 20 % MeOH and twice in phosphate buffered saline (PBS) containing 0.2 % Triton X-100 at RT. This was followed by another bleaching step in freshly filtered aqueous 50 % potassium metabisulfite solution and washing in distilled water for 1h each at RT. Staining was performed in 6 ml solution containing 0.001 % neutral red and a 1:7500 dilution of a 2 % methyl green stock solution[81] in PBS at pH 4 for 2.5 days. After that, the samples were turned on the other side, the solution was exchanged and incubation was continued for another 2.5 days. Then samples were washed 1h twice in PBS at pH 4, dehydrated once more in 20 %, 40 %, 60 %, 80 % and twice in 100 % MeOH. Samples were then transferred into 50 ml tubes and delipidated overnight in 66% dichloromethane (DCM)/33 % MeOH. Before immersion in ethyl cinnamate (ECi), sample were washed twice in 100 % DCM for 1h each. Samples were kept on a shaker during every step.

### Description of supplementary videos

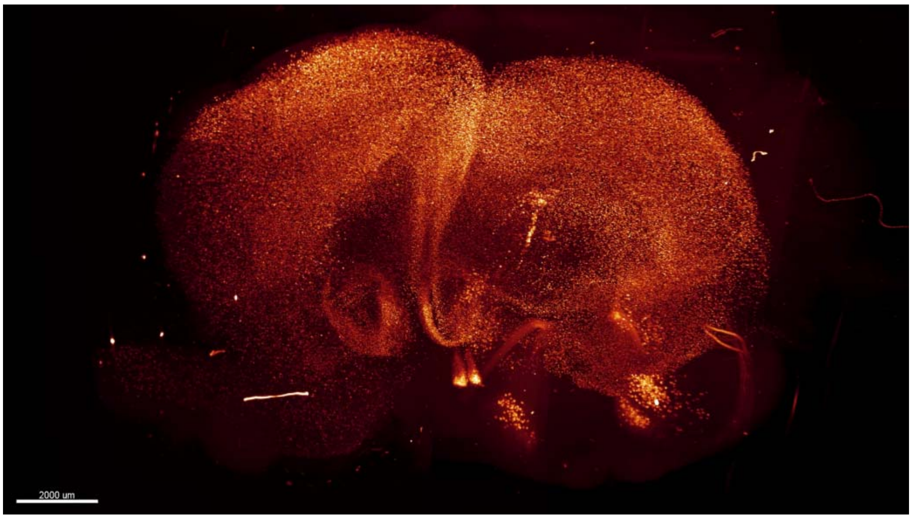

**Supplementary Video 1: Comparison of mesoSPIM imaging with and without ASLM in a VIPCre-tdTomato mouse brain.** A VIPCre-tdTomato mouse brain was processed using a passive CLARITY protocol and imaged with the mesoSPIM at 1× magnification. Without ASLM, only neurons along the midline are not blurred in the axial direction, whereas with ASLM, the mesoSPIM achieves quasi-isotropic imaging conditions.

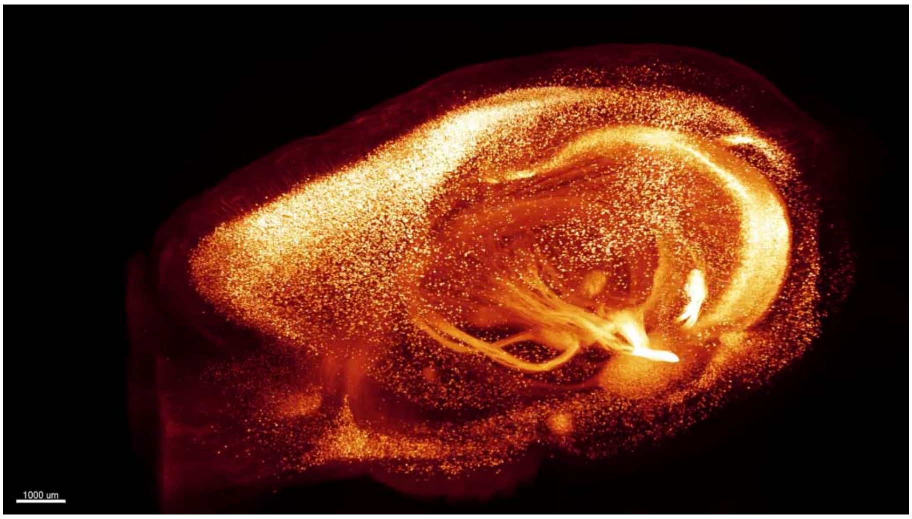

**Supplementary Video 2: Comparison of mesoSPIM imaging with and without ASLM in a Thy1-YFP mouse brain.** A VIP-tdTomato mouse brain was processed using an active CLARITY protocol and imaged with the mesoSPIM at 1× magnification. Without ASLM, only neurons along the midline are not blurred in the axial direction, whereas with ASLM, the mesoSPIM achieves quasi-isotropic imaging conditions.

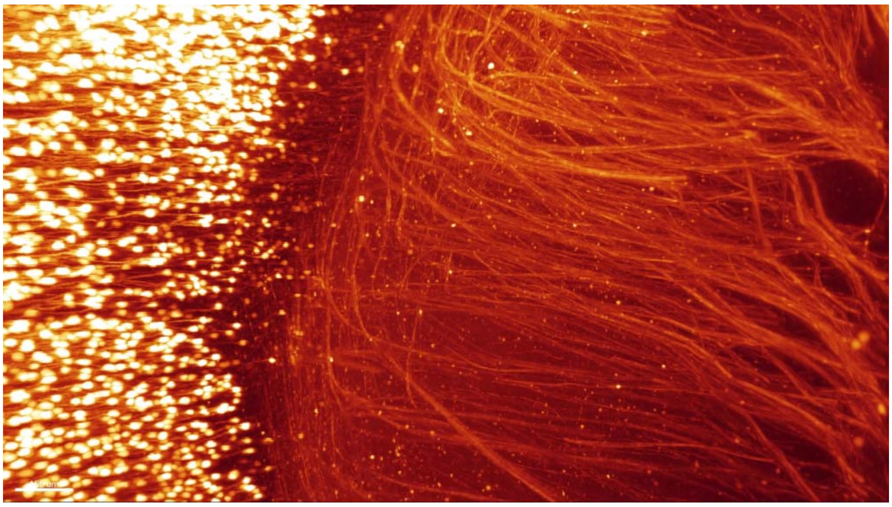

**Supplementary Video 3: Comparison of image quality with and without ASLM in a Thy1-YFP mouse brain at 4× magnification.** Without ASLM, the thickness inhomogeneity of the light-sheet leads to considerable blurring of features. After enabling the ASLM mode, axons become visible across the whole FOV.

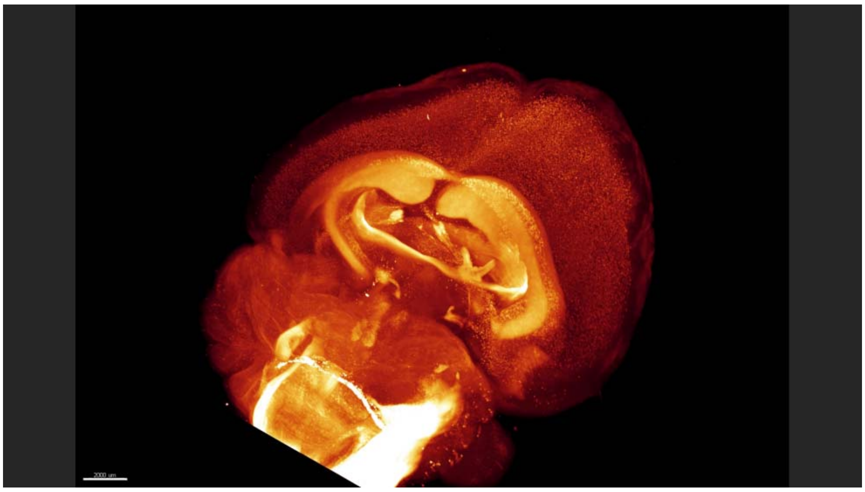

**Supplementary Video 4: Overview of a TPH2Cre-tdTomato mouse brain**. A TPH2Cre-tdTomato mouse brain was cleared using a passive CLARITY protocol and imaged with the mesoSPIM. Apart from expression in serotonergic nuclei, widespread labeling in the hippocampi and optic nerves is apparent.

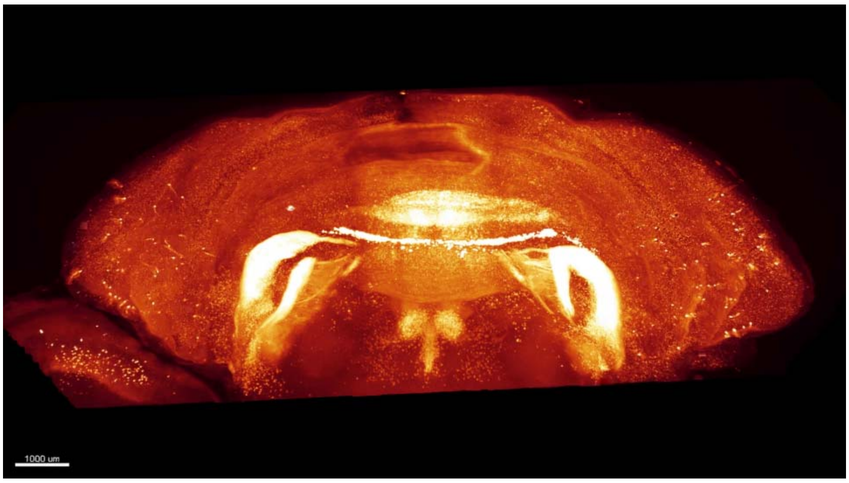

**Supplementary Video 5: Flythrough of the cerebellum of a TPH2Cre-tdTomato mouse brain**. Apart form widespread sparse expression in granule cells, a small subset of Purkinje neurons shows strong labeling. The dataset is stitched from 5 stacks taken at 4× magnification.

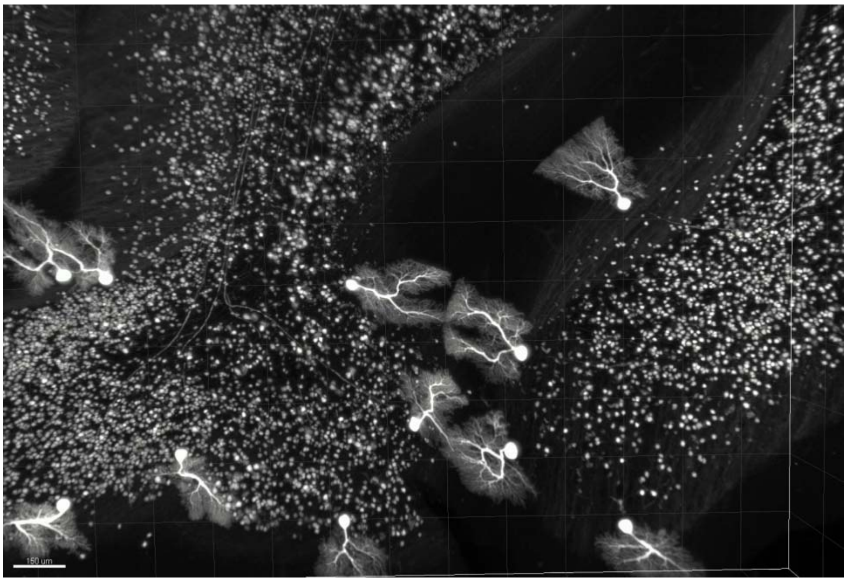

**Supplementary Video 6: Individual Purkinje cells in a TPH2Cre-tdTomato mouse brain**. Sparse expression leads to labeling of a subset of Purkinje neurons, including long-range axons to deep cerebellar nuclei.

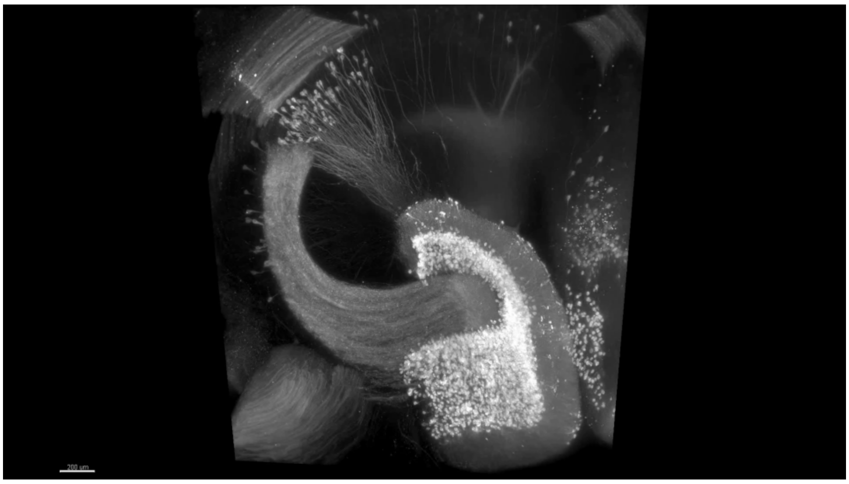

**Supplementary Video 7: Flythrough of the hippocampus of a  Rbp4Cre-YCX2.60 mouse**

A mouse brain expressing the calcium indicator YCX2.60 was cleared using a passive CLARITY protocol. Shown is a substack taken at 4× magnification in the hippocampus. Major structures such as the dentate gyrus, mossy fiber pathway and sparsely expressing neurons in CA2 are visible.

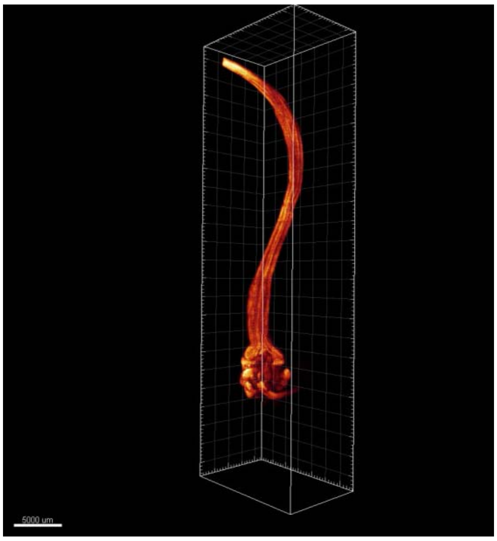

**Supplementary Video 8: Whole-CNS imaging with the mesoSPIM.** A whole CNS was dissected from a  GlyT2-EGFP-mouse and processed using the X-CLARITY protocol. The resuting dataset is stitched from individual stacks at 1× and 4× magnification and shows individual glycinergic neurons in the hindbrain and spinal cord.

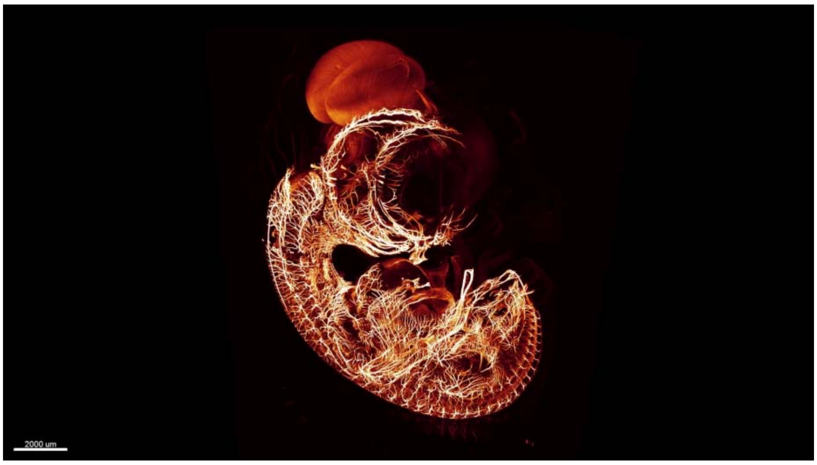

**Supplementary Video 9: Flythrough of a BABB-cleared chicken embryo.** A 7-day old chicken embryo (stage HH31) was stained for neurofilament and cleared using BABB. The resulting dataset shows the entire developing nervous system.

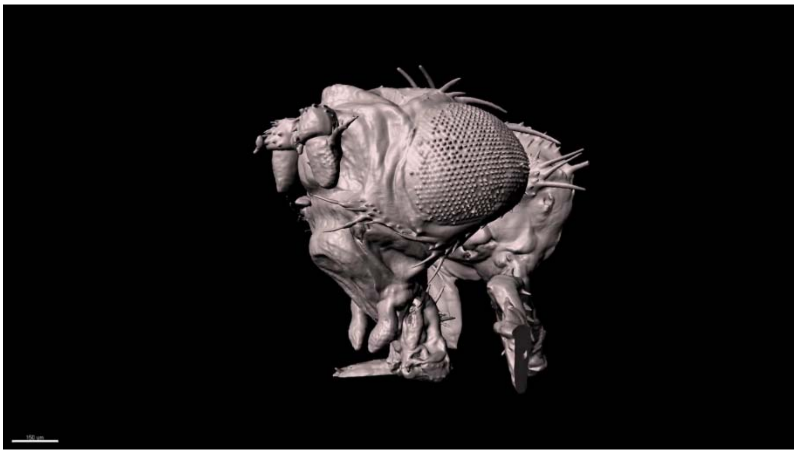

**Supplementary Video 10: Imaging of a BABB-cleared D. melanogaster.** A white eyed fly was cleared using the BABB protocol. In both single sections and surface renderings, the mesoSPIM allows visualization of large anatomical structures of interest.

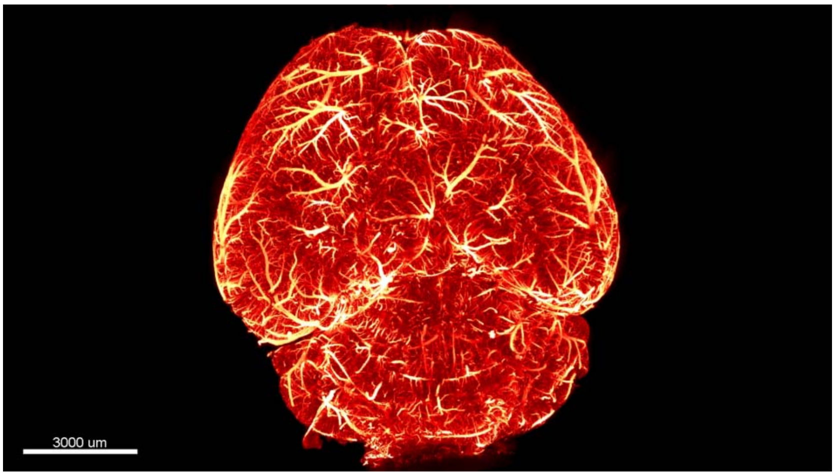

**Supplementary Video 11: Flythrough of a iDISCO mouse brain.** Vasculature in a iDISCO-cleared mouse brain was labeled using anti-IgG antibodies. Throughout the whole sample, single capillaries are visible.

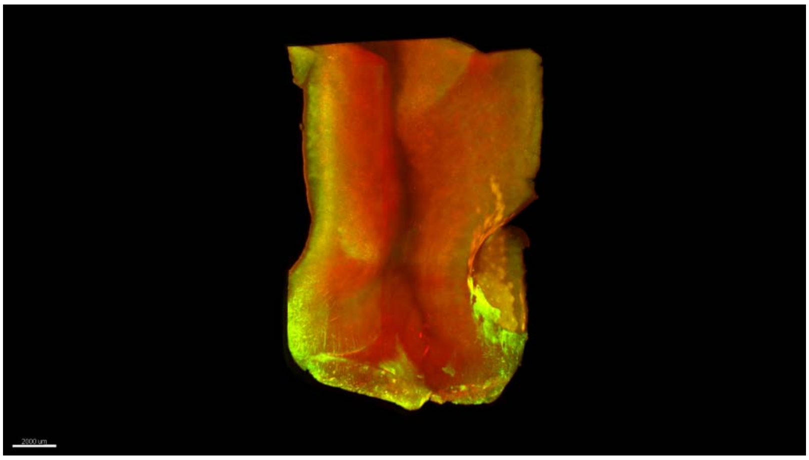

**Supplementary Video 12: Overview of an MASH (iDISCO/ECi)-processed human cortex sample.** A slab of human cortex was stained using the MASH protocol and then cleared using ECi. At low magnification (1×), the mesoSPIM allows visualization of cortical gyri and sulci.

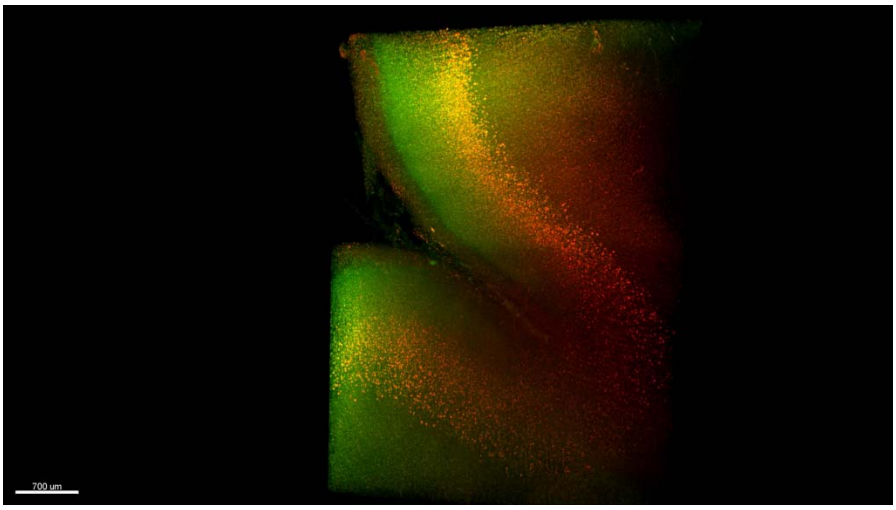

**Supplementary Video 13: Flythrough of an MASH (iDISCO/ECi)-processed human cortex sample.** A slab of human cortex was stained using the MASH protocol and then cleared using ECi. At high magnification (4×), the mesoSPIM allows visualization of individual neurons in a sulcus.
